# Genome-wide association studies reveal the complex genetic architecture of DMI fungicide resistance in *Cercospora beticola*

**DOI:** 10.1101/2020.11.12.379818

**Authors:** Rebecca Spanner, Demetris Taliadoros, Jonathan Richards, Viviana Rivera-Varas, Jonathan Neubauer, Mari Natwick, Olivia Hamilton, Niloofar Vaghefi, Sarah Pethybridge, Gary A. Secor, Timothy L. Friesen, Eva H. Stukenbrock, Melvin D. Bolton

## Abstract

Cercospora leaf spot is the most important disease of sugar beet worldwide. The disease is caused by the fungus *Cercospora beticola* and is managed principally by timely application of fungicides including those of the sterol demethylation inhibitor (DMI) class. However, reliance on DMIs has caused an increase in resistance to this class of fungicides in multiple *C. beticola* populations. To better understand the genetic and evolutionary basis for resistance in *C. beticola*, a genome-wide association study (GWAS) and selective sweep analysis were conducted for the first time in this fungal plant pathogen. We performed whole genome resequencing of 190 *C. beticola* isolates predominantly from North Dakota and Minnesota that were phenotyped for sensitivity to tetraconazole, the most widely used DMI fungicide in this region. GWAS identified mutations in genes associated with DMI fungicide resistance including a Regulator of G-protein Signaling (RGS) protein, an ATP-binding cassette (ABC) pleiotropic drug resistance transporter, a dual-specificity tyrosine phosphorylation-regulated kinase (DYRK), and a gene annotated as a hypothetical protein. A SNP upstream of *CbCYP51*, the gene encoding the target of DMI fungicides, was also identified via GWAS. Haplotype analysis of CbCYP51 identified a synonymous mutation (E170) in high linkage disequilibrium with the upstream SNP, and multiple non-synonymous mutations (L144F, I387M and Y464S) associated with DMI resistance. Additionally, a putative codon bias effect for the L144F substitution was identified that generated different resistance potentials. We also identified a CbCYP51 paralog in *C. beticola*, CbCYP51-like, with high protein homology to CYP51C found uniquely in *Fusarium* species but *CbCYP51-like* does not appear to influence DMI sensitivity. Genome-wide scans of selection showed that several of the GWAS mutations for fungicide resistance resided in regions that have recently undergone a selective sweep. Using radial plate growth on selected media as a fitness proxy, we did not find a trade-off associated with DMI fungicide resistance suggesting that resistance mutations can persist in *C. beticola* populations. Taken together, we show that population genomic data from a crop pathogen can allow the identification of mutations conferring fungicide resistance and inform about their origins in the pathogen population.

## Introduction

Demethylation inhibitor (DMI) compounds are effective antifungals in both medicine and agriculture for managing a broad range of fungal pathogens (Becher and Wirsel 2012). The DMIs, or azoles, inhibit fungal growth by interfering with sterol 14α-demethylase (Vanden Bossche et al. 1987), also known as cytochrome P450 monooxygenase family 51 (CYP51). Fungal CYP51 is required for synthesis of ergosterol, a key sterol component of fungal cell membranes required to maintain permeability and fluidity (Daum et al. 1998). DMIs have shown unique durability when compared to other single site fungicides, with control failures being rare even with widespread and prolonged use (Cools et al. 2013). However, resistance has still emerged in some fungal populations with long term exposure to DMIs, leading to reduced efficacy of the compounds in use (Staub 1991).

DMI resistance is often associated with changes to the molecular target CYP51 (Becher and Wirsel 2012). Amino acid substitutions in CYP51 (Kelly et al. 1999a; Kelly et al. 1999b; Lamb et al. 2000; Snelders et al. 2011) or overexpression of *CYP51* (Carter et al. 2014; Ghosoph et al. 2007; Hamamoto et al. 2000; Ma et al. 2006; Villani et al. 2016) can lead to decreased DMI sensitivity. Some filamentous fungi have two or more paralogous *CYP51* genes (Chen et al. 2020; Hawkins et al. 2014; Liu et al. 2011) which may result in an inherent reduction in DMI sensitivity and allow these species to overcome some biological costs by restricting acquired resistance to one paralog (Becher and Wirsel 2012; Cools et al. 2013). Gain-of-function mutations in transcription factors (Dunkel et al. 2008; Liu et al. 2015) regulating ergosterol biosynthesis genes have also been linked to reduced DMI efficacy.

Non-CYP51 mechanisms of resistance can also be important in fungi. Such mechanisms include enhanced efflux of DMIs (Hahn and Leroch 2015) by plasma membrane-bound transporters in the multi-facilitator (MFS) or ATP-binding cassette (ABC) superfamilies (de Ramón-Carbonell et al. 2019; Hayashi et al. 2002a, b; Hellin et al. 2018; Leroux and Walker 2011; Zwiers et al. 2002), calcium signaling regulators (Edlind et al. 2002; Jain et al. 2003; Jia et al. 2009; Jia et al. 2012; Li et al. 2019; Zhang et al. 2013), the pleiotropic effect of melanization (Lendenmann et al. 2015) and other uncharacterized genes (Ballard et al. 2019).

Genetic variants that confer reduced susceptibility to DMI fungicides may also have a fitness penalty (Hawkins and Fraaije 2018). However, if the benefits of the resistance mutation outweigh the costs, it will increase in frequency in a fungal population that is frequently exposed to DMI fungicides (Milgroom 1989). Signatures of positive selection have previously been detected for variants of *CYP51* in *Zymoseptoria tritici* (Brunner et al. 2016) and ABC transporter genes *CDR1* and *CDR2* in *Candida albicans* (Holmes et al. 2008). Positively selected mutations leave a distinct signature in the genome termed a ‘selective sweep’. A selective sweep is characterized by a locus deprived of genetic variation and high linkage disequilibrium in the genomic regions flanking the favorable mutation. This pattern reflects the “hitchhiking” of genetic variants linked to the beneficial mutation which also increase in frequency (Smith and Haigh 1974). The identification of fungicide resistance loci in selective sweep regions suggests fungicides are a major selective pressure in recent evolution of a fungal pathogen (Hartmann et al. 2020).

DMI fungicides are integral for managing many important crop diseases (Price et al. 2015), including Cercospora leaf spot (CLS) disease of sugar beet (*Beta vulgaris* spp. *vulgaris*) CLS remains the most destructive foliar disease of sugar beet worldwide (Rangel et al. 2020). In addition to adequate host genetic tolerance and cultural practices to suppress the causal pathogen, timely fungicide applications are critical for disease management (Secor et al. 2010). The Red River Valley (RRV) region of North Dakota and Minnesota, USA is the largest sugar beet production area in the United States (NASS 2020) and has historically experienced huge economic losses due to CLS with large reductions in yield and the application of non-efficacious fungicides (Bolton et al. 2012b; Secor et al. 2010). The use of DMI fungicides in the RRV region began in 1999 when the Environmental Protection Agency granted an emergency exemption for sugar beet growers to use tetraconazole to manage CLS (Secor et al. 2010). Management of fungicide resistance in the RRV has become a cooperative effort to maintain efficacy of fungicide groups (Khan and Smith 2005; Secor et al. 2010). This predominantly involves mixing and rotating fungicide chemistries within and between sprays, coupled with annual assessment of fungicide sensitivities in *C. beticola*.

The magnitude of DMI resistance (measured by EC_50_ values) in *C. beticola* and incidence of resistant isolates (EC_50_ > 1 μg/mL) in RRV field populations has steadily increased since 2006 (Rangel et al. 2020). Factors contributing to the rapid development of fungicide resistance in *C. beticola* are its polycyclic nature, high rate of sporulation and common spray programs used over large areas for disease management (Dekker 1986). Although experimental crosses of *C. beticola* are not possible to our knowledge, there is indirect population genetic evidence of sexual reproduction (Bolton et al. 2014; Bolton et al. 2012c; Groenewald et al. 2006b; Groenewald et al. 2008). In contrast, *C. beticola* populations from table beet exhibit microsatellites in linkage disequilibrium and do not always have equal mating type ratios, suggesting a mixed mode of reproduction (Knight et al. 2018; Knight et al. 2019; Vaghefi et al. 2016; Vaghefi et al. 2017).

Evaluating levels of resistance is an important part of CLS fungicide resistance management (Secor et al. 2010) and has been aided by the development of PCR-based mutation detection tools to expedite the process (Birla et al. 2012; Bolton et al. 2012a; Shrestha et al. 2020). These tools have greatly increased the ability to effectively identify fungicide-resistant strains, which enables growers to make educated fungicide application decisions based on the resistance profiles of their fields. However, molecular methods of resistance detection first require the identification of associated mutations. In *C. beticola*, overexpression of *CbCYP51* has been associated with high levels of DMI resistance in isolates from Greece (Nikou et al. 2009) and the US (Bolton et al. 2012a; Bolton et al. 2016). However, no upstream insertions, duplications or transposable elements have been found that could be associated with gene overexpression (Bolton et al. 2012a; Nikou et al. 2009). Nikou et al. (2009) found that a single synonymous mutation at position E170 was associated with *CbCYP51* overexpression and DMI resistance. However, Obuya et al. (2015) could not demonstrate its involvement in DMI resistance through heterologous expression in yeast. Amino acid substitutions L144F, I387M and Y464S in CbCYP51 appeared to coincide with DMI resistance in *C. beticola* isolates from Serbia (Trkulja et al. 2017) but overall, it has been difficult to clearly associate any *CbCYP51* haplotype with resistance (Bolton et al. 2012a; Trkulja et al. 2017). Factors including small sample size may play a significant role in the interpretation of results in such studies.

Genome-wide association study (GWAS) analysis is a powerful method for identifying genetic variants associated with complex traits such as DMI fungicide resistance (Sanglard 2019). This method can be performed using natural variation within a population and is a good alternative to biparental mapping when sexual crosses are not possible or convenient. The advent of cost-effective high-throughput sequencing technology is enabling whole genome sequencing for GWAS in a broad range of organisms (Power et al. 2017). Fungi are particularly amenable to GWAS due to their small genomes, haploidy and high rates of recombination (Falush 2016). GWAS has been successfully employed to identify loci associated with DMI resistance in several phytopathogenic fungi (Mohd-Assaad et al. 2016; Pereira et al. 2020; Talas et al. 2016). We hypothesized that GWAS would be an ideal strategy for finding genetic determinants underlying DMI resistance in *C. beticola*, a pathogen that cannot be experimentally crossed yet but shows considerable genetic variation (Bolton et al. 2012c; Groenewald et al. 2006a; Groenewald et al. 2008; Moretti et al. 2006; Moretti et al. 2004; Rangel et al. 2020; Vaghefi et al. 2016). In this study, we revealed the genetic architecture of DMI fungicide resistance in *C. beticola* by performing an unbiased GWAS in 190 *C. beticola* isolates. Thirteen loci across five genes were associated with resistance to tetraconazole. Ten of these loci overlapped with genome-wide selective sweep regions, revealing that they were recently selected in this population. Using radial plate growth assays as a fitness proxy, we did not identify a fitness penalty for tetraconazole resistance *in vitro*. Finally, we utilized a Cas9-RNP gene editing method to functionally assess potential fungicide resistance mutations.

## Results

### Field sampling of *C. beticola* isolates and selection for genome sequencing

We initially set out to investigate the genetic architecture of DMI fungicide resistance in *C. beticola* isolates collected from two adjacent fields near Fargo, North Dakota, USA in 2016. Since previous studies have demonstrated that *C. beticola* populations have substantial genetic variation (Groenewald et al. 2007; Moretti et al. 2006; Moretti et al. 2004), even exhibiting multiple genotypes on a single leaf (Bolton et al. 2012c), we reasoned that isolates collected from two fields would harbor enough genetic variation for GWAS. Approximately 300 isolates were clone-corrected using eight microsatellite markers and unique isolates were subsequently phenotyped for tetraconazole sensitivity by calculating EC_50_ values. From these isolates, a population of 62 unique *C. beticola* isolates was selected based on tetraconazole EC_50_: approximately half (34 isolates) had a tetraconazole EC_50_ of below 1.0 μg/mL (arbitrarily considered sensitive) (Bolton et al. 2012b) and the other half (28 isolates) had an EC_50_ value of 1.0 μg/mL or higher and were considered resistant (Table S1). These 62 isolates underwent Illumina whole genome resequencing with an aim of 25X or greater genome coverage. After filtering for genotype quality and read depth, 672,149 genetic variants (SNPs and indels) were identified. All GWAS attempts failed to produce models that corrected for population structure adequately and thus generated either highly overinflated or underinflated *P*-values (data not shown). To overcome this, we sequenced an additional 128 isolates that were collected from different infected commercial fields throughout the Red River Valley (RRV) region of Minnesota and North Dakota, except two isolates collected from Idaho, during field surveys in 2016 (n=80) and 2017 (n=48) (Table S1). These isolates were selected due to their more diverse geographic origins within the RRV and more extreme tetraconazole sensitivities. In total, 190 isolates of *C. beticola* were used in this study (Table S1).

EC_50_ values were calculated for all 190 isolates to tetraconazole, the active ingredient of Eminent^®^ fungicide which is widely used in the RRV region (Fig. S1A). The highest EC_50_ value measured was the maximum concentration used of 100 μg/mL indicating that the true value could be greater (Table S1). The lowest EC_50_ obtained was 0.008 μg/mL, comparable to EC_50_ values of *C. beticola* isolates collected in the RRV region in 1997-98 (Bolton et al. 2012b).

### Genome sequencing of *C. beticola* isolates

To map the genetic architecture of resistance to DMI fungicides, we performed whole genome resequencing of 190 *C. beticola* isolates. We mapped Illumina reads of each isolate to the 09-40 reference genome (de Jonge et al. 2018) (NCBI RefSeq assembly GCF_002742065.1). The resulting coverage per genome ranged from 18X to 40X with a mean coverage of 32X (Table S2). GATK HaplotypeCaller was used to call SNPs and indels against the reference genome for each isolate. We used the default diploid ploidy level, instead of −ploidy 1 option in our haploid fungus, to allow us to filter out variants in any poorly aligned regions that resulted in heterozygous calls. After filtering for genotype quality and read depth, 868,218 variants were identified including 732,852 SNPs, corresponding to an average SNP density of ~20 SNPs per kb. All heterozygous sites were transformed to missing data. For genome-wide association mapping, the identified SNPs/indels were filtered further. A minor allele frequency of 0.05 reduced variants to 424,456, eliminating over half of called variants. Filtering for 10% missing data reduced the total number to 320,530 variants.

Mapping power of GWAS was assessed by calculating linkage disequilibrium (LD) decay for the population. LD decayed to r^2^ < 0.2 rapidly within ~3.5 kb (Fig. S2), which is comparable to values found in populations of other closely-related filamentous fungal phytopathogens used successfully for GWAS such as *Zymoseptoria tritici* (Hartmann et al. 2017) and *Parastagonospora nodorum* (Gao et al. 2016; Pereira et al. 2020; Richards et al. 2019).

### Population structure analyses

We performed a principle component analysis (PCA) to assess population structure amongst the 190 *C. beticola* isolates. PC1 explained 11% of total variation followed by 3.4% and 3.0% for PCs 2 and 3, respectively. Analysis of population structure often reveals clustering of individuals according to geographical origin. However, pairwise plots of the first six PCs from PCA demonstrated that sampling location had little impact on clustering of the *C. beticola* isolates used in this study (Figs. 1A and S3). Intriguingly, the tight cluster of 66 isolates circled in Figures 1A and 1B were predominantly tetraconazole-sensitive (28 isolates are moderately sensitive, 34 isolates are sensitive), whereas the remaining scattered isolates were mainly tetraconazole-resistant. Some clustering of sensitive isolates was also visible in additional pairwise plots of the first six PCs from PCA (Fig. S4). This illustrates how fungicide use is shaping the population structure of *C. beticola* in RRV. Based on this observation, we hypothesized that certain regions in the genome encoding traits involved in fungicide resistance may explain more of the variation in the population. Therefore, we also conducted a PCA for each chromosome separately. Indeed, chromosome-specific PCAs revealed that chromosome nine had the highest proportion of variation explained by PC1 at 13% and had the strongest clustering of strains according to tetraconazole sensitivity in pairwise plots of the first two PCs (Fig. S5).

**Figure 1.**
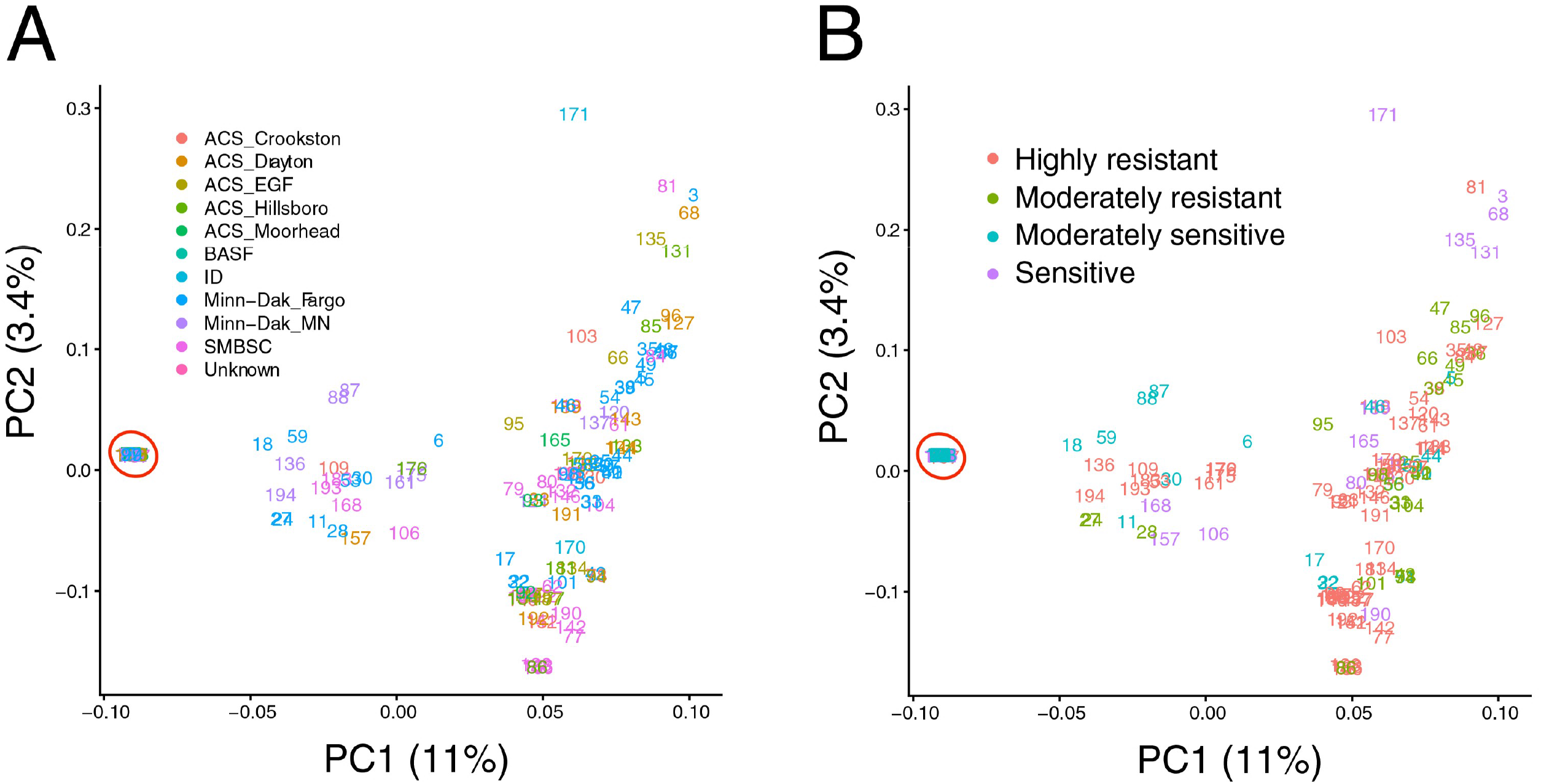
Principle component analyses. The first two principle components plotted from a PCA of *Cercospora beticola* isolates performed with 37,973 LD-pruned genome-wide SNPs. Plots use the same data but are color-coded by A) field sampling location and B) tetraconazole sensitivity. The cluster of strains circled in red is comprised of 66 isolates, 62 of which are either moderately sensitive or sensitive to tetraconazole. Highly resistant = isolates with EC_50_ ≥ 10 μg/mL; Moderately resistant = isolates 1 μg/mL ≤ EC_50_ < 10 μg/mL; Moderately sensitive = isolates with 0.1 μg/mL ≤ EC_50_ < 1 μg/mL; Sensitive = isolates with EC_50_ < 0.1 μg/mL.

### Genetic architecture of tetraconazole sensitivity

To determine the genetic architecture of tetraconazole sensitivity in *C. beticola*, we performed GWAS using all 190 isolates. Model selection was based on visualization of Q-Q plots to ensure correction for population structure, as well as the use of GAPIT Bayesian Information Criterion to determine the optimal number of principle components (zero in this case). We also looked for the most significantly associated markers to consistently appear throughout multiple models. The MLM model chosen yielded 13 significant associations at a significance threshold of −log_10_ (*P*-value) = 4.5 (Fig. 2, Table S3, Fig. S6). Of these associated markers, ten were intragenic SNPs, two were indels within 5’UTRs and one was within 124 bp of the nearest gene.

**Figure 2.**
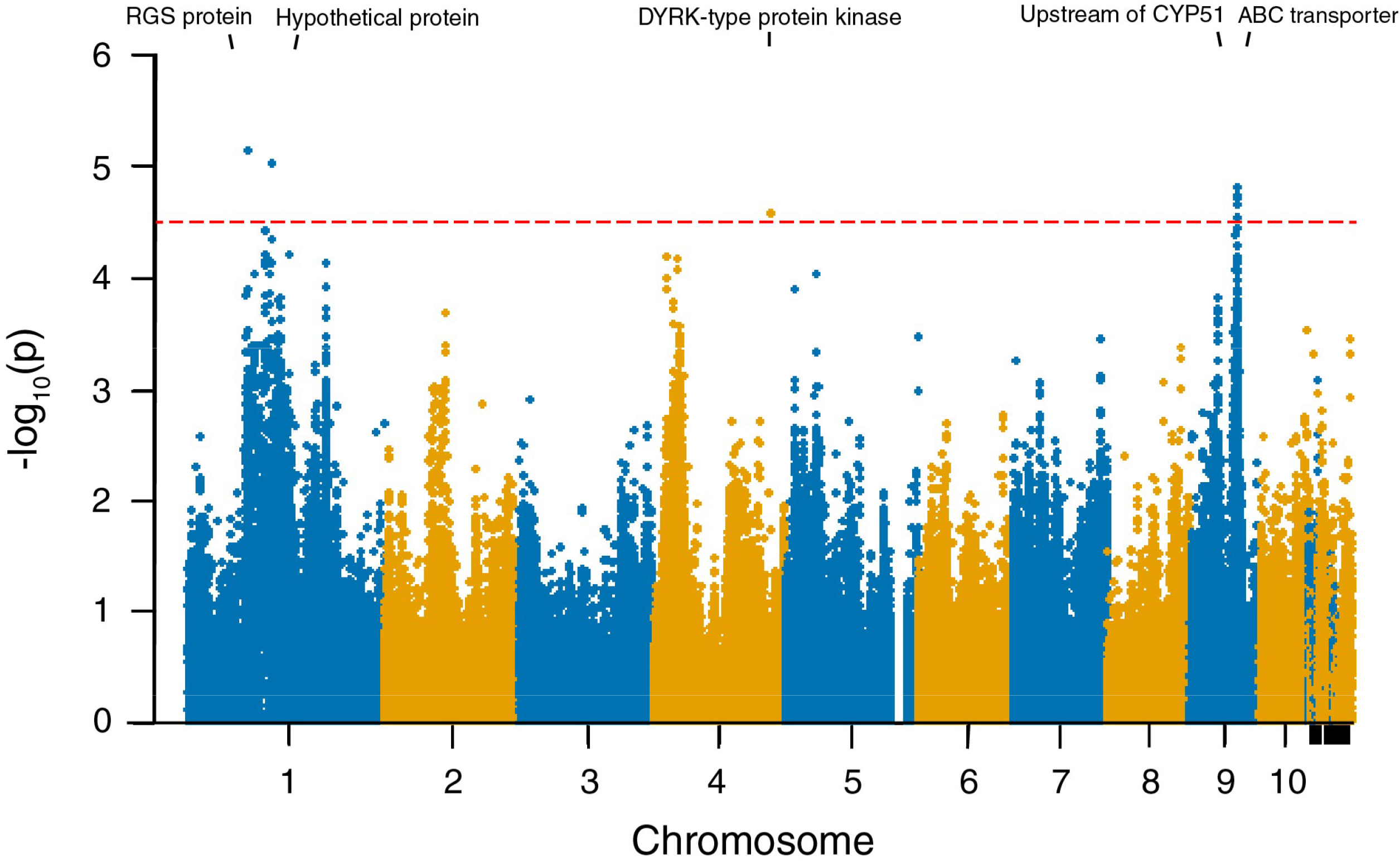
GWAS of tetraconazole sensitivity in *Cercospora beticola*. Manhattan plot displaying marker associations with tetraconazole EC_50_ values. The red line represents the genome-wide significance threshold of −log_10_(*P*)=4.5. The genomic position of genes with significantly associated markers are indicated above the plot.

### Genes associated with tetraconazole sensitivity

The potential effects of significantly associated markers on tetraconazole sensitivity were investigated. Since we also identified amino acid substitutions in CbCYP51 associated with tetraconazole sensitivity (discussed below), we compared the effects of GWAS markers on tetraconazole sensitivity between strains harboring and lacking CbCYP51 mutations.

#### RGS domain protein

The most significantly associated marker was 1_1890198, which was within a 76 bp indel in the 5’UTR of gene CB0940_00689 encoding a protein with a regulator of G-protein signaling (RGS) domain. The closest flanking markers were ~2 kb away and in relatively low LD (R^2^ < 0.5) with the significantly associated 1_1890198 indel (Fig. S7). The reference genome harbored the insertion (spanning 1,890,172-1,890,247 bp) that was absent in the alternate allele. The 5’UTR insertion was associated with a significant increase in tetraconazole EC_50_ value (*P* < 0.01) in the absence of CbCYP51 amino acid substitutions (Fig. 3A). The predicted 1,232 amino acid sequence had multiple conserved domains including the aforementioned RGS domain, a PXA domain, a flagellar switch protein FliM domain, the phosphoinositide binding Phox homology (PX) domain of yeast Mdm1 and a C-terminus sorting nexin domain.

**Figure 3.**
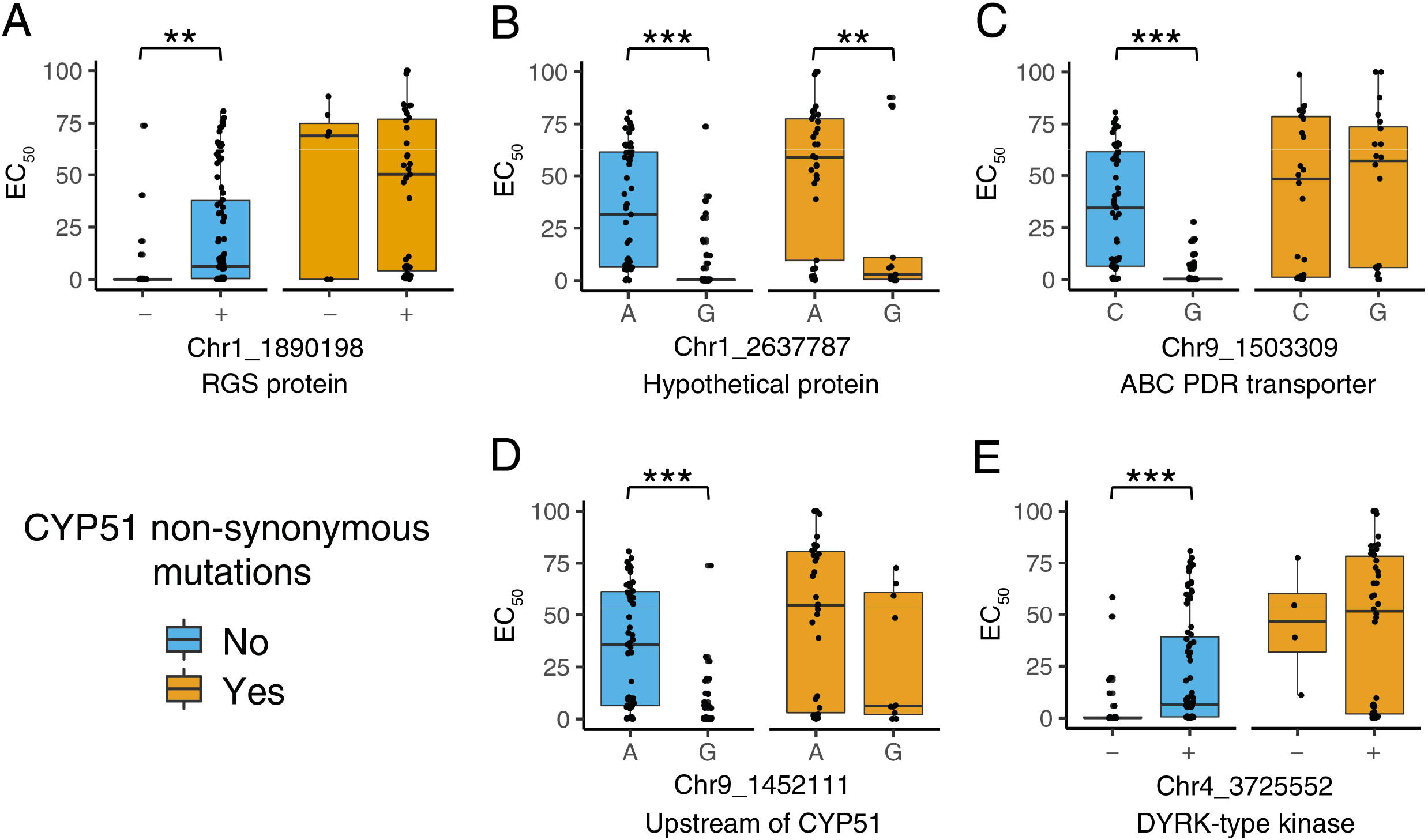
Effects of significantly associated markers on tetraconazole sensitivity. The effects of DMI fungicide resistance loci A) indel 1_1890198 in the 5’UTR of a RGS protein gene, B) SNP 1_2637787 within a hypothetical protein gene, C) SNP 9_1503309 within an ABC PDR transporter gene, D) SNP 9_1452111 upstream of *CbCYP51* and E) indel 4_3725552 in the 5’UTR of a DYRK gene, both with and without *CbCYP51* amino acid substitutions present. Significant differences in EC_50_ between alleles are displayed at *P* < 0.05 (*), *P* < 0.01 (**) and *P* < 0.001 (***).

#### Hypothetical protein

The second most significantly associated marker was the SNP 1_2637787 within hypothetical protein gene CB0940_00999 that lacked domains of known function. Whilst there were 134 other markers ±3 kb, the 1_2637787 SNP was in low LD with most of these (R^2^ ≈ 0) (Fig. S8). The SNP conferred an amino acid substitution from glycine to serine at codon 474 (G474S) in the hypothetical protein of 488 amino acids. The serine substitution gave rise to a significant increase in tetraconazole EC_50_ value (*P* < 0.001), even in the presence of *CbCYP51* non-synonymous mutations (*P* < 0.01) (Fig. 3B).

#### ABC PDR transporter

The third most significant association was SNP 9_1503309 within the ATP-binding cassette (ABC) transporter gene CB0940_11399. This mutation conferred a synonymous substitution at codon 191 (L191). The C allele was associated with a significant increase in tetraconazole EC_50_ when compared to the G allele (*P* < 0.001) in the absence of CbCYP51 amino acid substitutions (Fig. 3C). Eight additional SNPs within this gene were significantly associated with tetraconazole sensitivity, five of these were synonymous mutations (at codons G187, S189, T190, K193 and L195) in the first exon and three were within the second intron (Table S3). This protein sequence is comprised of 1,493 amino acids with a highly conserved ABC-2 transporter domain and pleiotropic drug resistance (PDR) sub-family nucleotide-binding domain. The majority of gene polymorphisms (98) were found within the first three exons and two introns and most were in perfect LD (R^2^ = 1) with 9_1503309 (Fig. S9). Of these markers, 76 were annotated as synonymous mutations, seven as intronic, three as splice site variants, nine as missense mutations and three as frameshift variants.

#### Upstream of CbCYP51

The fourth most significant association was SNP 9_1452111 and lies ~50 kb upstream of the ABC transporter gene on chromosome nine. This SNP was just 124 bp upstream of the start codon of *CbCYP51*, encoding the DMI fungicide target. When A was present instead of G, there was a significant increase in tetraconazole EC_50_ value (*P* < 0.001) in the absence of CbCYP51 amino acid substitutions (Fig. 3D). The 9_1452111 SNP was in low LD with the two flanking markers (R^2^ < 0.2) but was in higher LD with the intragenic *CbCYP51* SNP 9_1451478 (R^2^ = 0.7) and a block of markers upstream of the gene (R^2^ > 0.5) (Fig. S10).

#### Dual-specificity tyrosine phosphorylation-regulated protein kinase (DYRK)

The fifth most significantly associated locus was marker 4_3725552. This was an indel within the 5’UTR of a putative dual-specificity tyrosine phosphorylation-regulated protein kinase (DYRK) (gene CB0940_05141). Although there were 116 other markers ±3 kb of 4_3725552, it was in relatively low LD with each of these (R^2^ < 0.5) and had a much higher GWAS *P*-value (Fig. S11). The 61 bp insertion in the reference allele was associated with an increase in tetraconazole EC_50_ value (*P* < 0.001) in the absence of CbCYP51 amino acid substitutions (Fig. 3E). This 263-amino acid protein had a highly conserved DYRK domain that enables the kinase to phosphorylate serine/threonine residues in other proteins and autophosphorylate via tyrosine residues (Aranda et al. 2011).

### Synonymous and non-synonymous mutations in CbCYP51 are associated with DMI fungicide resistance

Genome-wide association analyses of tetraconazole sensitivity suggested the involvement of the *CbCYP51* locus, with a significantly associated SNP (9_1452111) 124 bp upstream of the *CbCYP51* start codon (Fig. 2). In *C. beticola, CbCYP51* is a single copy intron-free gene of 1,632 bp (NCBI XP_023450255.1, CB0940_11379) (Bolton et al. 2012a; Nikou et al. 2009). Since intrinsic *CbCYP51* expression was previously shown to be higher in some resistant *C. beticola* strains (Bolton et al. 2012a; Bolton et al. 2016; Nikou et al. 2009), we questioned whether mutations upstream of *CbCYP51* may affect *CbCYP51* expression and fungicide resistance. No insertions or retrotransposons were found immediately upstream of *CbCYP51* although several SNPs were identified within 3 kb either side of the gene (Fig. S10). Variation in read depth was not observed at the *CbCYP51* locus, suggesting no variation in copy number. SNP 9_1452111 was located 124 bp upstream of the *CbCYP51* locus and was notably in high LD with 9_1451478, a silent mutation at E170 (R^2^ = 0.7) (Fig. S10), thus exhibiting similar effects on EC_50_ value. All isolates with E170 also had the upstream SNP. Eighteen isolates harbored the upstream SNP without E170, but these also had non-synonymous changes in CbCYP51 (not shown). Therefore, we cannot evaluate the individual effects of the upstream SNP and E170 on tetraconazole EC_50_ value without functional studies.

The putative effects of the 9_1452111 SNP upstream of *CbCYP51* on transcription factor binding sites were predicted using the JASPAR database (Fornes et al. 2020) (Table S4). Query sequences of 21 bp revealed differences in putative binding sites (all defined experimentally in *Saccharomyces cerevisiae*) for both versions of the SNP (Table S4). UPC2 and STE12 bindings sites were present for both versions, but the resistant allele (T) had a stronger UPC2 binding site predicted on the negative strand and a weaker STE12 binding site predicted on the positive strand. The resistant allele (T) also had GLN3 and SPT23 binding sites predicted that were absent for the sensitive allele (C). The sensitive allele (C) had ECM22 and STB5 bindings sites uniquely predicted.

We also investigated the presence of target site mutations in *CbCYP51* that may influence DMI sensitivity. We found 11 different *CbCYP51* gene coding sequences (haplotypes) in our set of 190 RRV region isolates. There were three different “DMI-sensitive” haplotypes; the most common harbored by 85 isolates (haplotype 3), a highly divergent haplotype (haplotype 1) found in three isolates that harbored 113 SNPs and an 82 bp deletion and a third (haplotype 2) represented by a single sensitive isolate harboring a silent mutation at I122 and the amino acid substitution V467A (Fig. 4). When comparing the remaining haplotypes to the most common sensitive haplotype (#3), there were five different non-synonymous mutations (L144F, I309T, I387M, Y464S and V467A; Fig. 4). The presence of amino acid substitutions L144F, I387M or Y464S gave higher tetraconazole EC_50_ values when compared to sensitive haplotype #2 (Fig. 4).

**Figure 4.**
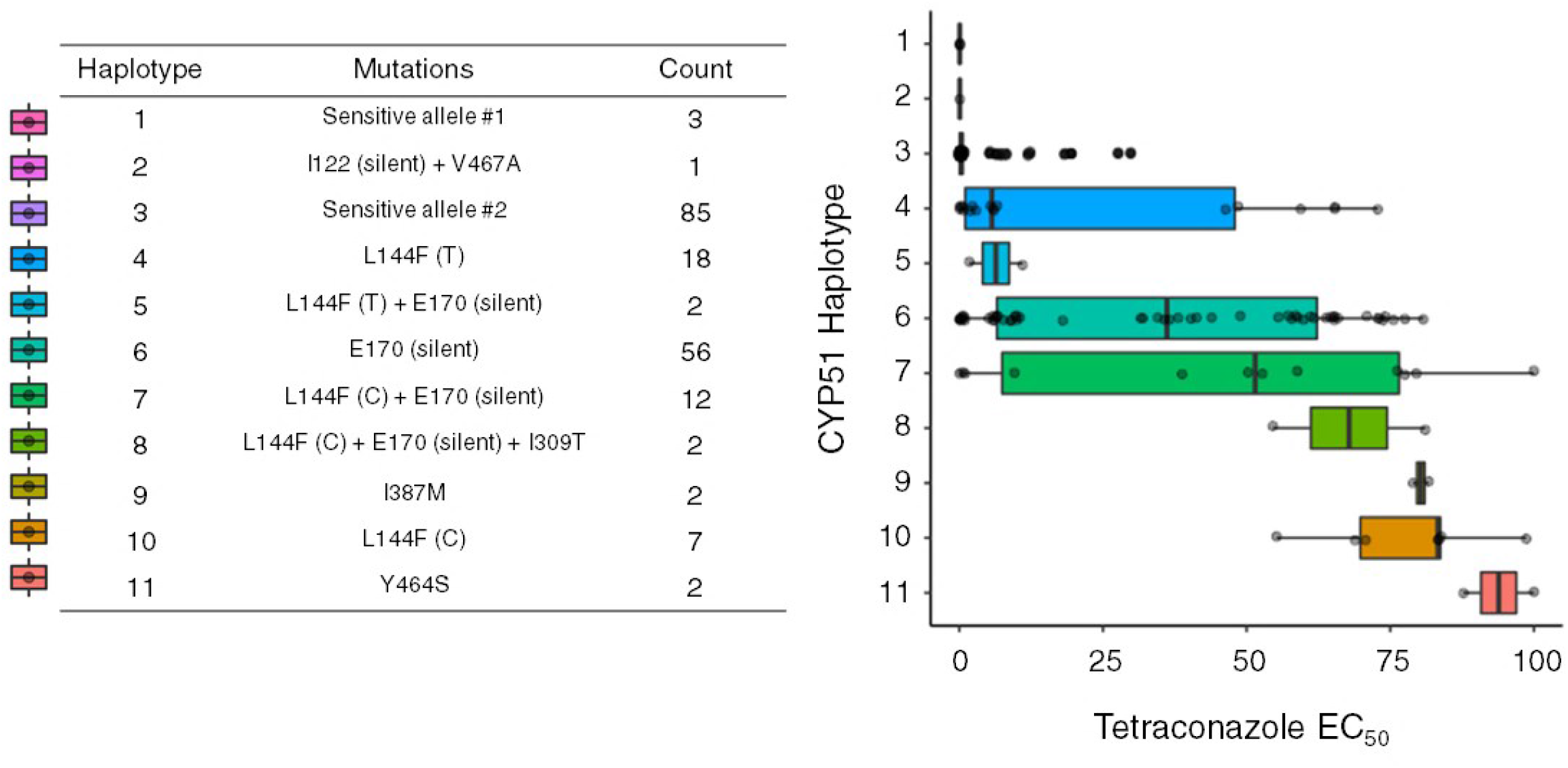
Effects of *CbCYP51* haplotypes on tetraconazole sensitivity. The left panel displays the 11 different *CbCYP51* coding sequence haplotypes found in our *Cercospora beticola* population with the respective number of isolates with each haplotype. Mutations were identified as compared to the most common sensitive Haplotype (#3). The right panel displays box and whiskers plots for tetraconazole EC_50_ values for each *CbCYP51* haplotype.

The most common amino acid substitution found was L144F (41 isolates) and could be achieved by either a T or C mutation in the 3^rd^ position of the 144^th^ codon TTG (Fig. 4, S12, S13). Both TTT and TTC versions of L144F were associated with increased tetraconazole EC_50_ values (*P* < 0.01 and *P* < 0.001, respectively), but the TTC codon had a significantly higher mean EC_50_ value than the TTT codon (*P* < 0.001) (Fig. S12). Since synonymous codons at position 144 were associated with differential tetraconazole resistance, we questioned whether codon bias might help explain this phenomenon. Consequently, we calculated genome-wide codon usage for *C. beticola* (Table S5). The phenylalanine codon TTC was found in the coding sequence 70% of the time compared to TTT, which was found in the remaining 30%. This is the largest difference in codon usage for any amino acid found in *C. beticola*.

The most common *CbCYP51* haplotype associated with resistance (56 isolates) had a single silent mutation at E170 changing the 170^th^ codon from GAG to GAA, both of which encode glutamic acid (Fig. 4). Presence of the E170 mutation in strains, all of which also had the upstream SNP 9_1452111, was associated with a significant increase in tetraconazole EC_50_ value (*P* < 0.001, Fig. S12). Since codon usage again could underscore DMI resistance, we assessed codon frequencies for glutamic acid (E) and found that the GAG codon was used slightly more often (56%) than the GAA codon (44%) (Table S5).

To further investigate the involvement of these mutations in DMI fungicide resistance, we sequenced *CbCYP51* in 52 additional *C. beticola* isolates harvested in 2019 from commercial fields in the RRV of North Dakota and Minnesota. The results corroborated the haplotype analyses of the 2016 and 2017 GWAS isolates. As before, the most common haplotype associated with resistance had the silent mutation E170 (Fig. S13). We found that the amino acid substitutions L144F and Y464S were again associated with increased tetraconazole EC_50_ values (Fig. S13). The amino acid mutation H306R was also found alongside L144F in a single 2019 isolate (Fig. S13).

### Genome-wide scan for recent selection in the North American *C. beticola* population

We next addressed if DMI fungicide application had left signals of recent selection in the *C. beticola* population. To this end we focused on the population of 89 isolates exhibiting DMI fungicide resistance to conduct a genome-wide screen for selective sweeps. A genomic region that has undergone a selective sweep is deprived of genetic variation and characterized by high LD. Since similar signals can also be caused by demographic processes, it is necessary to account for past demographic events in analyses of selective sweeps. Therefore, we first inferred the recent population history of the *C. beticola* population using a simulation approach. We assessed the likelihood of four demographic scenarios: 1) a recent population expansion, 2) a recent population bottleneck, 3) a bottleneck followed by a population expansion and 4) a population bottleneck followed by a second recent bottleneck (Fig. S14). Comparing the Akaike information criterion (AIC) values of the four models, we found evidence for a demographic scenario involving one single recent bottleneck (Table S6). The model including two consecutive bottlenecks also showed a high likelihood, while the model with population expansion was the less likely demographic scenario. We further compared the observed site frequency spectrum (SFS) to the expected SFS for each model. One hundred replicate runs of each 10,000 simulations were created to infer parameter values (Fig. S15). Using this strategy, we also found that the residuals between the expected and the observed SFS were minimized under the single bottleneck model and therefore provided further support for this scenario.

Two independent approaches to detect selective sweeps, OmegaPlus and RAiSD, identified 1,800 and 3,647 selective sweep regions, respectively, distributed across the *C. beticola* chromosomes (Fig. 5, Tables S7 and S8). These regions show significantly higher values of the statistics ω and *μ* when compared to the highest statistic values obtained from the 10,000 simulations under the neutral demographic scenario. The selective sweep regions ranged in length from 0.7 kb to 104 kb for OmegaPlus and from 1 kb to 671 kb for RAiSD, and the number of variants included in these regions varied from one to 2,754 SNPs for OmegaPlus and from one to 6,118 for RAiSD. We further compared the output of the two independent approaches of OmegaPlus and RAiSD. Although these analyses detect different signatures in the genome data, some selective sweep regions were overlapping (Fig. 5). In total we identified 8,807 overlapping regions of the two selective sweep maps (using a minimum overlap size of 150 bp) spanning from 151 bp to 7.7 kb. Most of these overlapping selective sweeps encode genes (from 0 to 55 genes) revealing a set of functional traits that have experienced recent selection (Tables S9 and S10).

**Figure 5.**
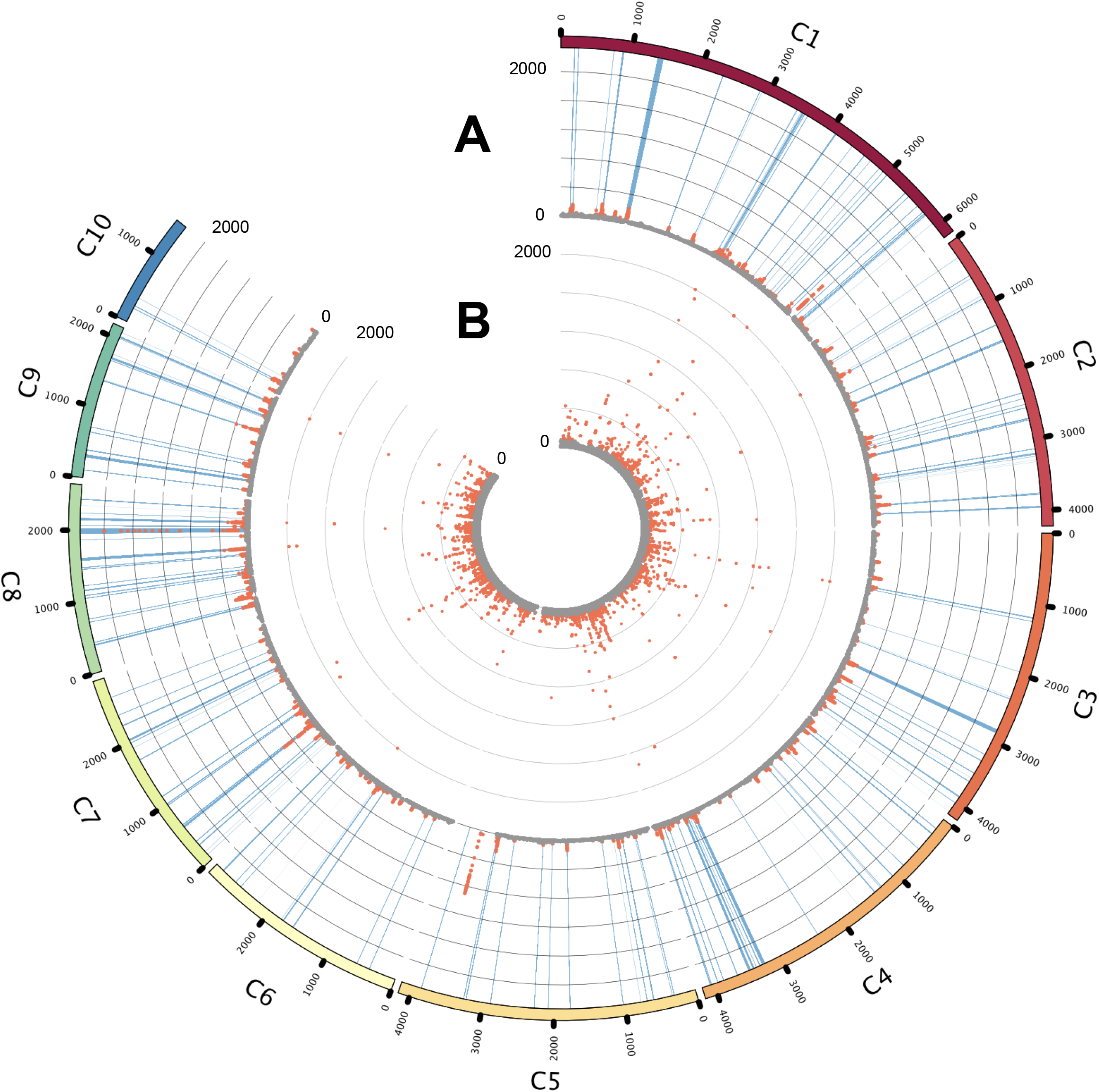
Genomic scans for selective sweeps using RAiSD and OmegaPlus. (A) Selective sweep map obtained by RAiSD, the μ-statistic values was calculated and plotted along the genome. Significant outlier loci are shown in red. (B) Selective sweep map obtained by OmegaPlus, ω-statistic values plotted across the genome. Significant outlier loci are shown in red. The significance thresholds of the μ and ω statistics were determined with demographic simulations (see Materials and Methods). Blue lines indicate selective sweeps longer than 150 bp detected with both methods.

### Signatures of recent selective sweeps co-localize with DMI fungicide resistance candidates in the *C. beticola* population

Two of the selective sweep regions on chromosome 4 and chromosome 9 identified by RAiSD contained variants identified in the GWAS analysis that correlate with DMI fungicide resistance (Table S8). On chromosome 4, nine overlapping windows with significantly higher μ values also include the gene encoding the DYRK identified through GWAS (Table S9). Merging these nine windows results in a 79,345 bp long region that starts at position 3,723,514 and ends at position 3,802,859 and thereby includes the marker identified via GWAS in the 5’UTR of the same gene, at position 3,725,552. An additional 19 predicted protein-coding sequences are located in the same region.

Another region of 7,827 bp on chromosome 9 (from position 1,495,335 to position 1,503,162) has also undergone a recent selective sweep. This region includes a part of the coding sequence of the ABC PDR transporter identified via GWAS and the intragenic SNPs at positions 1,502,965, 1,502,966, and 1,502,968 (Tables S3 and S9) that also associate with DMI fungicide resistance. An additional nine predicted genes are located in this region. Coordinates of the predicted genomic features in the selective sweep regions on chromosomes 4 and 9 are summarized in Tables S12 and S13.

### Assessing fitness penalties for DMI fungicide resistance loci

To investigate whether there was a fitness penalty associated with DMI fungicide resistance, we measured the radial growth rates of all 190 *C. beticola* strains as a proxy for fitness (Fig. S1B, Table S1). We performed association analyses for radial growth rate and obtained allelic effect estimates for each marker. There was a very weak positive correlation between allelic effects of DMI fungicide resistance (tetraconazole EC_50_ values) and radial growth rate (Pearson correlation coefficient = 0.174, *P* < 2.2e-16), indicating no fitness penalty between mycelial growth and DMI fungicide resistance (Fig. 6A). The most significant markers associated with tetraconazole resistance also appeared to have a negligible effect on growth rate (Table S14, Fig. 6A). We also performed association analyses for radial growth of isolates under salt stress (1M NaCl) (Fig. S1C). There was a slightly stronger positive correlation between the allelic effects of DMI fungicide resistance (tetraconazole EC_50_ values) and radial growth rate under salt stress (Pearson correlation coefficient = 0.322, *P* < 2.2e-16) (Fig. 6B). This suggests that some mechanisms of DMI fungicide resistance may also lead to increased salt tolerance. However, the most significant markers associated with tetraconazole resistance did not appear to have meaningful impact on growth rates under salt stress (Table S14, Fig. 6B).

**Figure 6.**
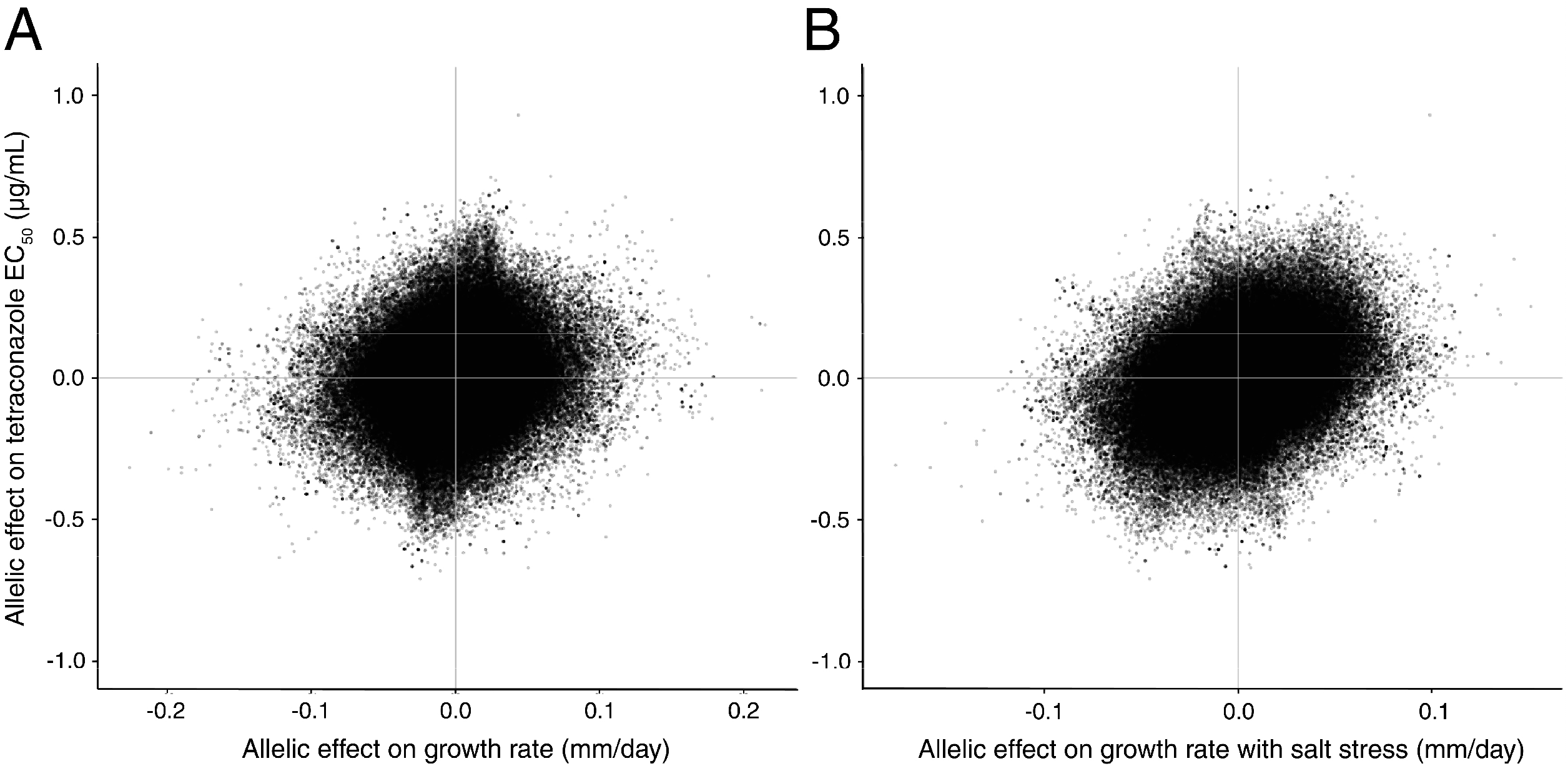
Correlation of genome-wide allelic effects on tetraconazole sensitivity (EC_50_) and radial growth rates (mm/day) of *Cercospora beticola* cultures under different conditions. The allelic effect estimates were obtained for 290,496/320,530 markers from association mapping performed in GAPIT. Pearson correlation test revealed a slight positive correlation between the allelic effects for tetraconazole sensitivity and radial growth rate (A) (coefficient = 0.1739). There was a stronger positive correlation between the allelic effects for tetraconazole sensitivity and radial growth under 1M NaCl salt stress (B) (coefficient = 0.3218).

### Discovery of a *CYP51-like* gene that does not appear to affect DMI sensitivity

The existence of multiple CYP51 paralogs has been described in several ascomycete fungi (Becher and Wirsel 2012; Fan et al. 2013; Hawkins et al. 2014; Liu et al. 2011). In the 09-40 *C. beticola* reference genome, there is a second gene annotated as Eburicol 14a-demethylase on chromosome 4 (NCBI XP_023452197.1, CB0940_05116) with 51.5% protein identity to CYP51. A protein homology of 40% is the minimum requirement for belonging to the same cytochrome P450 family (Nelson et al. 1993). Previous Southern blot analyses of *CbCYP51* suggested that a second P450 gene in *C. beticola* with some sequence similarity to *CbCYP51* (Bolton et al. 2012a). This *CbCYP51-like* gene is 1,530 bp and intron-free with a top NCBI BLASTp hit of a hypothetical protein in *Fonsecaea monophora* (NCBI XP_022511803.1) at 67.25% protein identity followed by multiple CbCYP51-like hits from *Fonsecaea, Cladophialophora, Arthrobotrys* and *Fusarium* spp. at 55-68% protein identity. Phylogenetic analyses of the *C. beticola* CYP51-like protein alongside well-characterized CYP51 proteins from other ascomycete phytopathogenic fungi placed it in a clade with *Fusarium gramineaum* CYP51C (Fig. 7). Analysis of *CbCYP51-like* gene haplotypes in 190 *C. beticola* isolates revealed that seven isolates had a deletion of 1,751 bp spanning ~750 bp upstream of the ATG and the first 999 bp of coding sequence. This could be regarded as gene absence, but the remaining coding sequence could hypothetically still encode a truncated protein of 222 amino acids since there is a methionine still in frame. Overall, the deletion suggests that *CbCYP51-like* is non-essential or is functionally compensated by a yet-unidentified gene when absent. Several amino acid substitutions were also identified but no association could be found between CbCYP51-like gene haplotype and EC_50_ value (Fig. S16).

**Figure 7.**
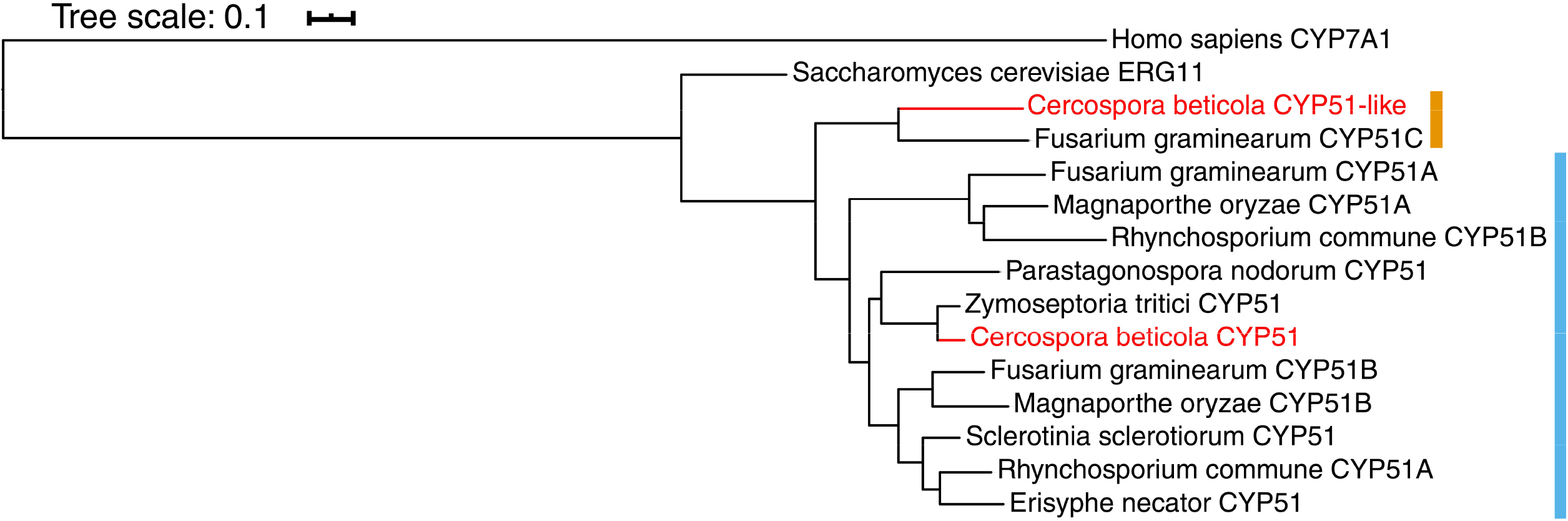
Phylogenetic analysis of *Cercospora beticola* CYP51-like protein. Maximum-likelihood phylogenetic analysis was performed for *Cercospora beticola* CYP51-like protein sequence alongside CbCYP51 and known CYP51 paralogs from closely related ascomycete species. The protein sequences for *Saccharomyces cerevisiae* ERG11 and *Homo sapiens* CYP7A1 were included as out-groups. CYP51-like is highlighted with a red line, where it forms a clade with *Fusarium gramineaum* CYP51C (clade marked in orange). CYP51 is also highlighted with a red line, where it forms a clade with the remaining ascomycete CYP51A and CYP51B sequences (clade marked in blue).

### Development of Cas9 RNP-mediated gene editing to characterize fungicide resistance mutations in *Cercospora beticola*

Current functional studies of fungicide resistance mutations within essential genes largely rely on heterologous expression of the mutant allele in yeast (Cools et al. 2010; Obuya et al. 2015). However, we hypothesized that studying the effect of synonymous mutations, such as CbCYP51 E170, would not be suitable in alternative hosts that ostensibly harbor differing codon biases. A split marker gene replacement method has been utilized for *C. beticola* (de Jonge et al. 2018; Ebert et al. 2019), but is not ideally suited for generating single nucleotide changes in target genes. In order to test the involvement of candidate fungicide resistance mutations, we decided to use a gene editing method in *C. beticola*. Cas9 ribonucleoprotein-mediated gene editing was first developed by Foster et al. (2018) in *Magnaporthe oryzae*, which was adapted here using polyethylene glycol (PEG) transformation of *C. beticola* protoplasts. Briefly, Cas9 was complexed with an *in vitro*-synthesized single guide RNA (Cas9 ribonucleoprotein) and simultaneously introduced with an ~80 bp donor DNA template (with the intended bp change in the center) into protoplasts via PEG transformation (Table S15).

Benzimidazole fungicides target and inhibit beta-tubulin in fungi, and the E198A target site mutation is known to confer high levels of resistance in *C. beticola* (Davidson et al. 2006; Rosenzweig et al. 2015; Shrestha et al. 2020; Trkulja et al. 2013). Therefore, we initially introduced the E198A beta-tubulin mutation into a benzimidazole-sensitive strain 16-1124 to establish a working gene editing method, which resulted in a benzimidazole-resistant phenotype (Fig. S17). Having established Cas9 ribonucleoprotein-mediated gene editing in *C. beticola*, we then attempted to introduce the L144F (TTG to TTC or TTT) and E170 (GAG to GAA) mutations into DMI-sensitive strain 16-100, using at least three different single guide RNA designs per mutation. However, all attempts to recover edited mutants failed, with none of the protoplasts growing through the tetraconazole selective media harboring the intended *CbCYP51* mutations. We also attempted to create these mutations using a co-editing approach, where a hygromycin resistance cassette was simultaneously introduced at a different location, allowing for simple selection on hygromycin and screening for edits at the *CbCYP51* locus. This method also failed to recover edited individuals. Taken together, these data suggest that we either did not identify the ideal guide RNA or donor template combination to generate *CbCYP51* mutations, or that the *in vivo* editing of *CbCYP51* in this fungus is lethal.

## Discussion

GWAS is a powerful method for identifying the genetic basis of complex phenotypic traits using variation existing within a population. Although it has been predominantly exploited for finding markers associated with disease in humans, the advent of cost-effective high-throughput sequencing technology is enabling the use of GWAS for a broad range of organisms (Power et al. 2017). Fungi are particularly amenable to GWAS because, like other microbes, they have relatively small genomes and high rates of recombination, which can increase the statistical power for complex traits (Falush 2016). For fungal crop pathogens, GWAS may have direct applications in managing disease through the discovery of key genes underlying virulence and fungicide resistance. Populations of filamentous fungi have previously been used in genome-wide association analyses to find genetic determinants of adaptive traits (Atwell et al. 2018; Ganeshan et al. 2018; Gao et al. 2016; Hartmann et al. 2017; Martin et al. 2020; Palma-Guerrero et al. 2013; Richards et al. 2019; Zhong et al. 2017), including several studies for DMI resistance (Mohd-Assaad et al. 2016; Pereira et al. 2020; Talas et al. 2016).

In this study, we used a population of *C. beticola* strains harvested largely from the RRV region of North Dakota and Minnesota, which represents the largest sugar beet production region in the U.S. DMI fungicides have been continually used since 1999 to manage *C. beticola* (Secor et al. 2010) and whilst it was initially thought that fitness penalties would preclude the spread of DMI resistance (Brown et al. 1986), resistance has increased over time leading to the occurrence of isolates with wide-ranging sensitivities and reduced disease control (Bolton et al. 2012b; Rangel et al. 2020; Secor et al. 2010). We inferred from principal component analyses that there was minor underlying population structure due to a cluster of tetraconazole-sensitive strains with more similar genetic backgrounds. Meanwhile, tetraconazole-resistant strains were generally more distantly related. This could be attributed to strong selection pressure exerted on North American *C. beticola* populations due to widespread and repeated use of DMI fungicides, enabling the survival and proliferation of DMI-resistant isolates, indiscriminate of genetic background. The underlying population structure explained by tetraconazole sensitivity could be confounding in downstream association mapping analyses, leading to false positive associations, and therefore underscoring why it is important to correct for this population structure in models. This observed population structure was corrected for using a kinship matrix within a mixed linear model when performing GWAS (Power et al. 2017).

GWAS identified 13 markers on chromosomes one, four and nine associated with tetraconazole sensitivity. These markers were associated with five distinct genes, four of which represent newly described DMI fungicide resistance mechanisms for *C. beticola* (Fig. 8). The most significant association was an insertion within the 5’UTR of a gene encoding a regulator of G-protein signaling (RGS) protein CB0940_00689 on chromosome one. The insertion appears to reduce tetraconazole sensitivity and may act through up-regulation of the RGS protein. Similar RGS proteins with phosphoinositide binding Phox homology PXA and PX domains have been identified and characterized in multiple filamentous fungi (Kim et al. 2017; Wang et al. 2013; Zhang et al. 2011). Due to their intracellular signaling roles, they may be required for multiple important processes such as sexual and asexual development, stress responses, secondary metabolism and virulence (Kim et al. 2017; Zhang et al. 2011). RgsC in *Aspergillus fumigatus* was shown to be required for regular growth, asexual development, oxidative stress response, cell wall stress response, virulence and external nutrient sensing (Kim et al. 2017). MoRgs4 in *M. oryzae* was also found to have roles in regulation of mating, conidiation, appressorium formation and pathogenicity (Zhang et al. 2011). In the case of *C. beticola*, it is possible that altered expression of the RGS domain protein may affect DMI sensitivity pleiotropically through modulation of multiple pathways such as growth, stress responses and/or metabolism such as ergosterol biosynthesis.

**Figure 8.**
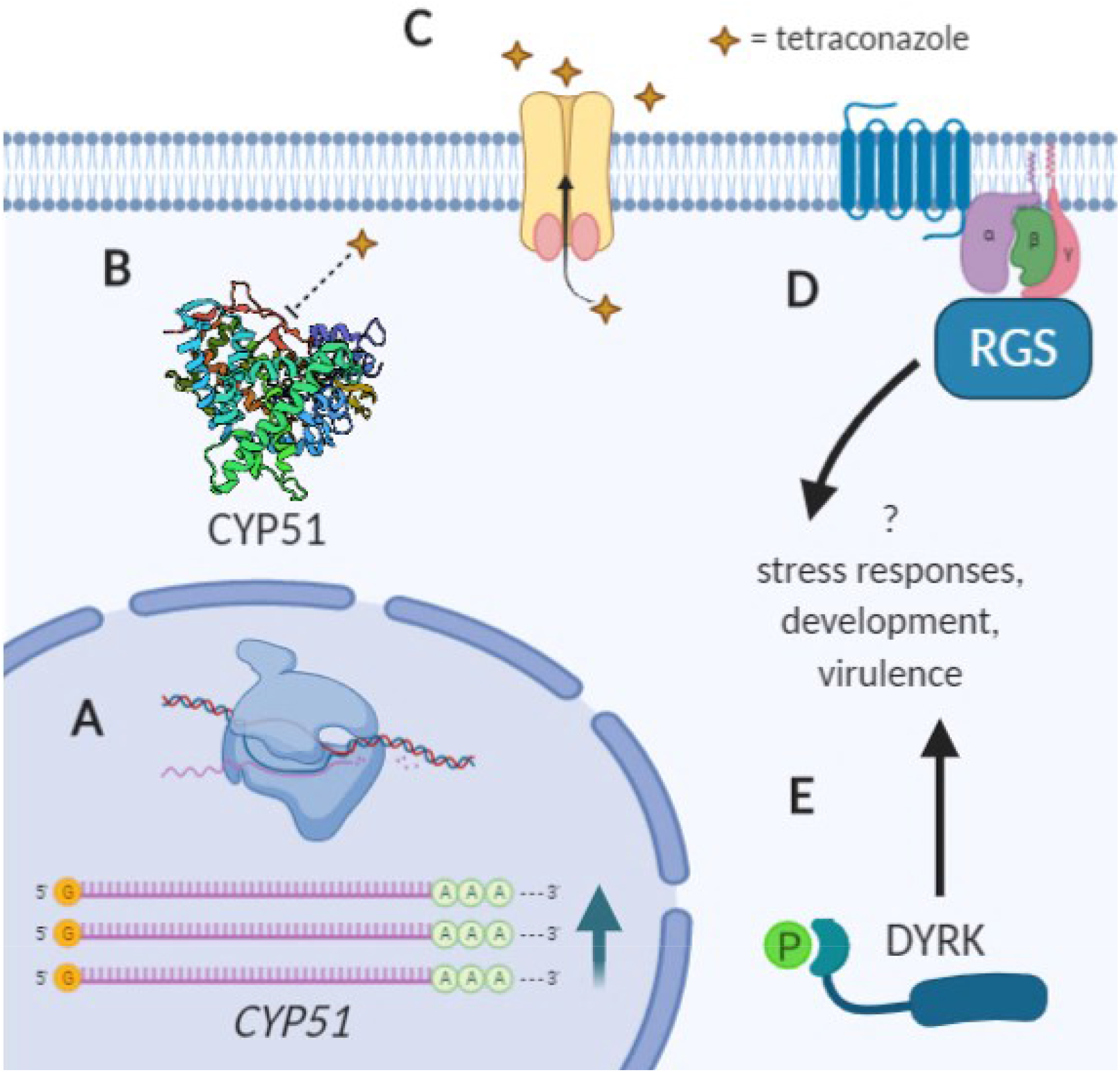
Putative mechanisms of DMI fungicide resistance in *Cercospora beticola*. *CbCYP51* induced overexpression gives rise to resistance (A) (Nikou *et al*. 2009, Bolton *et al*. 2012, 2016), as well as (B) amino acid substitutions in CbCYP51, leading to weakened binding and inhibition by DMIs (Trkulja *et al*. 2017, Shrestha *et al*. 2020). Multidrug transporters such as the ABC PDR transporter identified in this study may be pumping DMIs out across the membrane in a non-specific manner (C). There may be overexpression of ABC/MFS transporters such as seen in RNAseq study by Bolton *et al*. 2016, or amino acid substitutions to improve export of DMIs. Cellular signaling may also play a role in DMI resistance (D, E). Differential expression of a regulator of G-protein signaling (RGS) domain protein (D) or increased expression of a DYRK-type protein kinase (E) may increase DMI resistance through signaling pathways that modify expression of downstream genes such as those involved in growth, development, stress responses and/or ergosterol biosynthesis (such as *CbCYP51*). Figure created with BioRender.com.

A second gene on chromosome 1 was significantly associated with tetraconazole sensitivity. The presence of an amino acid substitution (G474S) in hypothetical protein CB0940_00999 significantly increases tetraconazole EC_50_ value, even when it co-occurs with CbCYP51 amino acid substitutions, suggesting there may be epistatic interactions between the hypothetical protein and CbCYP51. Since the protein lacks conserved domains, it is difficult to hypothesize its function in DMI resistance.

On chromosome 9, nine SNPs within ABC pleiotropic drug resistance (PDR) transporter gene CB0940_11399 were significantly associated with tetraconazole sensitivity. Whilst these all yielded synonymous mutations or were within introns, there are multiple other intragenic mutations nearby in complete LD that give rise to amino acid substitutions. Most of the mutations were found within the first ~250 of 1,493 amino acids in the protein and may alter transporter function, leading to differential pumping of tetraconazole out of fungal cells (Holmes *et al*. 2002). ABC transporters, namely those with multidrug resistance (MDR) or pleiotropic drug resistance (PDR) domains, have been implicated previously in non-specific efflux of DMI fungicides in pathogenic fungi (Hamamoto et al. 2001; Hayashi et al. 2002b). Whilst overexpression of ABC transporters is a common mechanism of drug resistance in fungi (Cannon et al. 2009; Hamamoto et al. 2001; Hayashi 2003; Hellin et al. 2018; Kretschmer et al. 2009; Nakaune et al. 1998; Slaven et al. 2002), there has been evidence for the involvement of point mutations affecting substrate binding and function as well (Holmes et al. 2006).

The other significant locus on chromosome nine was a SNP 124 bp upstream of the DMI target *CbCYP51*. It is possible that a single base pair change upstream of the gene could affect binding of a transcriptional regulator and in turn affect gene expression (Buckland 2006; Chen et al. 2016). Indeed, *CbCYP51* overexpression was previously associated with DMI resistance in *C. beticola* (Bolton et al. 2012a; Bolton et al. 2016; Nikou et al. 2009). When we investigated LD in the region, we found that the silent mutation E170 in *CbCYP51* was in high LD with the upstream SNP 9_1452111. However, we could not uncouple the effects of these mutations on tetraconazole EC_50_ value since the upstream SNP was always found with either E170 or a non-synonymous mutation in *CbCYP51*. We investigated the possibility of the upstream SNP influencing transcription factor binding sites using likely DNA-binding sites derived from *S. cerevisiae* (JASPAR database). Predicted binding sites differed for both versions of the upstream SNP, suggesting that differential *CbCYP51* regulation could occur. Differential binding site likelihoods were found for six different transcription factors including UPC2, ECM22 and STB5 which have all been described in other fungi as transcriptional regulators of *CYP51* and ergosterol biosynthesis (Flowers et al. 2012; Silver et al. 2004; Vik and Rine 2001), or pleiotropic drug resistance (Noble et al. 2013). It is possible that homologous transcriptional regulators in *C. beticola* differentially bind to the *CbCYP51* promoter due to the SNP (9_1452111), leading to up-regulation and reduced DMI sensitivity.

The fifth most significantly associated locus was an indel in the 5’UTR of DYRK CB0940_05141 on chromosome 4. The insertion appears to contribute to increased tetraconazole EC_50_ value. Intriguingly, RNAseq (Bolton et al. 2016) revealed this gene was differentially up-regulated in a tetraconazole-resistant strain in response to tetraconazole exposure when compared to a sensitive strain. Together, this suggests that the 5’UTR insertion c auses up-regulation of the protein kinase gene in response to tetraconazole. DYRK-type protein kinases are an evolutionarily conserved family of protein kinases which regulate cell growth and differentiation, such as Pom1 in fission yeast (Becker et al. 1998). DYRK family members phosphorylate many different substrates, including critical regulators of the cell cycle, and downstream effects include increased transcription factor activity, modulation of subcellular protein distribution and regulation of enzyme activity (Becker 2012). The increased expression of this DYRK in *C. beticola* could lead to reduced DMI sensitivity through multiple pleiotropic effects downstream of the signaling pathway.

We further found that DMI fungicide resistance loci within the ABC PDR transporter and DYRK were within putative selective sweep regions. Using DMI-resistant *C. beticola* isolates identified in this study, we show that genome-wide selective sweeps overlap with some loci associated with DMI fungicide resistance, suggesting that application of these fungicides has been a recent selection pressure for the North American population. This also suggests that the DMI-resistance mutations localized in selective sweep regions have been conferred a strong fitness advantage to *C. beticola*. For example, the favorable allele of the ABC PDR transporter may provide resistance to additional fungicide classes used to control *C. beticola*, in addition to the DMIs, since efflux pumps often give rise to multidrug resistance (Gulshan and Moye-Rowley 2007). Genome-wide selection scans have been performed previously for phytopathogenic fungi, including *Zymoseptoria tritici* (Hartmann et al. 2020), and they also found that multiple fungicide resistance loci identified in GWAS (50%) overlapped with selective sweep regions.

We demonstrated that the 13 tetraconazole sensitivity markers identified through GWAS had little effect on radial growth rate, with or without salt stress. Considering fungal growth rate as a proxy for fitness, this suggested the absence of a fitness penalty for these DMI fungicide resistance loci. This was corroborated by finding no clear correlation between genome-wide allelic effects on tetraconazole EC_50_ values and radial growth rates on unamended media. Nikou et al. (2009) found no differences in radial growth, sporulation, or pathogenicity on sugar beet for five European *C. beticola* isolates resistant to DMI fungicide epoxiconazole (3 μg/mL < EC_50_ < 6 μg/mL), when compared to a sensitive isolate (EC_50_=0.05 μg/mL). In another study by Bolton et al. (2012b), US *C. beticola* isolates grouped by low, medium or high tetraconazole EC_50_ values showed no differences in CLS severity on sugar beet. However, an older study by Karaoglanidis et al. (2001) showed that European *C. beticola* isolates resistant to flutriafol (2.7 μg/mL ≤ EC_50_ ≤ 15.6 μg/mL) had significantly lower spore production and virulence on sugar beet than sensitive isolates (0.10 μg/mL ≤ EC_50_ ≤ 0.34 μg/mL). Competition assays performed on sugar beet in the field between sensitive and resistant isolates also resulted in a significant reduction in frequency of resistant isolates (Karaoglanidis et al. 2001). It is possible that a trade-off exists for DMI fungicide resistance in *C. beticola*, but we have not observed it with the proxy phenotype and conditions tested in this study.

Genome-wide allelic effects on radial growth rate under salt stress (1M NaCl) had a slight positive correlation with allelic effects on tetraconazole EC_50_. This suggested that some mechanisms conferring increased DMI fungicide resistance may also lead to enhanced salt tolerance. Fungal strategies for overcoming salt stress include morphological changes, cell wall reinforcement and accumulation of osmolytes such as glycerol (Liu et al. 2017). In some fungal species, there are common signaling components controlling both salt stress responses and antifungal drug resistance (Hayes et al. 2014), including the high osmolarity glycerol (HOG) response pathway (Kim et al. 2011; Zhang et al. 2002) and calcineurin signaling pathway (Jacob et al. 2015; Juvvadi et al. 2014). Genetic variants leading to enhanced signaling in response to stress may lead to both increased salt and fungicide tolerance in *C. beticola*.

With the possibility of *CbCYP51* mutations being associated with DMI resistance, we examined the coding sequence within all 190 GWAS isolates and an extra 52 *C. beticola* isolates collected in 2019. In this study, there were two main haplotypes associated with low tetraconazole EC_50_ values: a common haplotype found in 85 isolates, and a highly divergent sequence with 114 polymorphisms shared by three isolates. Since *C. apii* is morphologically indistinguishable from *C. beticola* (Groenewald et al. 2005), we questioned whether these three isolates might be another species. However, whole genome phylogenetic analyses, as well as analysis of calmodulin, TEF and actin genes identified these isolates as *C. beticola* (data not shown). Compared to the most common sensitive haplotype, there were two synonymous mutations (I122 and E170) and six different non-synonymous mutations: L144F, H306R, I309T, I387M, Y464S and V467A. The L144F, I387M and Y464S mutations were associated with increased tetraconazole EC_50_ value and so may be directly involved in DMI resistance.

In fungal human and plant pathogens, evolutionary flexibility of CYP51 to accept structural changes has often led to the accumulation of amino acid changes and selection of haplotypes that reduce DMI binding and inhibition (Becher and Wirsel 2012). The amino acid substitutions L144F, I387M and Y464S were previously reported in *C. beticola* strains from Serbia and, as in our study, were individually associated with DMI resistance (Trkulja et al. 2017). Mair et al. (2016) proposed a system for unifying labelling of amino acids in CYP51 through alignments between CYP51 proteins in relevant species fitted to the well-studied *Z. tritici* ‘archetype’. Orthologous amino acids in all species can then be consistently labelled based on the position of the amino acid in the archetype protein. We found that L144F was the most common CbCYP51 amino acid change in RRV *C. beticola* isolates from 2017, 2018 and 2019, which has not been reported in orthologous sites in other fungal species (Mair et al. 2016). The I387M mutation (haplotype 9) also does not appear to have orthologous mutations in other fungi. However, the Y464S mutation appears to be analogous to Y461S/G/H in *Z. tritici* (Cools and Fraaije 2012; Mair et al. 2016). In *Z. tritici*, changes in residues Y459 to Y461 occur frequently (Cools and Fraaije 2012). Additionally, alterations in equivalent residues in Y459 to Y461 have been found in *A. fumigatus* (Howard et al. 2006), *C. albicans* (Perea et al. 2001) and *Mycosphaerella fijiensis* (Cañas-Gutiérrez et al. 2009), all of which were associated with increased resistance to DMIs. Expression of *ZtCYP51* encoding Y461H in *S. cerevisiae* confers decreased sensitivity to all DMIs (Cools et al. 2010). Molecular modelling predicted this residue to be integral to the CYP51 active site with alterations directly impacting DMI binding (Mullins et al. 2011). Despite the widespread association of residues Y459 to Y461 to DMI resistance in fungal species, the Y464S amino acid exchange was not common in our study with only two isolates harboring this mutation.

To the best of our knowledge, we also present three novel CbCYP51 amino acid substitutions in *C. beticola*, H306R, I309T and V467A. The impact of these relatively rare mutations is still unclear. A single isolate from 2019 harbored H306R and it was present alongside L144F which is individually associated with DMI resistance. Only two isolates had I309T and the substitution was co-present with L144F and E170, which are both individually associated with DMI resistance. No analogous mutations for H306R or I309T have been found in other fungal species (not shown). Likewise, only one isolate in our study had the V467A substitution and it was present alongside the silent mutation I122. Intriguingly, this isolate was highly sensitive to DMIs, suggesting that V467A, although close to the Y464 residue, does not affect binding to CbCYP51.

Unexpectedly, we discovered a potential codon usage effect for the L144F substitution in CbCYP51. We observed that strains with L144F encoded by the TTT codon had a significantly lower EC_50_ value than strains with L144F encoded by the TTC codon. We did not find another mutation within or close to *CbCYP51* (±1 kb) in LD with the codon difference. In *C. beticola*, the phenylalanine codon TTT is used just 30% of the time in coding sequence when compared to the codon TTC at 70%, representing the biggest difference in codon usage for a single amino acid in *C. beticola*. The model fungus *N. crassa* exhibits a similar codon bias for phenylalanine with TTC used in ~67% of cases (Kazusa codon usage database). The use of rare versus optimal codons in *N. crassa* has been shown to impact transcript levels (Zhou et al. 2016; Zhou et al. 2018), protein abundance (Zhou et al. 2015) and co-translational folding of proteins (Yu et al. 2015). Since we did not observe differences in *CbCYP51* expression levels between L144F strains (data not shown), we suspect that the codon usage here instead impacts CbCYP51 protein folding or abundance. It has long been suggested that codon usage regulates protein folding by affecting the rate of translation (Goldman et al. 1995; Komar et al. 1999; Konigsberg and Godson 1983; Sørensen et al. 1989; Zhou et al. 2009). For example, a synonymous mutation to a rare codon within the human *multidrug resistance 1* (*MDR1*) gene results in an altered protein conformation and modified drug interactions (Kimchi-Sarfaty et al. 2007). The change in isoleucine codon from ATC (47% usage) to rarer codon ATT (35% usage) gave differential MDR1 protein conformation and decreased the ability of an efflux blocker to inhibit MDR1. An even rarer isoleucine codon ATA (18% usage) further decreased the inhibitor effects on MDR1. The authors hypothesized that the effect was due to codon context; when frequent codons are changed to rare codons within a cluster of rare codons, the timing of co-translational protein folding is affected (Anthony and Skach 2002). Zhou et al. (2015) used the filamentous fungus *N. crassa* and manipulation of codons in the circadian clock gene *frequency* (*frq*) to demonstrate a correlation between codon usage and protein disorder tendency. In this context, rarer codons are preferentially used in intrinsically disordered regions whereas more optimal codons are used in structured domains. We hypothesize that the phenylalanine codon TTT at position 144 of *CbCYP51* in *C. beticola* yields a different protein structure than codon TTC, which in turn causes differential binding of DMIs. Another possibility is that the optimized TTC codon leads to a higher overall translation rate of CbCYP51 (due to tRNA availability) and the higher protein level results in reduced DMI sensitivity. Functional studies will be necessary to confirm these hypotheses.

Intriguingly, we identified a silent mutation (E170) associated with DMI resistance in our study. Obuya et al. (2015) also associated this mutation with DMI resistance using RRV isolates, and it was also previously associated with resistance in *C. beticola* in isolates from Greece (Nikou et al. 2009) and Serbia (Trkulja et al. 2017). Obuya et al. (2015) heterologously expressed a *C. beticola CYP51* haplotype harboring E170 in *S. cerevisiae* strain R1 lacking multidrug resistance transporter Pdr5 (ΔPdr-5) and found no change in DMI sensitivity. However, it is possible that this mutation has a *C. beticola*-specific influence on DMI sensitivity through codon usage, and thus functional studies in alternative hosts may not be conclusive. For glutamic acid (E), the GAG codon seen in more DMI-sensitive strains is used slightly more often (56%) than the GAA codon (44%). It is possible that codon usage in this context leads to differential co-translational CbCYP51 folding, protein structure and DMI binding as suggested above for L144F. Since the GAA codon found in resistant strains is the nonoptimal codon, it seems unlikely that it would increase the translation rate and CbCYP51 protein levels. Another possibility is that the synonymous change influences DMI resistance via *CbCYP51* expression levels e.g. via promotion of premature transcription termination (Zhou et al. 2018), chromatin structure (Zhou et al. 2016), mRNA stability (Duan and Antezana 2003) or even small RNA-based gene regulation (Lee et al. 2010). Interestingly, a synonymous mutation in *CbCYP51* from the grapevine powdery mildew pathogen *Erysiphe necator* was only identified in isolates with high *CbCYP51* expression and increased resistance to DMIs (Rallos and Baudoin 2016). In humans, over 50 diseases are associated with synonymous mutations (Sauna and Kimchi-Sarfaty 2011) which have been linked to aberrant splicing, mRNA stability or codon bias effects. There are also at least five synonymous mutations in *Escherichia coli* protein TEM-1 β-lactamase that confer higher total and functional protein levels and increase resistance to cefotaxamine (Zwart et al. 2018).

Nikou et al. (2009) found that four highly DMI-resistant isolates harbored E170 and also overexpressed *CbCYP51*. Alternatively, the E170 mutation in RRV strains is in high LD with the upstream SNP 9_1452111, or other mutations, which themselves could affect *CbCYP51* gene expression and be directly involved in DMI resistance. However, since isolates from disparate locations including the RRV (Obuya et al. 2015), Greece (Nikou et al. 2009), and Serbia (Trkulja et al. 2017) have identified an association between E170 and DMI resistance, it is tempting to speculate a direct involvement between this mutation and DMI resistance. Functional studies will be necessary to confirm the involvement of the upstream SNP and/or E170 with DMI resistance.

In some ascomycete fungi, there are two *CYP51* paralogs known as *CYP51A* and *CYP51B* that both have sterol 14 α-demethylase function in ergosterol biosynthesis (Brunner et al. 2016). This is the case in multiple plant pathogenic fungi such as *R. commune* (Hawkins et al. 2014), *Pyrenophora teres* f. sp. *teres* (Mair et al. 2016)*, F. graminearum* (Liu et al. 2011), *Penicillium digitatum* (Sun et al. 2011) and *Colletotrichum* spp. (Chen et al. 2020). DMI use can impose selection on favorable mutations within or upstream of either paralog. We identified a possible *CbCYP51* paralog, *CbCYP51-like*, in the 09-40 *C. beticola* reference genome which had 51.4% protein similarity to CbCYP51. Analysis of *CbCYP51-like* haplotypes in our population suggests that the gene is not associated with sensitivity to tetraconazole or other DMI fungicides tested.

Seven isolates had a deletion in *CbCYP51-like* giving partial gene absence, suggesting that the gene is non-essential or that functional redundancy exists. Phylogenetic and BLASTp analyses of protein sequences demonstrated that CbCYP51-like is more similar to CYP51C found in *Fusarium* spp. than CbCYP51 from *C. beticola* or paralogs from more closely related ascomycetes. The *CYP51C* gene was previously considered as being unique to *Fusarium* spp. (Fernández-Ortuño et al. 2010). Fan et al. (2013) demonstrated that CYP51C in *F. graminearum* had functionally diversified from the other CYP51A and CYP51B, since it failed to complement yeast CYP51 function and it was required for full virulence on wheat ears. It is possible that CbCYP51-like also lacks sterol 14α-demethylase function and plays another role in the fungus, which could be elucidated through the generation of deletion mutants.

As mentioned above, GWAS is a powerful tool to identify genomic variants associated with a trait of interest. However, to establish causality, the gold standard is functional validation. Gene replacement methods are useful for establishing gene function. For genes of known function, such as *CYP51*, we are often interested in the effects of specific mutations that require other methods such as heterologous expression of *CYP51* haplotypes in yeast (Cools et al. 2010). However, it is possible that mutations only give an observed phenotypic change when expressed in the native species e.g. due to codon usage bias. Genome editing provides an elegant way of introducing desired mutations into a target gene in the native species, without the need for co-introduction of a selectable marker if the mutation itself is selectable. Therefore, this method lends itself to investigating mutations involved in fungicide resistance. The *CYP51* gene has been edited in human fungal pathogens *Aspergillus fumigatus* (Umeyama et al. 2018) and *Candida* spp. (Morio et al. 2019) which exhibit azole resistance. To our knowledge, this has not yet been done in phytopathogenic fungi. We adapted a Cas9-RNP gene editing method developed by Foster et al. (2018) for use in *C. beticola* to functionally assess fungicide resistance mutations. To establish this method in our laboratory, we first edited the beta-tubulin gene to introduce the E198A mutation associated with benzimidazole resistance (Davidson et al. 2006; Shrestha et al. 2020).We successfully introduced the E198A mutation into a benzimidazole-sensitive *C. beticola* strain and recovered mutants using selection on benzimidazole-amended media. Despite these initial successes, we were unable to introduce L144F (TTT or TTC) or E170 mutations into CbCYP51. Likewise, attempts to co-edit through simultaneous introduction of a hygromycin-selectable marker at a different genomic location were also not successful. It is unclear why it was not possible for us to edit *CbCYP51*, but we conclude that we either did not find optimal sgRNA and template sequences or that editing the essential gene *CbCYP51* is lethal in *C. beticola*. An alternative strategy would be to generate a viable Δ*CbCYP51* knockout mutant by supplementing the media with ergosterol and transforming the mutant with different *CbCYP51* haplotypes to test their effects.

To conclude, association mapping and selective sweep analyses were used together for the first time in a *Cercospora* species. We identified mutations on chromosomes 1, 4 and 9 associated with DMI fungicide resistance within or upstream of five different genes: an RGS domain protein, a DYRK-type protein kinase, an ABC transporter, a hypothetical protein and the DMI target, CbCYP51. Mutations in the ABC transporter and DYRK genes overlapped with selective sweep regions, suggesting that DMI fungicide application has been a recent selection pressure for the population. There was no evidence of a fitness trade-off for DMI fungicide resistance using radial growth as a proxy. Haplotype analysis of *CbCYP51* demonstrated that intragenic mutations E170, L144F, I387M and Y464S are significantly associated with tetraconazole resistance. Future studies should establish if the mutations identified are directly involved in DMI fungicide resistance and clarify the role of *CbCYP51* overexpression. Cas9-RNP editing, although successful for beta-tubulin in *C. beticola*, did not allow us to functionally assess *CbCYP51* mutations. Overall, we have demonstrated that GWAS was useful even for local populations of *C. beticola*. The identification of markers associated with DMI resistance has allowed for the development of methodologies to identify resistant strains (Shrestha et al. 2020), which was a major goal for this study. Moreover, the available isolate genotyping data can be used in future GWAS studies to establish the genetic architecture of other traits of importance, including virulence on the sugar beet host.

## Materials and Methods

### Field sampling of *C. beticola*

The 190 *C. beticola* isolates were collected from sugar beet leaves harvested from naturally infected commercial fields in the RRV region of Minnesota and North Dakota, and Idaho (n=2), in 2016 (n=142) and 2017 (n=48) (Table S1). Conidia from the lesions on the sugar beet leaves were liberated into 30 μL sterile water (0.02% ampicillin) with a pipette tip and transferred to water agar plates (1.5% w/v agar (Sigma-Aldrich) and 0.02% ampicillin). Conidial germination occurred after 24 h at 22°C and then single conidia were transferred to Difco^™^ potato dextrose agar (PDA) plates which were incubated at 25°C for 14 days. These PDA plates served as source inoculum for all further studies for each of the 190 single spore-derived isolates. Of the 142 isolates collected in 2016, 62 were collected from two adjacent fields. Random representative sampling of strains was performed in these two fields, as outlined by McDonald (1997), by walking diagonally across the field and collecting a leaf every meter from a corner to the center of the field. Prior to fungicide application, 60 diseased leaves were harvested from each field and 100 leaves were harvested from each field post-application of tetraconazole (Eminent fungicide). All isolates collected from these two adjacent Fargo fields were genotyped using eight microsatellite markers to remove any potential clones, as described by Vaghefi et al. (2016), which led to the selection of 62 as part of the final population (n=190) (Table S1). The remaining isolates collected in 2016 (n=80) and 2017 (n=48) were obtained as part of annual *C. beticola* fungicide resistance surveys in the RRV region, where growers send infected sugar beet leaves to the Secor lab at North Dakota State University for fungal isolation and sensitivity testing.

### Phenotyping for DMI fungicide sensitivity

To measure sensitivity to the DMI fungicide active ingredient tetraconazole, EC_50_ values were calculated from radial growth of the *C. beticola* isolates on amended media, as described by Secor and Rivera (2012). The single spore subcultures for all 190 isolates were transferred to clarified V8 (CV8) medium plates (10% v/v clarified V8 juice (Campbell’s Soup Co.), 0.5% w/v CaCO3, 1.5% w/v agar (Sigma-Aldrich)) and incubated at 20°C for 15 days in a continuous light regime. An agar plug of 4 mm in diameter was excised from the growing edge of the colony and placed in the center of a set of CV8 plates: one non-amended control plate and the rest amended with serial ten-fold dilutions of technical grade tetraconazole (active ingredient of Eminent 125SL (Sipcam Agro)) from 0.001 to 100 μg/mL. All plates were incubated in the dark at 20 °C for 15 days after which two perpendicular measurements were made across the colonies and the diameter averaged. The percentage reduction in growth compared to the non-amended media was calculated for each tetraconazole concentration. The EC_50_ value for each isolate was calculated by plotting the percentage reduction in growth against logarithmic tetraconazole concentration and using regression curve fitting to find the tetraconazole concentration that reduced growth by 50%. Statistical analysis was performed in RStudio (Team 2015) and was comprised of one-way ANOVA followed by a post-hoc Tukey test to identify significant differences between groups.

### Radial growth assays

All 190 isolates were grown on CV8 plates for 15 days at 20°C in a continuous light regime, as described above. An agar plug of 4 mm in diameter was taken from the leading edge of these cultures and transferred to a new CV8 plate. Three unamended CV8 plates were initiated per isolate, and these were grown at 23°C under continuous light. The radius of each culture was measured after 2, 6, 9, 13 and 16 days and a mean value was calculated for each day. Three CV8 plates amended with 1M NaCl were also initiated per isolate and grown under the same conditions. The radius was measured for these cultures after 6, 9, 12, 16, 20 and 23 days and a mean value was calculated for each day. Linear regression of radius (mm) versus time (days) was performed using SAS^®^ software, to establish the rate of radial growth in mm/day for both unamended and salt stress conditions.

### DNA extraction and whole genome resequencing

High quality genomic DNA was extracted for library preparation from liquid cultures of *C. beticola*. A single 6 mm agar plug excised from the source PDA plate was sliced into small pieces and used to inoculate 100 mL Difco^™^ potato dextrose broth (PDB). Cultures were grown at 25 °C for seven days, shaking at 150 rpm. The mycelia were filtered through Miracloth, flash frozen in liquid nitrogen and ground into a fine powder using a mortar and pestle. The method of Zhang et al. (1996) was followed for large scale isolation of genomic DNA but replacing the chloroform:isoamyl (24:1) with phenol:chloroform:isopropanol (25:24:1). The resultant DNA was then cleaned up further using the Qiagen DNeasy Plant Mini Kit (Cat No. 69106) according to manufacturer’s instructions. DNA samples were sent to Beijing Genome Institute (BGI) for library preparation (400 bp inserts) and 100 bp or 150 bp paired-end whole genome resequencing using the Illumina HiSeq 4000 platform to achieve approximately 25X genome coverage per isolate. All sequencing reads were deposited in the NCBI short read archive under BioProject PRJNA673877.

### Variant calling

Sequencing read quality was analyzed using FastQC (Andrews 2017) and Trimmomatic (Bolger et al. 2014) was subsequently used to trim reads (HEADCROP:10) and remove unpaired reads. The trimmed reads were aligned to the reference *C. beticola* 09-40 genome (NCBI RefSeq assembly GCF_002742065.1) (de Jonge et al. 2018) using BWA-MEM (Li 2013). SAMtools (Li et al. 2009) was used to convert the output sam files to sorted, indexed bam files and to index the reference genome. Duplicate reads (PCR and optical) were removed from bam files using Picard MarkDuplicates (Institute 2016). Genome Analysis Tool Kit (GATK) V4.0.8.1 HaplotypeCaller (McKenna et al. 2010) was used to identify SNPs and small indels between each isolate and the 09-40 Reference sequence. GATK CombineGVCFs was used to combine all HaplotypeCaller gVCFs into a multi-sample gVCF which was genotyped by GATK GenotypeGVCFs to produce the final VCF. Vcftools (Danecek et al. 2011) was then used to filter variants for genotyping quality (--minGQ 10) and sequencing depth (--minDP 3).

### Population structure and LD decay analyses

Before performing principle component analysis (PCA), the VCF was filtered using Vcftools to retain SNPs only (--remove-indels). Plink (Chang et al. 2015) was used to prune the SNPs for linkage disequilibrium (LD), with option −indep-pairwise 50 10 0.1, to analyze pairwise association between SNPs (r^2^) in chromosomal windows of 50 SNPs at a time and removing pairs with r^2^ > 0.1 before shifting the window 10 bp. PCA was performed using the SMARTPCA function as part of the EIGENSOFT package (Patterson et al. 2013) and plots of PCs were created using PCAviz (Novembre et al. 2018).

To assess LD decay within the mapping population, the PopLDdecay software version 3.40 (Zhang et al. 2019) was used to estimate pairwise linkage disequilibrium (R^2^) between all SNPs within 10 kb of each other and this was plotted as LD decay.

### Association mapping

Association mapping was performed using GAPIT v3.0 (Wang and Zhang 2018). The imported genotyping VCF was first filtered in TASSEL v5.0 (Bradbury et al. 2007) to convert heterozygous calls to missing data and to establish a minor allele frequency of 0.05 and minimum SNP count of 171/190 (10% missing). As the tetraconazole EC_50_ phenotype had highly positive skewing (not normally distributed), all values were log_10_ transformed prior to association mapping. A general linear model was run as a naïve model and as a model incorporating the optimal number of components derived from PCA as fixed effects to correct for population structure. GAPIT BIC was used to determine the optimum number of PCs for the model. A mixed linear model was selected for both tetraconazole sensitivity and radial growth phenotypes, incorporating a kinship matrix (K, calculated using the default VanRaden algorithm) as a random effect. The most robust model for a trait association was selected through visualization of the quantile-quantile (Q-Q) plots, which show the relationship between observed and expected *P*-values. The R package qqman (Turner 2014) was used to generate Manhattan and Q-Q plots. Allelic effect estimates for phenotypes were derived from association mapping in GAPIT. R v.4.0.2 was used for the Pearson’s product-moment correlation test.

### Evaluation of associated loci

To assess LD at significantly associated loci, LDheatmap (Shin et al. 2006) was used to plot color-coded values of pairwise LD (R^2^) between all markers in the filtered VCF ±3 kb of the significantly associated marker. SNPEff (Cingolani et al. 2012) was used to predict the effects of associated mutations within genes.

### Inference of demographic history

Prior to the scan of selective sweeps along the *C. beticola* genome, we computed the site frequency spectrum (SFS) to infer the demographic history of the population of isolates showing DMI fungicide resistance. Our analysis was based on the fit of four demographic models (Fig. S7) to the observed frequency spectrum of minor alleles (Minor Allele Frequency Spectrum, MAFS). We extracted the MAFS from the VCF file obtained from the population genomic dataset, and filtered the dataset to include only SNPs with at least 1 kb distance to predicted coding sequences and 0.3 kb distance from each other to minimize the effects of selection and linkage disequilibrium. We used the data conversion tool PGDSpider (Lischer and Excoffier 2012) to transform the VCF file into the “arlequin project” (ARP) format and used Arlequin to compute the site frequency spectrum (SFS) based on a total of 20,066 SNPs (Excoffier and Lischer 2010).

To infer the demographic history of the *C. beticola* population, we used fastsimcoal2 (Excoffier et al. 2013). fastsimcoal2 performs coalescent simulations to approximate the likelihood of the data given a certain demographic model and specific parameter values. Maximization of the likelihood was achieved using several Expectation Maximization iterations. To this end we generated: 1) 100,000 simulations to approximate the likelihood with high precision, 2) 40 cycles of the expectation maximization algorithm to ensure that the maximum was reached and 3) several independent replicate estimations to ensure that the global maximum likelihood was found.

We compared a set of models with different population size change scenarios. The four demographic scenarios that we compared were: 1) a recent population expansion, 2) a recent population bottleneck, 3) a bottleneck followed by a population expansion and 4) a population bottleneck followed by a second recent bottleneck (Fig. S6). To define the search range of the current effective population size, we estimated the present-day effective population size of the *C. beticola* population based on Watterson’s θ and a neutral mutation rate of 1 ×10^-8^ mutation per site per generation. As the neutral mutation rate of *Cercospora spp*. is unknown we additionally performed 100,000 simulations using the same demographic scenario with four different mutation rates (5×10^-7^, 5×10^-8^, 3×10^-8^, 1×10^-8^ mutation per site per generation) in 20 replicated runs per specified mutation rate (details of this scenario are summarized in the Supplementary Text). For our final simulations, we choose the neutral mutation rate that showed the lowest difference between the expected maximum log-likelihood and the observed log-likelihood (Table S11).

For each demographic model, we performed 100,000 simulations, 40 cycles of the expectation maximization and 50 replicate runs from different random starting values. We recorded the maximum likelihood parameter estimates that were obtained across replicate runs. Finally, we calculated the Akaike Information Criterion (AIC) and selected the model with the lowest AIC as the demographic model that best fitted the data. Parameter values were inferred in a second step by performing 100,000 simulations, 40 iterations of the expectation maximization and 100 replicate runs from different random starting values. Details on the demographic inference are summarized in the Supplementary Text.

### Genome scans for selective sweeps

Genomic scans for selective sweeps were performed by two independent approaches with the programs 1) OmegaPlus v. 3.0.3 (Alachiotis et al. 2012) and 2) RAiSD v 2.9 (Alachiotis and Pavlidis 2018). OmegaPlus is a scalable implementation of the ω statistic (Nielsen et al. 2005) that can be applied to whole-genome data. It applies a maximum likelihood framework and utilizes information on the linkage disequilibrium (LD) between SNPs. The selective sweep analysis by OmegaPlus was performed for each chromosome separately. The grid size (the number of positions for which the ω statistic is calculated) was equal to the number of variants that each chromosome contained (28,698 - 77,617 points). The minimum and maximum window sizes were set to 1,000 bp and 100,000 bp respectively. RAiSD computes the *μ* statistic, a composite evaluation test that scores genomic regions by quantifying changes in the SFS, the levels of LD, and the amount of genetic diversity along the chromosome (Alachiotis and Pavlidis 2018). We used RAiSD to detect outlier loci using VCF files of individual chromosomes providing further the length of the chromosome, the number of variant positions and a default window size of 50 kb.

To determine the significance of the identified selective sweeps we computed the ω and *μ* statistics on 10,000 datasets simulated under the best neutral demographic scenario using the program ms (Hudson 2002). Setting a significance threshold for the deviation of the ω and *μ* statistics based on simulated data sets under the best neutral demographic model allowed us to control for the effect of demographic history of the population on the SFS, LD and genetic diversity along the genome (Nielsen et al. 2005; Pavlidis et al. 2013).

### *CbCYP51* gene sequencing

*C. beticola* isolates were sampled and single-spored from the RRV region of North Dakota and Minnesota in 2019 (n=52), as described above. DNA was extracted directly from PDA cultures using a quick sodium dodecyl sulfate (SDS) lysis prep (Fran Lopez Ruiz, personal communication). The entire *CbCYP51* gene sequence (NCBI XP_023450255.1) was amplified in PCR with primers 530 and 532 from Bolton et al. (2012a), using standard conditions. Sanger sequencing of the entire PCR product was carried out using external primers 530 and 532, and internal primers 426, 349 and 566 from Bolton et al. (2012a).

### Transcription factor binding site prediction

The JASPAR CORE fungal database (www.jaspar.genereg.net/) (Fornes et al. 2020) containing experimentally-determined preferred DNA binding sites of transcription factors was queried with two different 21 bp sequences upstream of *CbCYP51*, which had the C or T SNP at 9_1452111 in the middle of the sequence: AATGTTGAAACCGTACGATGG (sensitive) and AATGTTGAAATCGTACGATGG (resistant). The 184 non-redundant matrix models from fungi were used to scan the two sequences using a relative profile score threshold of 80%, and differential results were reported.

### Codon usage assessment

Predicted coding sequences (CDS) for the 09-40 *C. beticola* reference were downloaded from NCBI (RefSeq assembly accession GCF_002742065.1) and entered into the Codon Usage tool in the Sequence Manipulation Suite (Stothard 2000) in order to calculate number and frequency of each codon type.

### Phylogenetic analysis

Ascomycete CYP51 protein sequences (including all paralogs CYP51A, B and C when present in a species) were obtained from DOE Joint Genome Institute (JGI) or NCBI protein databases along with *Saccharomyces cerevisiae* ERG11 and *Homo sapiens* CYP7A1, which was used as an outgroup. Amino acid sequence alignment was performed in MUSCLE 3.8.31 (Edgar 2004) and the alignment was refined using Gblocks 0.91b (Castresana 2002). Maximum-likelihood phylogenetic analysis was executed in PhyML 3.0 (Guindon et al. 2010) and TreeDyn 198.3 was used to render the final tree (Chevenet et al. 2006). All tools were implemented using the phylogeny.fr platform (Dereeper et al. 2008).

### Cas9 RNP-mediated gene editing

The method used was adapted from Foster et al. (2018). SgRNAs were selected using the E-CRISP program (http://www.e-crisp.org/E-CRISP/) (Table S7) and then BLASTn was performed against the *C. beticola* 09-40 reference genome to ensure the absence of off-target binding sites. Oligonucleotides for sgRNA synthesis were designed using NEB’s online tool (http://nebiocalculator.neb.com/#!/sgrna). SgRNAs were synthesized *in vitro* using the EnGen sgRNA synthesis kit, *S. pyogenes* (NEB #E3322) and were purified and concentrated into 25 μL using the RNA Clean & Concentrator-25 kit (Zymo Research #R1017). SgRNAs were complexed to 6 μg EnGen Spy Cas9-NLS (NEB #M0646) in an approximately 1:1 molar ratio for 15 min at RT. Donor DNA for introducing targeted SNPs was an 81 bp oligonucleotide with the desired single bp change in the center and 40 bp flanks of homologous sequence (Table S7). Forward and reverse complement versions of the donor oligonucleotide were complexed together at 95°C for five min prior to transformation. For co-editing, the *Hph* hygromycin resistance gene was amplified from pDAN (Friesen et al. 2006) using oligonucleotides with an additional 5’ 40 bp of homologous sequence flanking the desired locus for insertion (Table S7) and was purified and concentrated using ethanol-sodium acetate precipitation. Protoplasts were generated from cultures of benzimidazole-sensitive strain 16-1124 and DMI-sensitive strain 16-100 as described in Bolton et al. (2016). The complexed Cas9-RNP was added to 150 μL of protoplasts at 10^8^/mL concentration alongside 2 μL donor DNA (for co-editing 2 x Cas9-RNP were added at half concentration, one to create the SNP change in *CbCYP51* and the other to insert *Hph* simultaneously at another locus) and the protoplasts were incubated on ice for 30 min. The remainder of the procedure starting from PEG addition occurred as described by Liu and Friesen (2012). The only changes were in the addition of compounds for selective growth in the regeneration media agar: 10 μg/mL topsin (for E198A in beta-tubulin editing attempts) and 5-10 μg/mL tetraconazole (for *CbCYP51* editing attempts). Putative transformants growing on the surface of the agar after ~5-10 days at RT were isolated and DNA extracted to perform Sanger sequencing over the relevant intended mutation.

## Acknowledgements

The M.D. Bolton laboratory is supported by USDA project 3060-21000-044-00-D and grants from the Sugar Beet Research and Education Board of Minnesota and North Dakota, and the Beet Sugar Development Foundation. The involvement of N. Vaghefi and S. J. Pethybridge was supported by the U.S. Department of Agriculture National Institute of Food and Agriculture Hatch project NYG-625424.The authors thank Nathan Wyatt, Shaun Clare, Subidhya Shrestha and Roshan Sharma Poudel for fruitful discussions about bioinformatics, association mapping and CRISPR-Cas9 editing strategies.

