## Supplementary material for "Genome-wide association studies reveal the complex genetic architecture of DMI fungicide resistance in *Cercospora beticola*": SI

**Supplementary Information**

**Supplementary Figures**

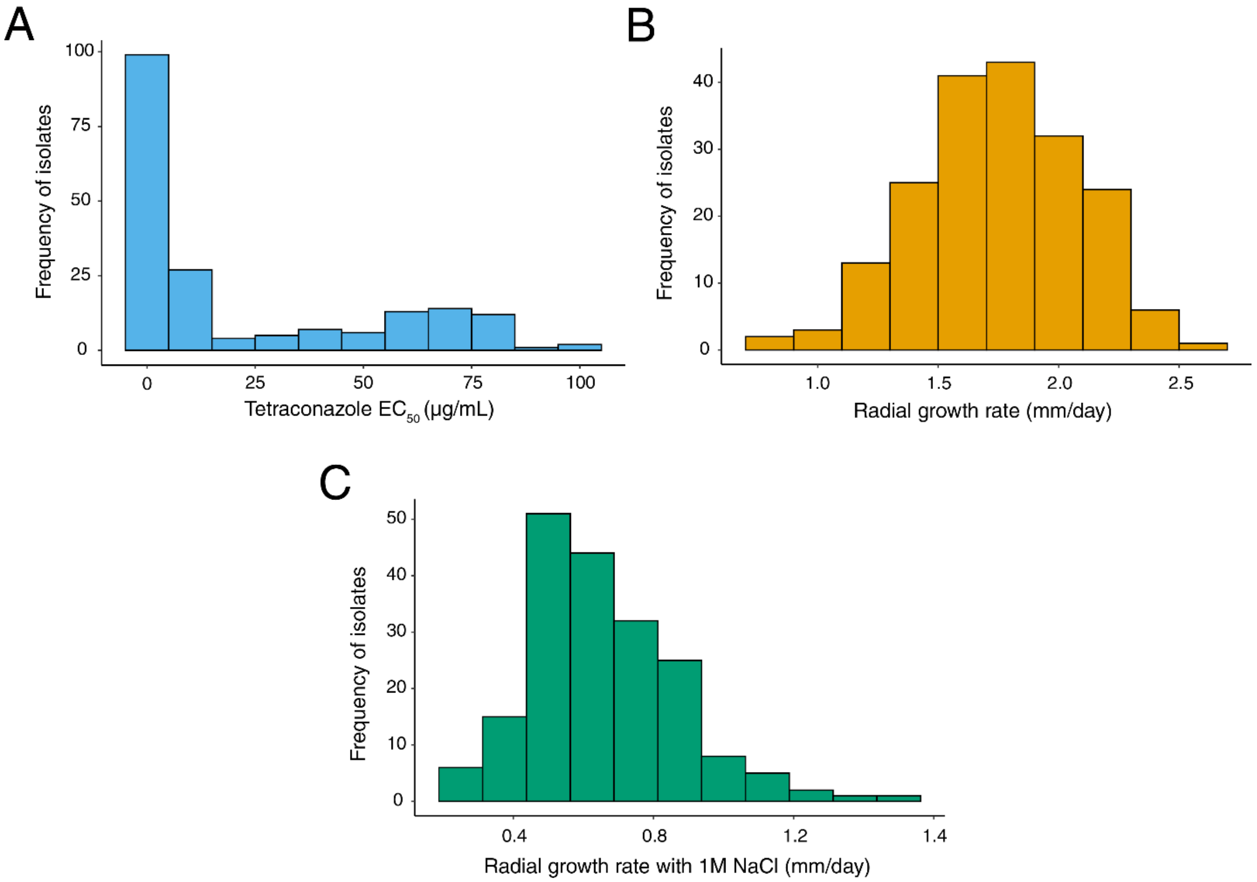


**Figure S1. Phenotypic distributions for (A) tetraconazole sensitivity ($EC_{50}$ in µg/mL), (B) radial growth rate (mm/day) and (C) radial growth under 1M NaCl salt stress (mm/day) in 190 *C. beticola* isolates.**

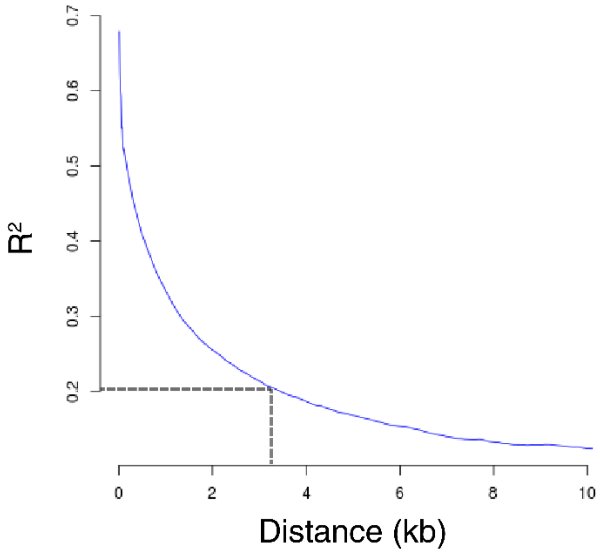


**Figure S2. Linkage disequilibrium (LD) decay in *C. beticola*.** LD decay was calculated as squared correlation of allele frequencies ($R^2$) between all pairwise combinations of markers within 10 kb of each other in the population of 190 *C. beticola* isolates. LD decays to 0.2 within 3.5 kb.

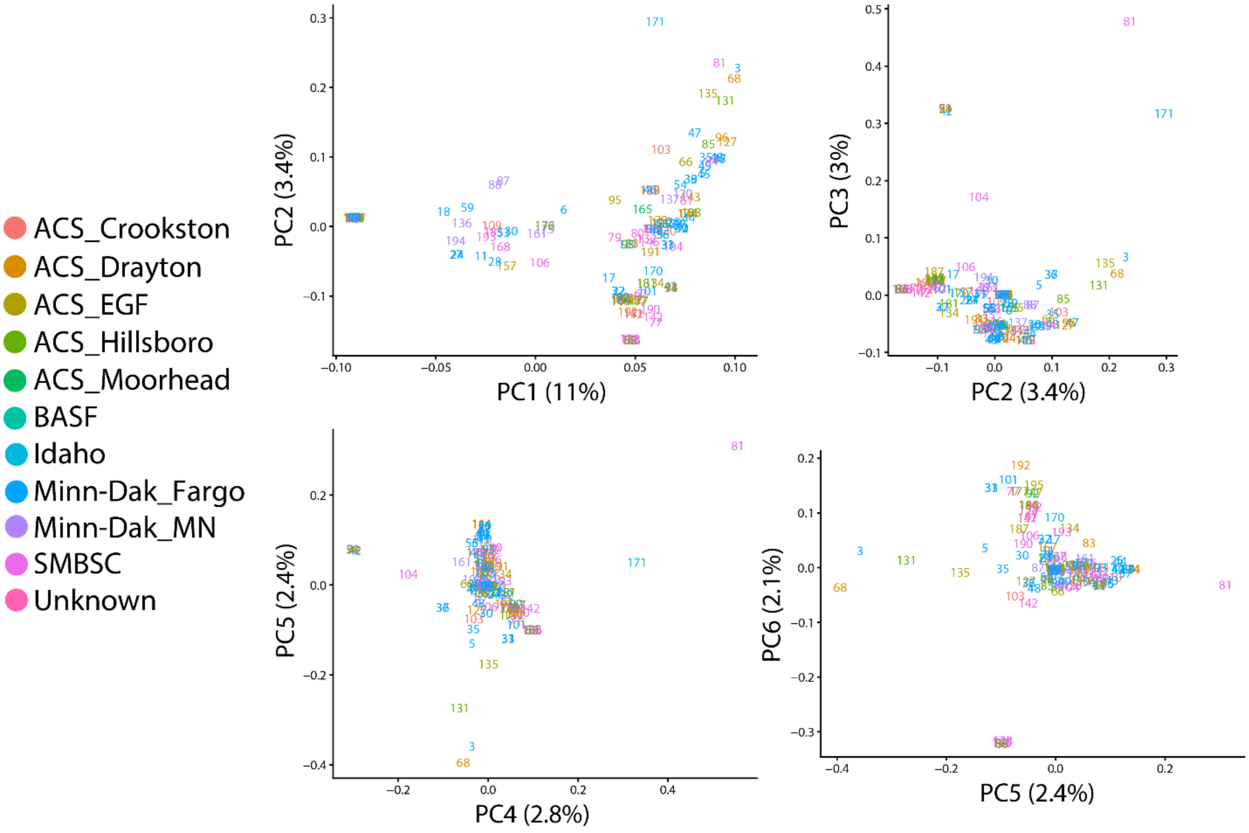


**Figure S3. Pairwise plots of the first six principle components from PCA, color-coded by sampling location.** Each location represents a different sugar beet growing area from which *C. beticola* was sampled in 2016 or 2017.

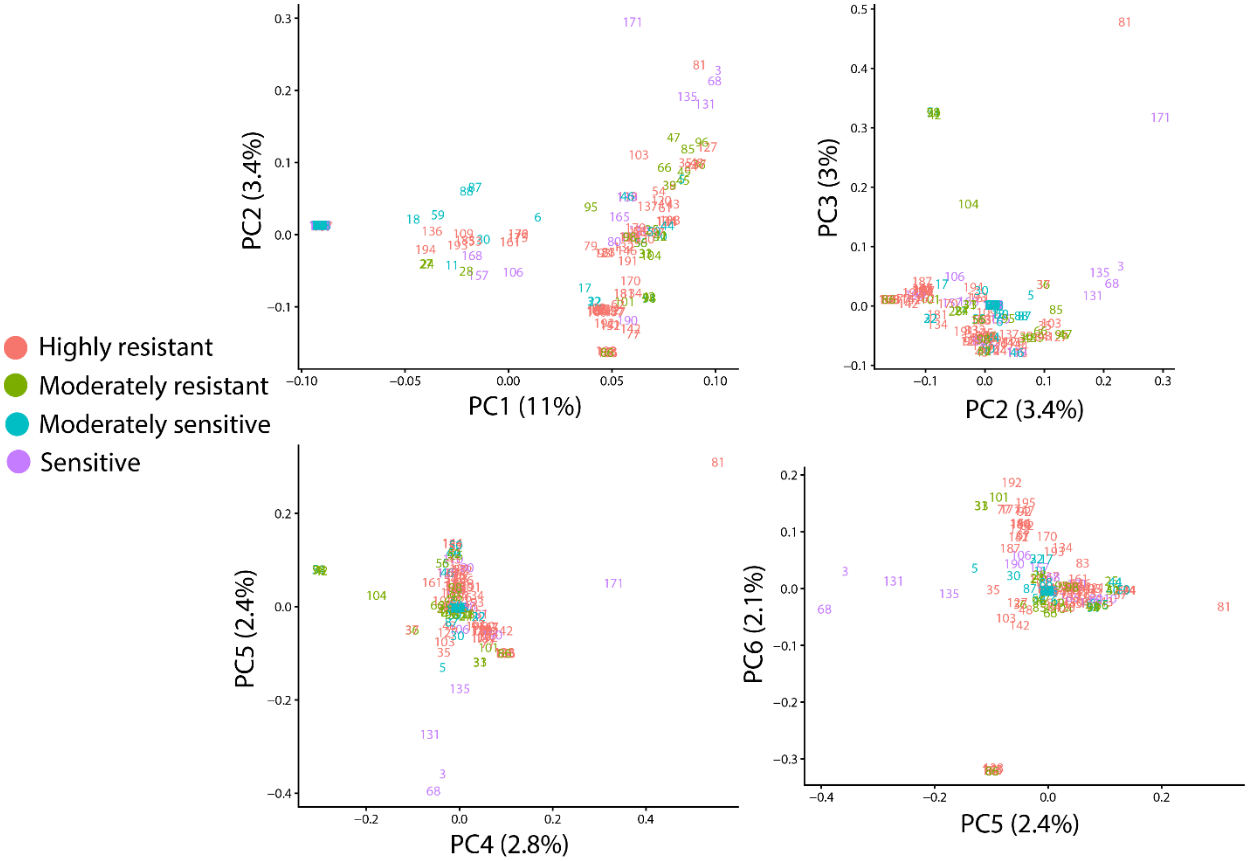


**Figures S4. Pairwise plots of the first six principle components from PCA, color-coded by tetraconazole sensitivity.** Highly resistant = isolates with $EC_{50} \geq 10$ μg/mL; Moderately resistant = isolates 1 μg/mL $\leq EC_{50} < 10$ μg/mL; Moderately sensitive = isolates with 0.1 μg/mL $\leq EC_{50} < 1$ μg/mL; Sensitive = isolates with $EC_{50} < 0.1$ μg/mL.

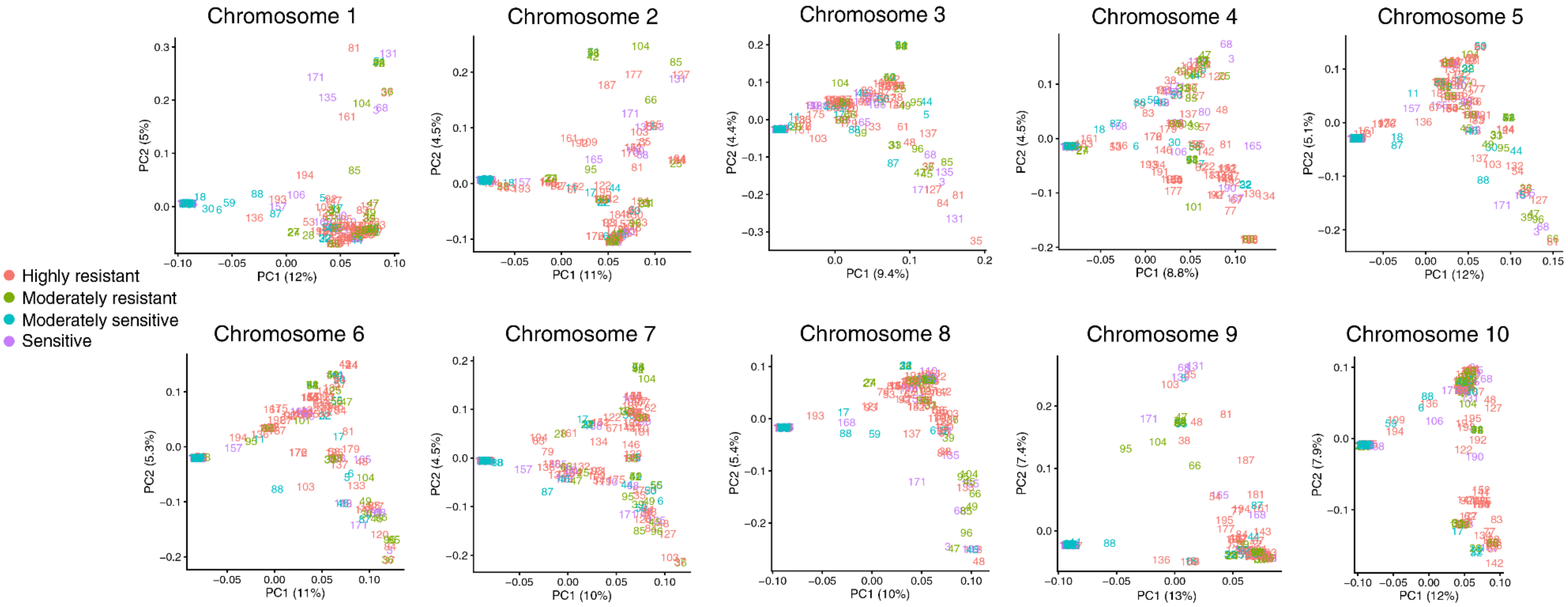


**Figure S5. Pairwise plots of the first two principle components from PCAs performed individually for LD-pruned SNPs on chromosomes one to ten, color-coded by tetraconazole sensitivity.** Chromosome one = 6,244 SNPs; chromosome two = 6,586 SNPs; chromosome three = 6,249 SNPs; chromosome four = 5,473 SNPs; chromosome five = 4,376 SNPs; chromosome six = 4,949 SNPs; chromosome seven = 5,864 SNPs; chromosome eight = 3,651 SNPs; chromosome nine = 1,507 SNPs; chromosome ten = 1,918 SNPs. Highly resistant = isolates with $EC_{50} \geq 10$ µg/mL; Moderately resistant = isolates 1 µg/mL $\leq EC_{50} < 10$ µg/mL; Moderately sensitive = isolates with 0.1 µg/mL $\leq EC_{50} < 1$ µg/mL; Sensitive = isolates with $EC_{50} < 0.1$ µg/mL.

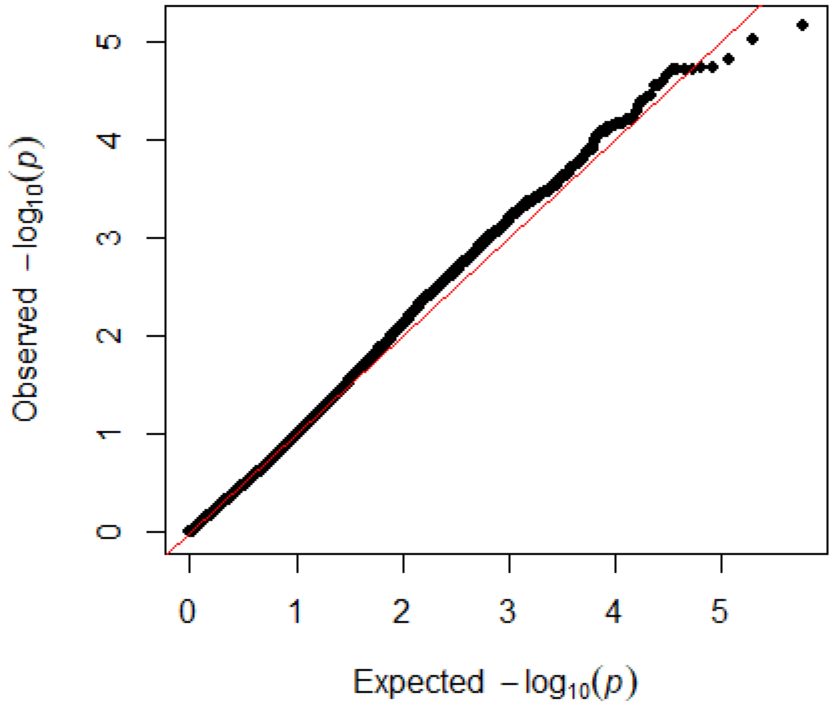


**Figure S6. Quantile-Quantile plot for genome-wide association of tetraconazole sensitivity.**

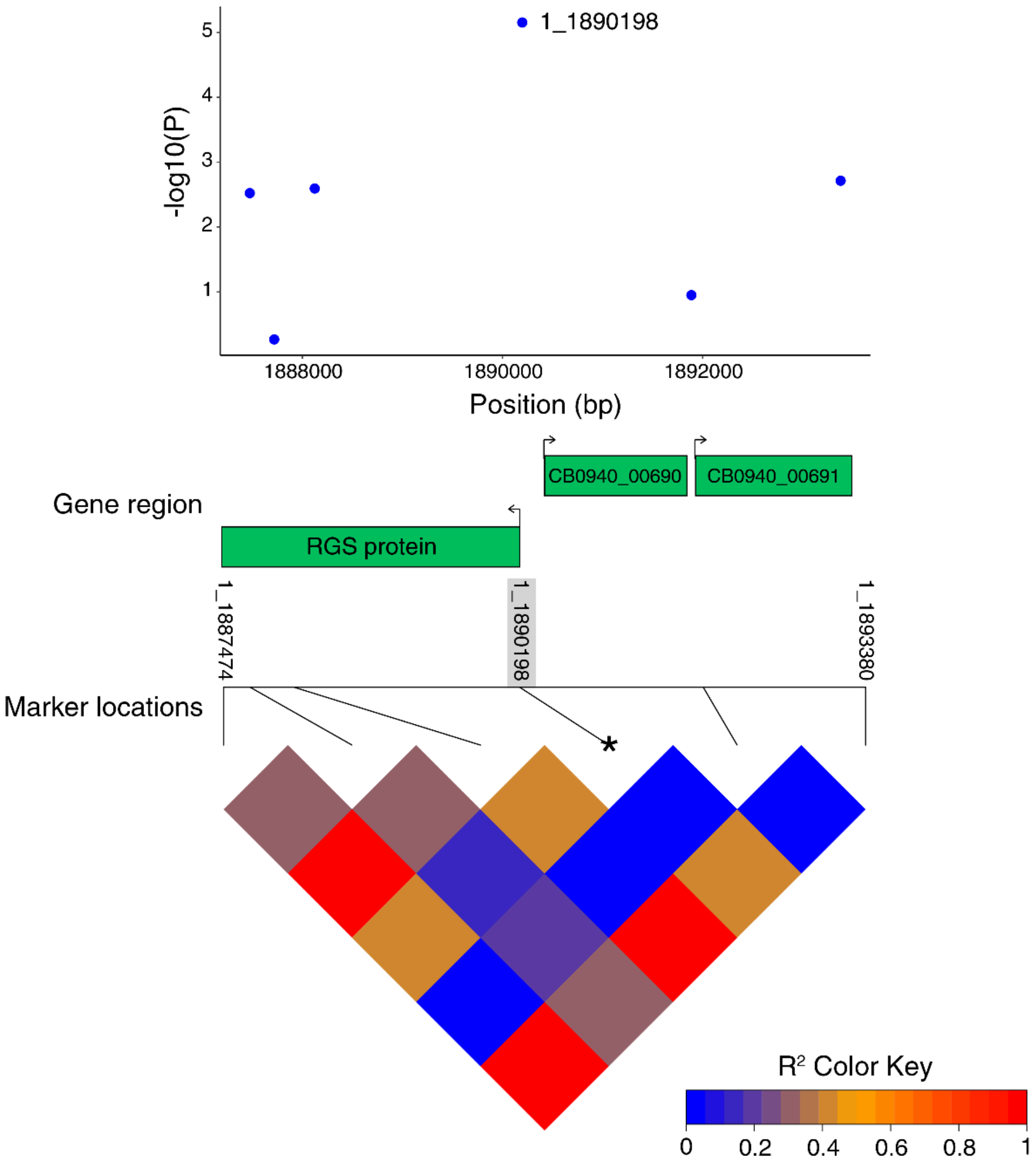


**Figure S7. Genomic context of the 1_1890198 marker in RGS gene CB0940_00689 significantly associated with tetraconazole sensitivity.** Top panel: Scatter plot of GWAS *P*-values centered on the 1_1890198 marker ±3 kb (six markers in total). Predicted genes CB0940_00689 (RGS protein), CB0940_00690 and CB0940_00691 are shown in green at approximate locations within this 5.9 kb region. Bottom panel: an LD heat map showing the pairwise LD values ($R^2$) between all six markers within the 5.9 kb region surrounding 1_1890198. The 1_1890198 marker is highlighted in grey and indicated with a star.

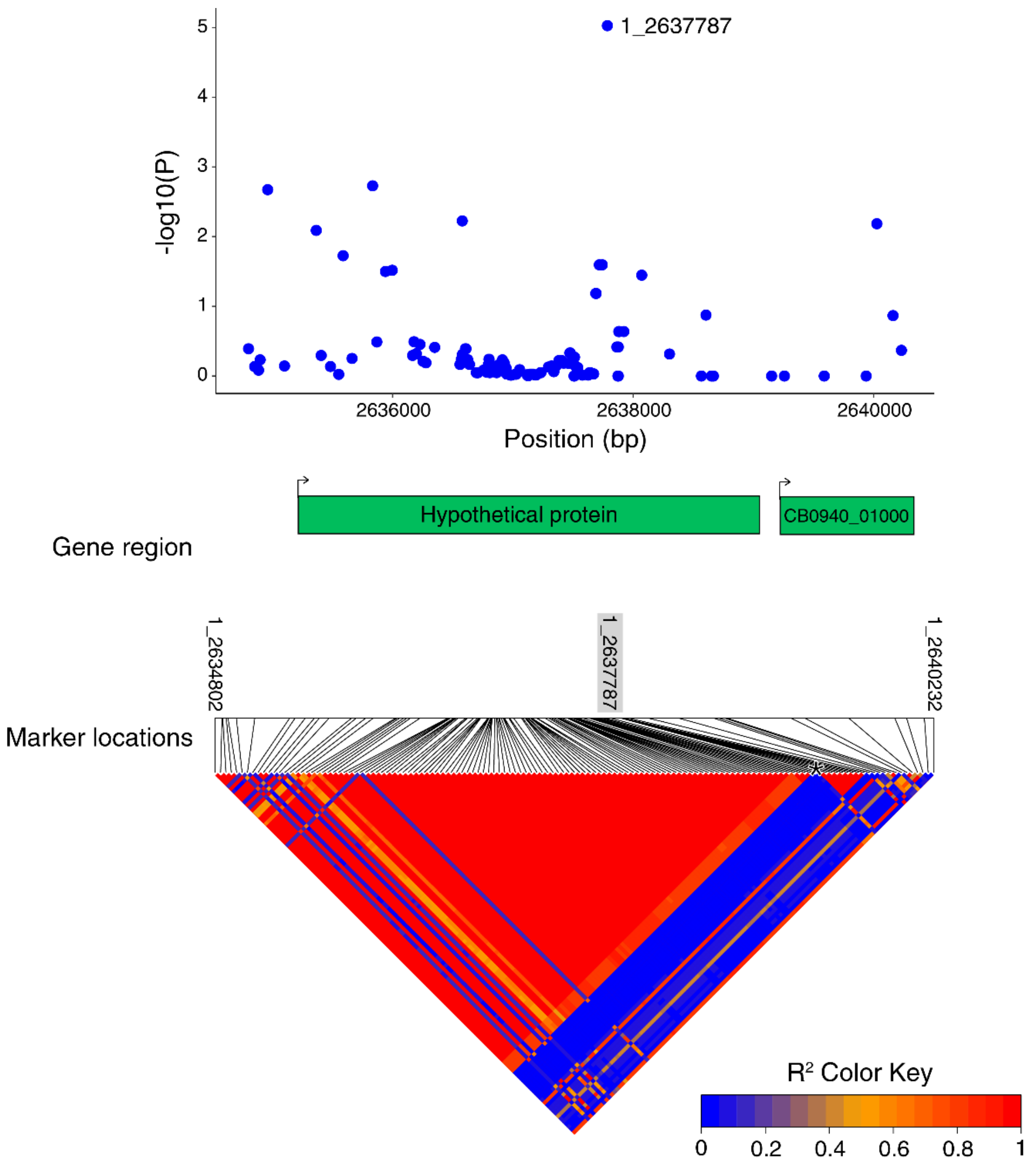


**Figure S8. Genomic context of the 1_2637787 marker in gene CB0940_0999 (hypothetical protein) significantly associated with tetraconazole sensitivity.** Top panel: Scatter plot of GWAS *P*-values centered on the 1_2637787 SNP ±3 kb (135 markers in total). Predicted genes CB0940_00999 (hypothetical protein) and CB0940_01000 are shown in green at approximate locations within this 5.4 kb region. Bottom panel: an LD heat map showing the pairwise LD values ($R^2$) between all 135 markers within the 5.4 kb region surrounding 1_2637787. The 1_2637787 marker is highlighted in grey and indicated with a star.

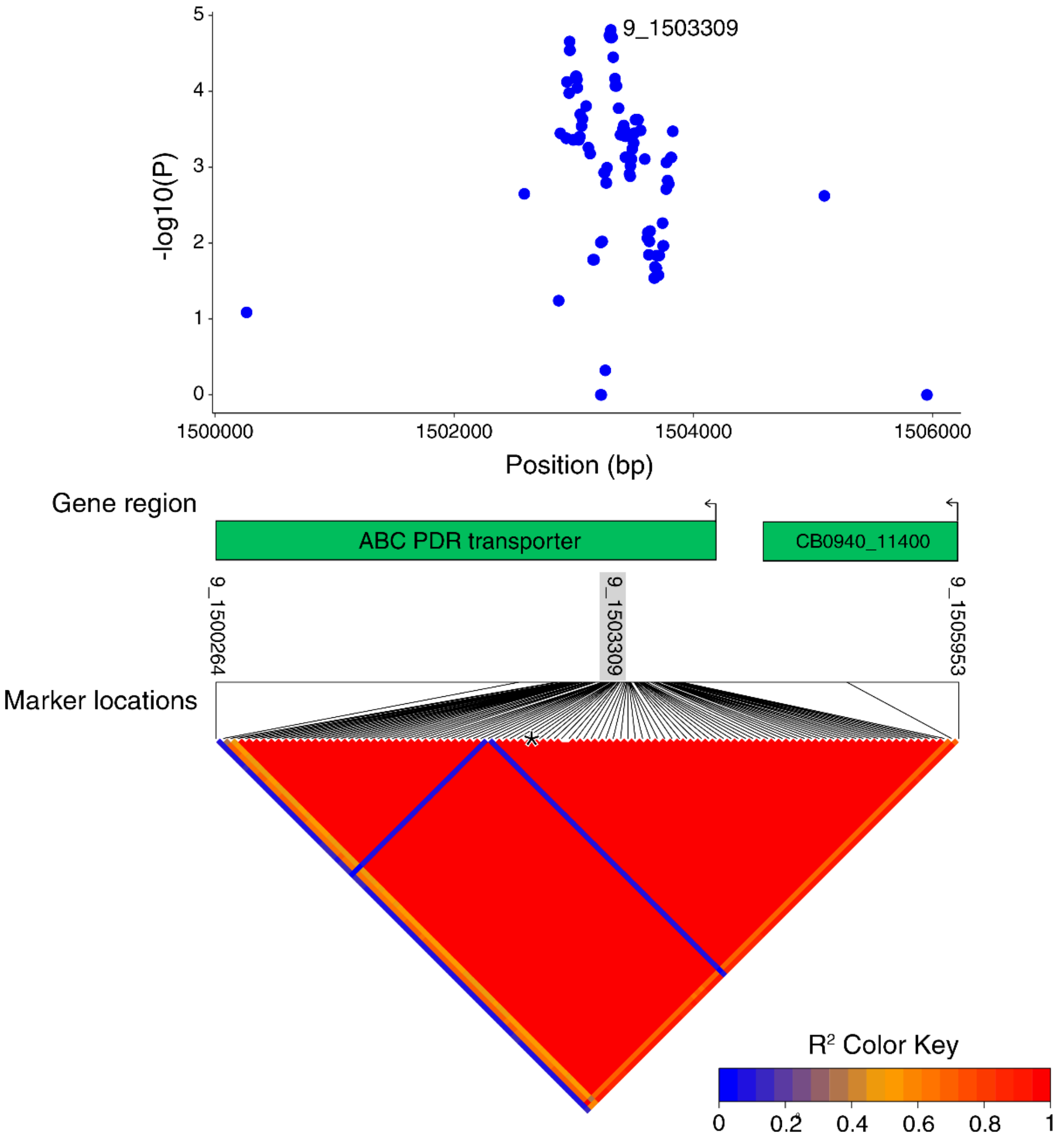


**Figure S9. Genomic context of the 9_1503309 marker in gene CB0940_11399 (ABC PDR transporter) significantly associated with tetraconazole sensitivity.** Top panel: Scatter plot of GWAS *P*-values centered on the 9_1503309 SNP ±3 kb (102 markers in total). Predicted genes CB0940_11399 (ABC PDR transporter) and CB0940_11400 are shown in green at approximate locations within this 5.7 kb region. Bottom panel: an LD heat map showing the pairwise LD values ($R^2$) between all 102 markers within the 5.7 kb region surrounding 9_1503309. The 9_1503309 marker is highlighted in grey and indicated with a star.

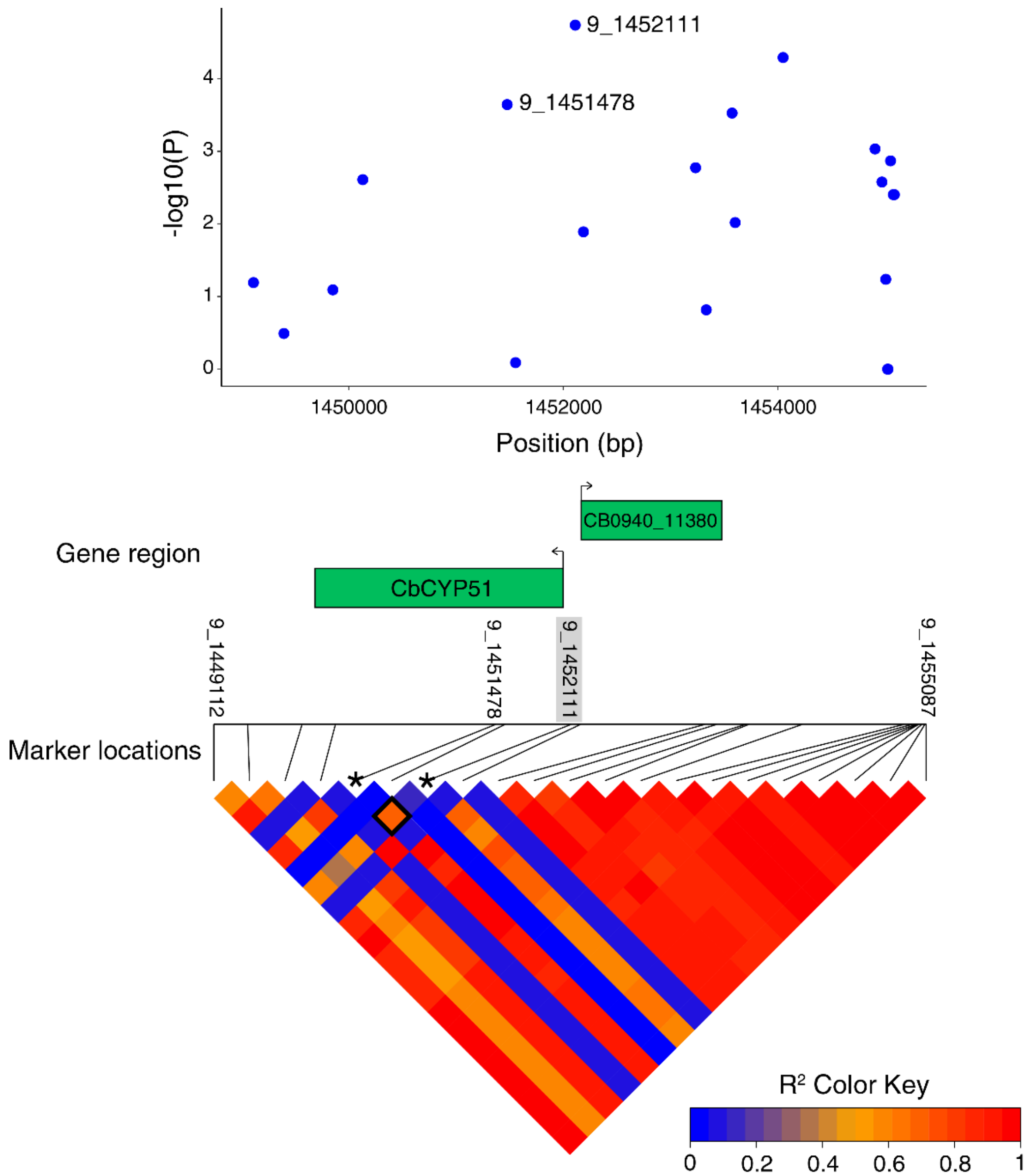


**Figure S10. Genomic context of the 9_1452111 marker upstream of gene CB0940_11379 (*CbCYP51*) significantly associated with tetraconazole sensitivity.** Top panel: Scatter plot of GWAS *P*-values centered on the 9_1452111 SNP ±3 kb (21 markers in total). Predicted genes CB0940_11379 (*CbCYP51*) and CB0940_11380 are shown in green at approximate locations within this 6 kb region. Bottom panel: an LD heat map showing the pairwise LD values ($R^2$) between all 21 markers within the 6 kb region surrounding 9_1452111. The 9_1452111 marker is highlighted in grey and indicated with a star. The 9_1451478 marker (synonymous mutation

E170) is also indicated with a star, and the square representing pairwise LD ($R^2$) with 9_1452111 is outlined in black.

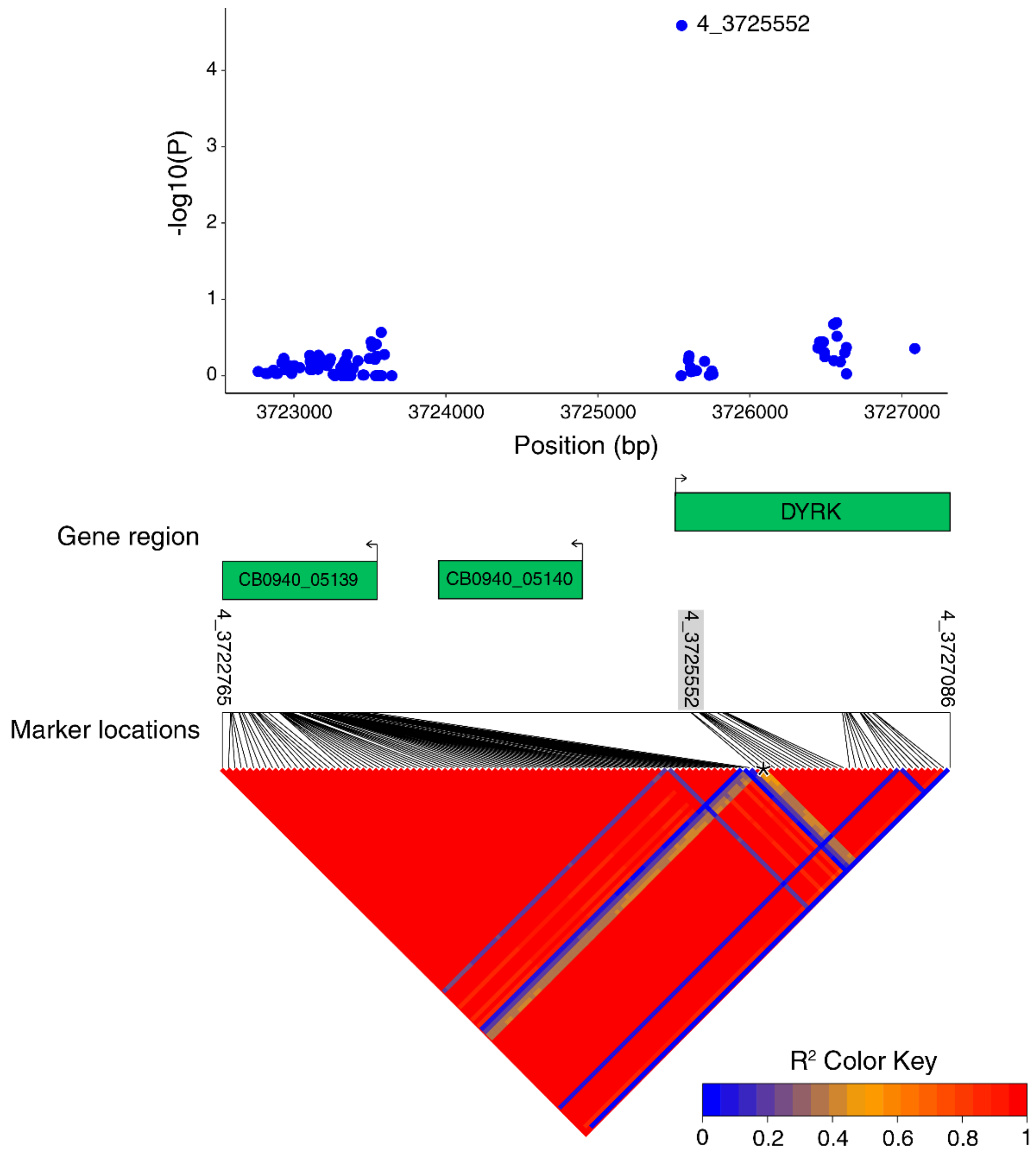


**Figure S11. Genomic context of the 4_3725552 marker in gene CB0940_05141 (DYRK) significantly associated with tetraconazole sensitivity.** Top panel: Scatter plot of GWAS *P*-values centered on the 4_3725552 SNP ±3 kb (117 markers in total). Predicted genes CB0940_05139, CB0940_05140 and CB0940_05141 (DYRK) are shown in green at approximate locations within this 4.3 kb region. Bottom panel: an LD heat map showing the pairwise LD values ($R^2$) between all 117 markers within the 4.3 kb region surrounding 4_3725552. The 4_3725552 marker is highlighted in grey and indicated with a star.

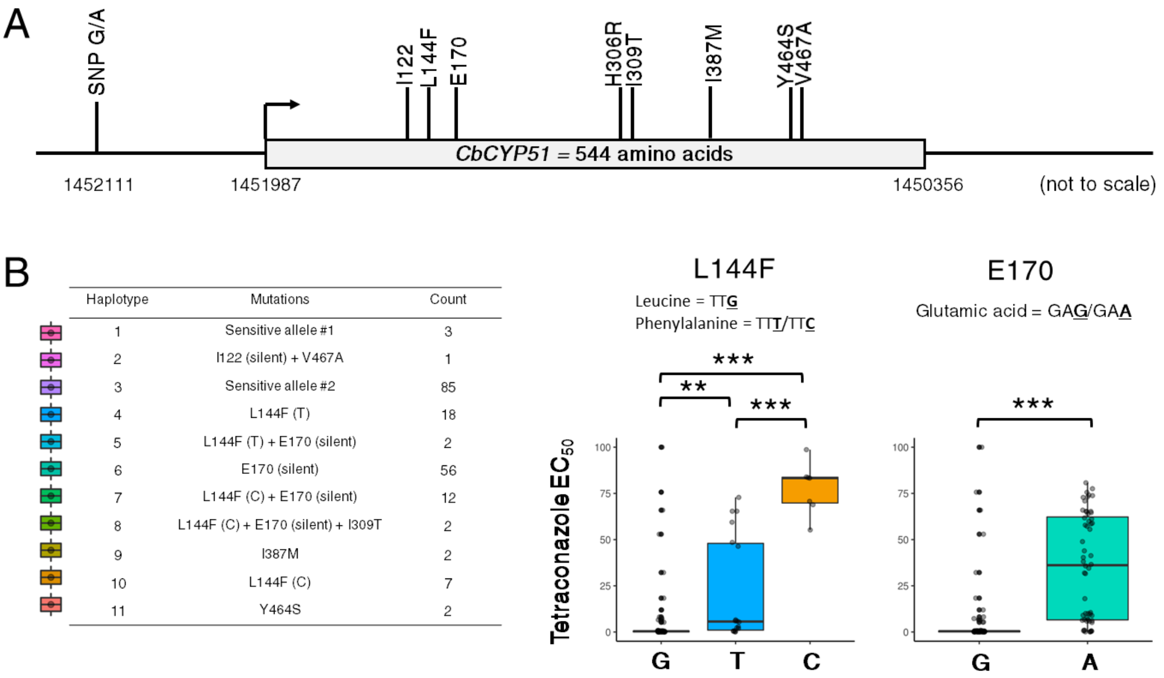


**Figure S12.** ***CbCYP51*** **gene model in** ***C. beticola*** **and the effects of L144F and E170 mutations on tetraconazole $EC_{50}$ value.** A) *CbCYP51* gene model displaying all coding mutations found in this study, and a single upstream mutation. B) Isolates with single CbCYP51 mutations were used to compare the effects of L144F codons; phenyalanine codons TTT and TTC both gave significantly higher $EC_{50}$ values than leucine codon TTG ($P < 0.01$), but codon TTC also gave significantly higher $EC_{50}$ values than codon TTT ($P < 0.001$). Isolates with single CbCYP51 mutations were also used to compare the effects of E170; glutamic acid codon GAA gave significantly higher $EC_{50}$ values than codon GAG ($P < 0.001$).

| Haplotype | Mutations | Count |
|---|---|---|
| 1 | Sensitive allele #1 | 1 |
| 3 | Sensitive allele #2 | 9 |
| 4 | L144F (T) | 11 |
| 6 | E170 (silent) | 12 |
| 7 | L144F (C) + E170 (silent) | 11 |
| 8 | L144F (C) + E170 (silent) + I309T | 2 |
| 10 | L144F (C) | 4 |
| 11 | Y464S | 1 |
| 12 | L144F (T) + H306R | 1 |

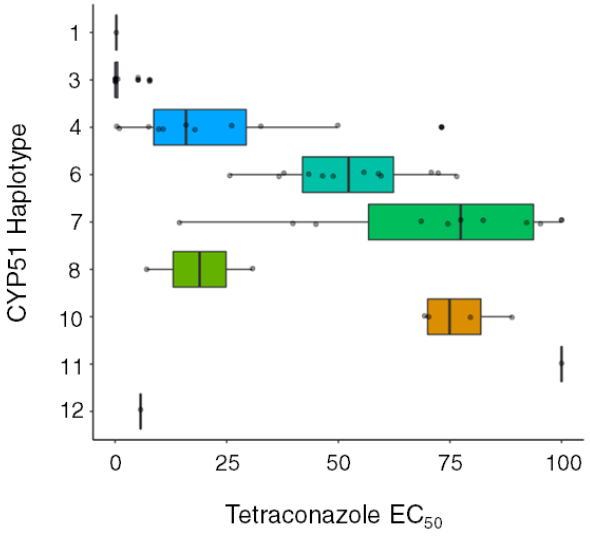


**Figure S13. Effect of *CbCYP51* haplotypes on tetraconazole sensitivity in 2019 *C. beticola* strains**. The left panel displays 9 different *CbCYP51* coding sequence haplotypes found in 52 *C. beticola* isolates from 2019. The haplotypes have consistent numbering with the 190 *C. beticola* isolates from 2016 and 2017, displayed in Figure 4. Mutations were identified as compared to the most common sensitive Haplotype (#3). The right panel displays box and whiskers plots for tetraconazole $EC_{50}$ values for each *CbCYP51* haplotype.

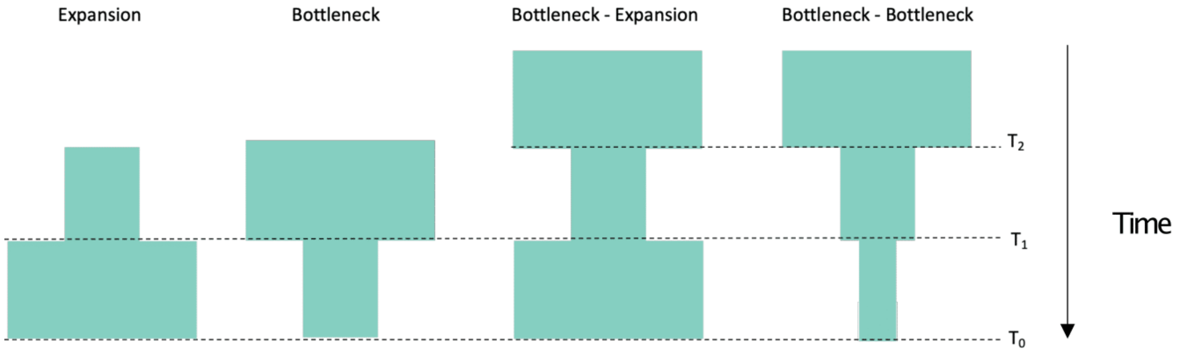


**Figure S14. Illustration of the four demographic models that were compared in the demography inference of *C beticola*.** The effective population size is represented by the width of the figure. T0 represents the present time, T1 represents the time of the most recent population size change and T2 a more ancient population size change.

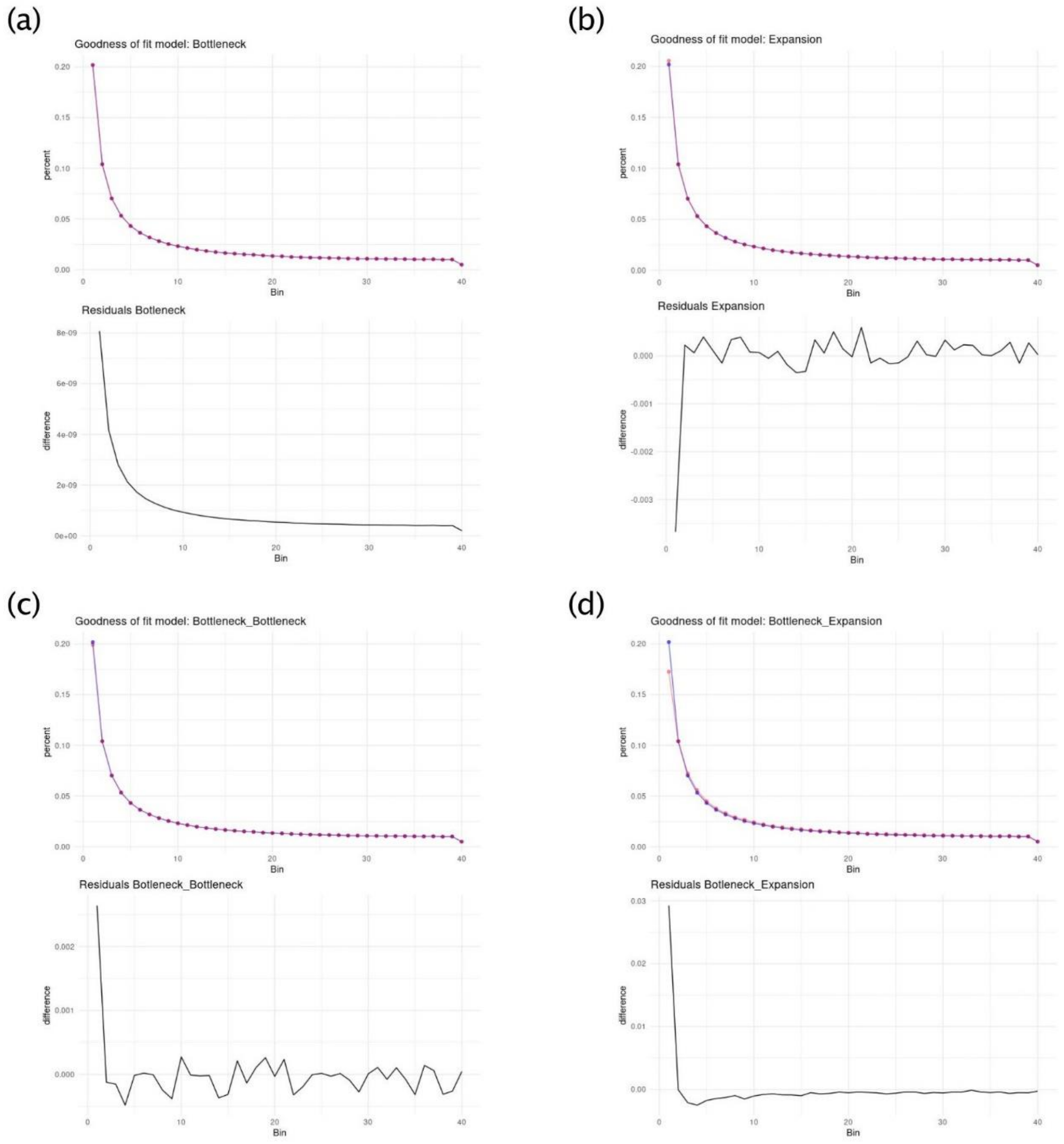


**Figure S15. Comparison of the observed site frequency spectrum (SFS) and the expected SFS for the four demographic scenarios: (a) bottleneck, (b) expansion, (c) bottleneck – bottleneck, (d) bottleneck - expansion and their residuals.**

| Haplotype | Mutations | Count |
|---|---|---|
| 1 | 999bp deletion | 7 |
| 2 | Reference 09-40 | 120 |
| 3 | G57A | 50 |
| 4 | G57A + L224 | 5 |
| 5 | G57A + 7 more SNPs | 8 |

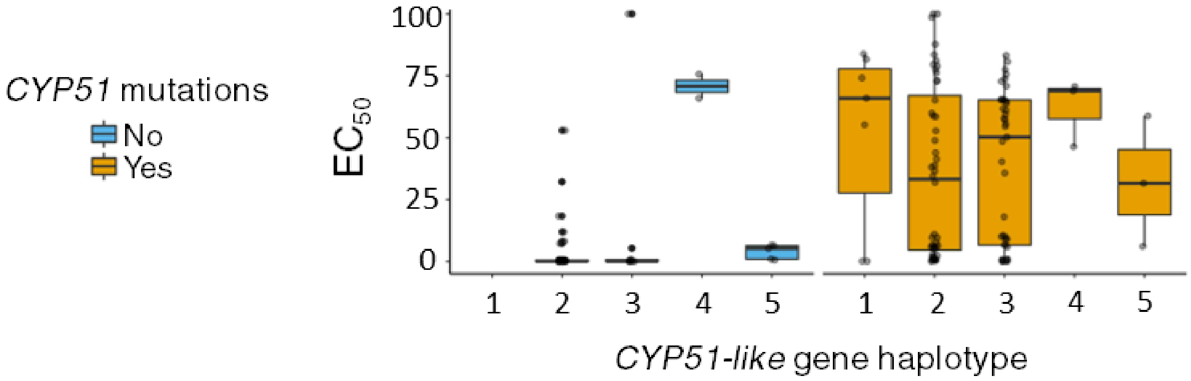


**Figure S16. Effects of *CbCYP51-like* haplotypes on tetraconazole sensitivity.** The table shows the five different *CbCYP51* coding sequence haplotypes found in our *C. beticola* population with the respective number of isolates with each haplotype. Below are box and whiskers plots for tetraconazole $EC_{50}$ values for each *CbCYP51-like* haplotype, grouped by the presence or absence of *CbCYP51* mutations.

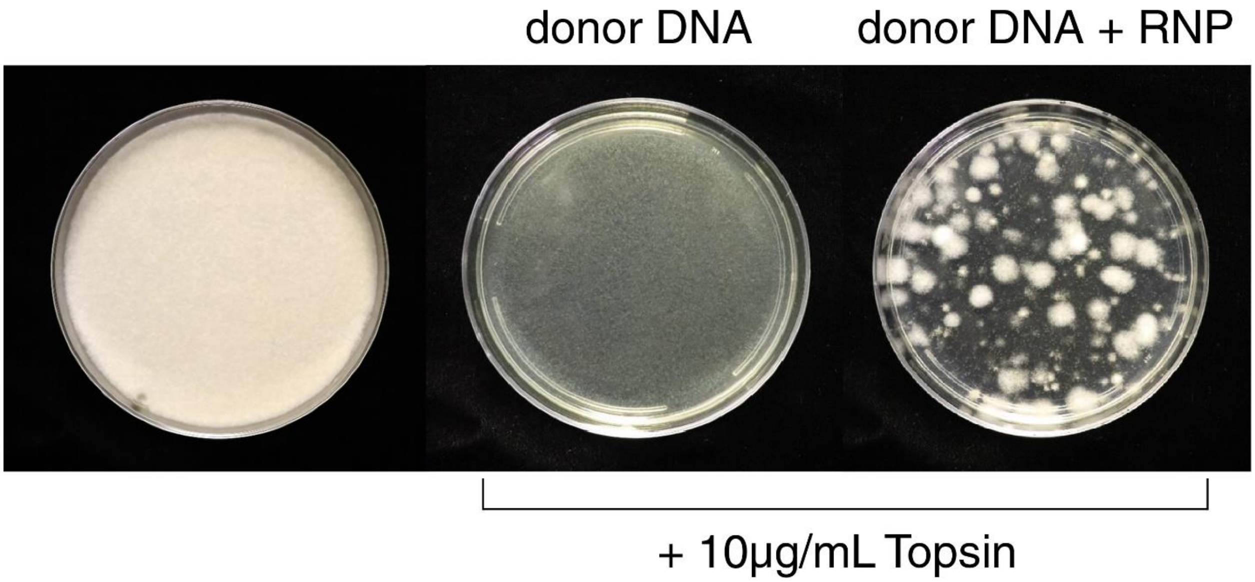


**Figure S17. Successful use of Cas9-RNP editing in *C. beticola* to introduce the E198A mutation in beta-tubulin conferring resistance to benzimidazole fungicides.** From left to right: protoplasted isolate 16-1124 grown on regeneration media without topsin, as a positive control; protoplasted isolate 16-1124 transformed with just the donor DNA and grown on regeneration media with 10 µg/mL topsin, as a negative control; protoplasted isolate 16-1124 transformed with Cas9-RNP and donor DNA to introduce E198A mutation into beta-tubulin with regeneration on media containing 10 µg/mL topsin.

**Supplementary Tables**

**Table S1. Isolates used in this study.**

| Isolate number[a] | Isolate name[b] | Sampling location[c] | Tetraconazole EC50[d] | Radial growth rate (mm/day)[e] | Radial growth rate with 1M NaCl (mm/day)[f] |
|---|---|---|---|---|---|
| 1 | 16-F1 | Minn-Dak_Fargo | 0.569 | 2.182 | 0.595 |
| 2 | 16-F2 | Minn-Dak_Fargo | 0.542 | 1.813 | 0.497 |
| 3 | 16-F3 | Minn-Dak_Fargo | 0.083 | 1.791 | 1.001 |
| 4 | 16-F4 | Minn-Dak_Fargo | 0.521 | 1.807 | 0.840 |
| 5 | 16-F5 | Minn-Dak_Fargo | 0.116 | 2.059 | 0.622 |
| 6 | 16-F6 | Minn-Dak_Fargo | 0.585 | 1.680 | 0.571 |
| 7 | 16-F7 | Minn-Dak_Fargo | 0.567 | 2.138 | 0.375 |
| 9 | 16-F9 | Minn-Dak_Fargo | 0.553 | 2.022 | 0.540 |
| 11 | 16-F11 | Minn-Dak_Fargo | 0.546 | 1.727 | 0.573 |
| 12 | 16-F12 | Minn-Dak_Fargo | 0.503 | 1.853 | 0.558 |
| 13 | 16-F13 | Minn-Dak_Fargo | 0.354 | 2.029 | 0.824 |
| 14 | 16-F14 | Minn-Dak_Fargo | 0.579 | 1.631 | 0.593 |
| 15 | 16-F15 | Minn-Dak_Fargo | 0.528 | 1.310 | 0.555 |
| 16 | 16-F16 | Minn-Dak_Fargo | 0.063 | 1.582 | 0.804 |
| 17 | 16-F17 | Minn-Dak_Fargo | 0.637 | 1.824 | 0.726 |
| 18 | 16-F18 | Minn-Dak_Fargo | 0.560 | 1.764 | 0.564 |
| 19 | 16-F19 | Minn-Dak_Fargo | 0.428 | 1.339 | 0.524 |
| 20 | 16-F20 | Minn-Dak_Fargo | 0.509 | 0.879 | 0.467 |
| 21 | 16-F21 | Minn-Dak_Fargo | 0.517 | 1.600 | 0.415 |
| 22 | 16-F22 | Minn-Dak_Fargo | 0.740 | 2.166 | 1.061 |
| 23 | 16-F23 | Minn-Dak_Fargo | 38.107 | 1.889 | 0.377 |
| 24 | 16-F24 | Minn-Dak_Fargo | 1.929 | 1.761 | 0.584 |
| 25 | 16-F25 | Minn-Dak_Fargo | 1.741 | 1.676 | 0.578 |
| 26 | 16-F26 | Minn-Dak_Fargo | 0.503 | 1.950 | 0.473 |
| 27 | 16-F27 | Minn-Dak_Fargo | 5.441 | 1.838 | 0.626 |

| | | | | | |
|---|---|---|---|---|---|
| 28 | 16-F28 | Minn-Dak_Fargo | 2.275 | 1.961 | 0.539 |
| 29 | 16-F29 | Minn-Dak_Fargo | 0.561 | 1.302 | 0.500 |
| 30 | 16-F30 | Minn-Dak_Fargo | 0.585 | 1.651 | 0.592 |
| 31 | 16-F31 | Minn-Dak_Fargo | 2.914 | 1.404 | 0.686 |
| 32 | 16-F32 | Minn-Dak_Fargo | 0.443 | 1.687 | 0.677 |
| 33 | 16-F33 | Minn-Dak_Fargo | 6.021 | 1.265 | 0.747 |
| 34 | 16-F34 | Minn-Dak_Fargo | 8.158 | 1.909 | 0.529 |
| 35 | 16-F35 | Minn-Dak_Fargo | 36.549 | 1.652 | 0.850 |
| 36 | 16-F36 | Minn-Dak_Fargo | 6.070 | 2.114 | 0.881 |
| 37 | 16-F37 | Minn-Dak_Fargo | 31.584 | 2.129 | 0.670 |
| 38 | 16-F38 | Minn-Dak_Fargo | 43.903 | 1.863 | 0.636 |
| 39 | 16-F39 | Minn-Dak_Fargo | 8.921 | 2.106 | 0.735 |
| 40 | 16-F40 | Minn-Dak_Fargo | 0.598 | 2.036 | 0.464 |
| 41 | 16-F41 | Minn-Dak_Fargo | 9.596 | 1.274 | 0.708 |
| 42 | 16-F42 | Minn-Dak_Fargo | 5.276 | 2.038 | 0.887 |
| 43 | 16-F43 | Minn-Dak_Fargo | 10.537 | 1.502 | 0.639 |
| 44 | 16-F44 | Minn-Dak_Fargo | 0.988 | 1.550 | 0.275 |
| 45 | 16-F45 | Minn-Dak_Fargo | 9.708 | 1.501 | 0.706 |
| 46 | 16-F46 | Minn-Dak_Fargo | 0.932 | 1.404 | 0.596 |
| 47 | 16-F47 | Minn-Dak_Fargo | 5.120 | 2.524 | 0.819 |
| 48 | 16-F48 | Minn-Dak_Fargo | 61.118 | 1.168 | 0.735 |
| 49 | 16-F49 | Minn-Dak_Fargo | 5.800 | 2.286 | 0.656 |
| 50 | 16-F50 | Minn-Dak_Fargo | 0.964 | 1.889 | 0.578 |
| 51 | 16-F51 | Minn-Dak_Fargo | 0.955 | 1.456 | 0.786 |
| 52 | 16-F52 | Minn-Dak_Fargo | 7.661 | 2.158 | 0.752 |
| 53 | 16-F53 | Minn-Dak_Fargo | 10.000 | 1.150 | 0.526 |
| 54 | 16-F54 | Minn-Dak_Fargo | 41.295 | 2.284 | 0.841 |
| 55 | 16-F55 | Minn-Dak_Fargo | 0.710 | 1.046 | 0.431 |
| 56 | 16-F56 | Minn-Dak_Fargo | 5.367 | 2.354 | 0.649 |
| 57 | 16-F57 | Minn-Dak_Fargo | 80.749 | 0.739 | 0.514 |
| 58 | 16-F58 | Minn-Dak_Fargo | 61.722 | 1.308 | 0.748 |

| | | | | | |
|---|---|---|---|---|---|
| 59 | 16-F59 | Minn-Dak_Fargo | 0.650 | 1.668 | 0.480 |
| 61 | 16-16 | SMBSC | 31.945 | 1.985 | 0.719 |
| 62 | 16-45 | SMBSC | 19.361 | 2.264 | 0.604 |
| 63 | 16-46 | SMBSC | 59.412 | 1.785 | 0.555 |
| 64 | 16-62 | Unknown | 12.331 | 1.988 | 0.500 |
| 65 | 16-92 | SMBSC | 64.560 | 1.737 | 0.819 |
| 66 | 16-131 | ACS_EGF | 6.653 | 1.532 | 0.407 |
| 67 | 16-175 | SMBSC | 83.875 | 2.181 | 1.453 |
| 68 | 16-339 | ACS_Drayton | 0.067 | 1.451 | 0.273 |
| 69 | 16-591 | ACS_EGF | 0.395 | 1.945 | 0.461 |
| 71 | 16-665 | Minn-Dak_MN | 0.371 | 1.781 | 1.224 |
| 72 | 16-735 | Minn-Dak_MN | 0.403 | 1.684 | 0.579 |
| 73 | 16-781 | Minn-Dak_MN | 0.205 | 1.609 | 0.825 |
| 74 | 16-787 | Minn-Dak_MN | 0.455 | 1.384 | 0.524 |
| 75 | 16-805 | ACS_Moorhead | 0.404 | 1.166 | 0.722 |
| 76 | 16-810 | ACS_Moorhead | 0.400 | 2.242 | 0.784 |
| 77 | 16-890 | SMBSC | 55.224 | 2.336 | 1.220 |
| 78 | 16-1148 | ACS_Crookston | 6.362 | 1.653 | 0.813 |
| 79 | 16-26 | SMBSC | 34.512 | 1.224 | 0.508 |
| 80 | 16-33 | SMBSC | 0.094 | 1.629 | 0.551 |
| 81 | 16-72 | SMBSC | 29.821 | 1.859 | 0.522 |
| 82 | 16-93 | SMBSC | 65.370 | 1.711 | 1.176 |
| 83 | 16-317 | ACS_Drayton | 77.542 | 1.724 | 0.888 |
| 84 | 16-414 | SMBSC | 65.161 | 1.130 | 0.634 |
| 85 | 16-498 | ACS_Hillsboro | 6.622 | 1.633 | 0.783 |
| 86 | 16-523 | ACS_Hillsboro | 5.889 | 2.003 | 0.773 |
| 87 | 16-666 | Minn-Dak_MN | 0.432 | 1.875 | 0.623 |
| 88 | 16-704 | Minn-Dak_MN | 0.415 | 1.681 | 0.378 |
| 89 | 16-716 | Minn-Dak_MN | 0.451 | 1.912 | 0.527 |
| 90 | 16-722 | Minn-Dak_MN | 0.436 | 1.560 | 0.799 |
| 91 | 16-724 | Minn-Dak_MN | 0.412 | 1.342 | 0.636 |

| | | | | | |
|---|---|---|---|---|---|
| 92 | 16-790 | ACS_Moorhead | 83.546 | 2.211 | 0.818 |
| 93 | 16-819 | ACS_Moorhead | 63.830 | 1.541 | 0.719 |
| 94 | 16-868 | ACS_EGF | 6.579 | 1.507 | 0.870 |
| 95 | 16-1105 | ACS_EGF | 5.841 | 1.877 | 1.315 |
| 96 | 16-1169 | ACS_Drayton | 6.302 | 1.147 | 0.594 |
| 97 | 16-F61 | Minn-Dak_Fargo | 0.46 | 1.776 | 0.780 |
| 98 | 16-F62 | Minn-Dak_Fargo | 9.03 | 1.404 | 0.826 |
| 99 | 16-F63 | Minn-Dak_Fargo | 0.55 | 2.056 | 0.334 |
| 100 | 16-F64 | Minn-Dak_Fargo | 0.52 | 1.275 | 0.406 |
| 101 | 16-F65 | Minn-Dak_Fargo | 6.57 | 1.829 | 0.606 |
| 102 | 16-100 | SMBSC | 0.08 | 1.068 | 0.457 |
| 103 | 16-1022 | ACS_Crookston | 27.639 | 1.342 | 1.127 |
| 104 | 16-1083 | SMBSC | 7.258 | 2.241 | 0.898 |
| 105 | 16-1110 | ACS_EGF | 0.072 | 1.894 | 0.827 |
| 106 | 16-1124 | SMBSC | 0.081 | 1.949 | 0.622 |
| 107 | 16-1129 | SMBSC | 18.330 | 1.380 | 0.504 |
| 108 | 16-1138 | ACS_Crookston | 0.07 | 1.561 | 0.505 |
| 109 | 16-1151 | ACS_Crookston | 11.00 | 1.599 | 0.805 |
| 110 | 16-1155 | ACS_Crookston | 0.073 | 1.821 | 0.603 |
| 111 | 16-1211 | ACS_Drayton | 0.076 | 1.629 | 0.499 |
| 112 | 16-1214 | ACS_Drayton | 0.076 | 1.180 | 0.434 |
| 114 | 16-123 | BASF | 0.095 | 1.378 | 0.675 |
| 115 | 16-141 | ACS_Moorhead | 0.09 | 1.687 | 0.483 |
| 116 | 16-163 | ACS_Moorhead | 0.083 | 2.381 | 1.098 |
| 117 | 16-1044 | ACS_EGF | 0.512 | 1.592 | 0.460 |
| 118 | 16-181 | SMBSC | 40.302 | 2.073 | 0.761 |
| 119 | 16-186 | SMBSC | 0.091 | 1.713 | 0.842 |
| 120 | 16-224 | Minn-Dak_MN | 65.277 | 1.711 | 0.853 |
| 121 | 16-270 | Minn-Dak_MN | 65.26 | 1.927 | 0.741 |
| 122 | 16-29 | SMBSC | 87.739 | 2.169 | 0.946 |
| 123 | 16-326 | ACS_Drayton | 0.098 | 1.567 | 0.945 |

| | | | | | |
|---|---|---|---|---|---|
| 124 | 16-351 | ACS_Drayton | 57.213 | 1.842 | 0.630 |
| 125 | 16-36 | SMBSC | 0.081 | 1.235 | 0.606 |
| 126 | 16-363 | ACS_Drayton | 0.082 | 1.592 | 0.355 |
| 127 | 16-374 | ACS_Drayton | 55.56 | 1.971 | 0.655 |
| 128 | 16-412 | SMBSC | 48.483 | 1.319 | 0.667 |
| 129 | 16-438 | ACS_Crookston | 0.088 | 1.377 | 0.523 |
| 130 | 16-451 | ACS_Crookston | 65.927 | 2.039 | 0.938 |
| 131 | 16-503 | ACS_Hillsboro | 0.092 | 1.880 | 0.728 |
| 132 | 16-54 | SMBSC | 74.127 | 1.929 | 0.781 |
| 133 | 16-546 | ACS_Hillsboro | 18.033 | 1.354 | 0.883 |
| 134 | 16-569 | ACS_EGF | 81.667 | 2.112 | 0.816 |
| 135 | 16-638 | ACS_EGF | 0.075 | 1.463 | 0.637 |
| 136 | 16-723 | Minn-Dak_MN | 73.782 | 2.048 | 0.863 |
| 137 | 16-731 | Minn-Dak_MN | 48.907 | 1.976 | 0.903 |
| 138 | 16-795 | ACS_Moorhead | 11.942 | 2.088 | 0.574 |
| 139 | 16-827 | ACS_Moorhead | 0.081 | 2.111 | 0.245 |
| 140 | 16-842 | ACS_Crookston | 79.556 | 1.437 | 0.818 |
| 141 | 16-879 | SMBSC | 70.707 | 2.129 | 0.459 |
| 142 | 16-90 | SMBSC | 78.959 | 0.967 | 0.744 |
| 143 | 16-938 | ACS_Drayton | 72.796 | 1.757 | 0.694 |
| 144 | 16-946 | ACS_Drayton | 57.830 | 1.677 | 1.130 |
| 145 | 16-987 | ACS_Drayton | 0.098 | 1.649 | 0.311 |
| 146 | 17-1015 | SMBSC | 75.57 | 1.422 | 0.309 |
| 147 | 17-1045 | ACS_EGF | 98.66 | 1.842 | 0.635 |
| 148 | 17-1047 | ACS_EGF | 0.034 | 1.976 | 0.332 |
| 149 | 17-1070 | ACS_Crookston | 0.061 | 1.804 | 0.509 |
| 150 | 17-1096 | ACS_Drayton | 0.056 | 1.513 | 0.547 |
| 151 | 17-1109 | ACS_Hillsboro | 0.058 | 2.076 | 0.432 |
| 152 | 17-1121 | ACS_Crookston | 68.88 | 1.905 | 1.021 |
| 153 | 17-1127 | ACS_Drayton | 0.062 | 1.576 | 0.634 |
| 154 | 17-1132 | ACS_Drayton | 58.81 | 2.307 | 0.483 |

| | | | | | |
|---|---|---|---|---|---|
| 155 | 17-1162 | SMBSC | 76.20 | 1.890 | 0.320 |
| 156 | 17-132 | Minn-Dak_MN | 0.072 | 1.614 | 0.550 |
| 157 | 17-177 | ACS_Drayton | 0.091 | 1.819 | 0.553 |
| 158 | 17-214 | ACS_Drayton | 0.008 | 2.125 | 0.481 |
| 159 | 17-235 | ACS_Drayton | 0.008 | 1.800 | 0.529 |
| 160 | 17-251 | ACS_Drayton | 0.062 | 1.493 | 0.556 |
| 161 | 17-300 | Minn-Dak_MN | 38.781 | 2.309 | 0.705 |
| 162 | 17-307 | Minn-Dak_MN | 0.077 | 2.282 | 0.614 |
| 163 | 17-354 | SMBSC | 65.364 | 1.733 | 0.541 |
| 164 | 17-363 | SMBSC | 0.063 | 1.605 | 0.572 |
| 165 | 17-385 | ACS_Moorhead | 0.093 | 1.179 | 0.422 |
| 166 | 17-390 | SMBSC | 72.840 | 1.260 | 0.787 |
| 167 | 17-410 | SMBSC | 0.083 | 1.467 | 0.503 |
| 168 | 17-443 | SMBSC | 0.088 | 1.543 | 0.557 |
| 169 | 17-478 | ACS_EGF | 0.06 | 1.674 | 0.459 |
| 170 | 17-48 | ID | 81.087 | 1.411 | 0.732 |
| 171 | 17-51 | ID | 0.082 | 1.522 | 1.106 |
| 172 | 17-516 | Minn-Dak_MN | 73.124 | 1.774 | 0.494 |
| 173 | 17-528 | Minn-Dak_MN | 0.077 | 1.909 | 0.471 |
| 174 | 17-585 | Minn-Dak_MN | 0.08 | 2.346 | 0.705 |
| 175 | 17-594 | Minn-Dak_MN | 50.235 | 2.046 | 0.807 |
| 176 | 17-633 | ACS_Hillsboro | 59.885 | 2.039 | 0.772 |
| 177 | 17-661 | ACS_Hillsboro | 77.500 | 1.852 | 0.685 |
| 178 | 17-696 | ACS_Crookston | 0.008 | 1.750 | 0.655 |
| 179 | 17-724 | ACS_EGF | 35.714 | 2.205 | 0.548 |
| 180 | 17-741 | ACS_EGF | 0.008 | 1.692 | 0.570 |
| 181 | 17-760 | ACS_Hillsboro | 54.563 | 1.577 | 0.526 |
| 182 | 17-799 | ACS_Drayton | 0.07 | 1.809 | 0.465 |
| 183 | 17-82 | SMBSC | 58.507 | 1.813 | 0.458 |
| 184 | 17-837 | ACS_Hillsboro | 100.000 | 1.813 | 0.904 |
| 185 | 17-852 | ACS_Hillsboro | 70.898 | 1.997 | 0.805 |

| | | | | | |
|---|---|---|---|---|---|
| 186 | 17-869 | ACS_Hillsboro | 0.096 | 2.154 | 0.590 |
| 187 | 17-912 | ACS_EGF | 46.362 | 2.187 | 0.975 |
| 188 | 17-933 | ACS_EGF | 0.51 | 1.703 | 0.547 |
| 189 | 16-1170 | ACS_Drayton | 0.076 | 1.971 | 0.241 |
| 190 | 16-892 | SMBSC | 0.084 | 1.736 | 0.462 |
| 191 | 17-1087 | ACS_Drayton | 52.75 | 1.972 | 0.435 |
| 192 | 17-256 | ACS_Drayton | 0.070 | 1.555 | 0.946 |
| 193 | 17-432 | SMBSC | 19.501 | 1.340 | 0.544 |
| 194 | 17-501 | Minn-Dak_MN | 58.843 | 1.430 | 0.662 |
| 195 | 17-950 | ACS_EGF | 83.290 | 2.123 | 0.935 |

[a] Numbers used to refer to individual isolates in PCA plots and other analysis performed in this study.

[b] Isolate names used for NCBI short read archive. 16/17 refers to the year of isolate collection (2016/17).

[c] Indicates factory district from which isolate was collected. ACS = American Crystal Sugar. BASF = BASF chemicals company. EGF = East Grand Forks. Minn-Dak = Minn-Dak Farmers Cooperative. MN = Minnesota. SMBSC = Southern Minnesota Beet Sugar Cooperative.

[d] Effective concentration to reduce growth by 50% ($EC_{50}$) values were calculated for tetraconazole as described by Secor and Rivera (2012).

[e] Radial growth rate for each isolate in mm/day was calculated by growing them on CV8 plates and using linear regression for mean measurements taken at 2, 6, 9, 13 and 16 days.

[f] Radial growth for each isolate under salt stress in mm/day was calculated by growing them on CV8 plates amended with 1M NaCl and using linear regression for mean measurements taken at 6, 9, 13, 16, 20 and 23 days.

**Table S2. Genome sequencing statistics.**

| Isolate Name | Biosample ID[a] | Total number of reads | Sequence length (bp) | Reference genome alignment rate (%) | Mean reference genome coverage (X) |
|---|---|---|---|---|---|
| 16-F1 | SAMN16625845 | 12718465 | 100 | 95.87 | 33 |
| 16-F2 | SAMN16625846 | 13166029 | 100 | 95.47 | 34 |
| 16-F3 | SAMN16625847 | 10804832 | 100 | 94.42 | 28 |
| 16-F4 | SAMN16625848 | 12773284 | 100 | 96.6 | 33 |
| 16-F5 | SAMN16625849 | 9502166 | 100 | 96.18 | 25 |
| 16-F6 | SAMN16625850 | 11783600 | 100 | 96.62 | 31 |
| 16-F7 | SAMN16625851 | 11936524 | 100 | 96.35 | 31 |
| 16-F9 | SAMN16625852 | 12795308 | 100 | 96.01 | 33 |
| 16-F11 | SAMN16625853 | 11072823 | 100 | 95.66 | 29 |
| 16-F12 | SAMN16625854 | 11310699 | 100 | 95.88 | 29 |
| 16-F13 | SAMN16625855 | 8903227 | 100 | 96.8 | 23 |
| 16-F14 | SAMN16625856 | 11152696 | 100 | 96.66 | 29 |
| 16-F15 | SAMN16625857 | 9859917 | 100 | 96.66 | 26 |
| 16-F16 | SAMN16625858 | 14266731 | 100 | 96.76 | 37 |
| 16-F17 | SAMN16625859 | 10947158 | 100 | 96.2 | 28 |
| 16-F18 | SAMN16625860 | 10098019 | 100 | 96.41 | 26 |
| 16-F19 | SAMN16625861 | 12044026 | 100 | 91.03 | 30 |
| 16-F20 | SAMN16625862 | 11250660 | 100 | 96.44 | 29 |
| 16-F21 | SAMN16625863 | 11589045 | 100 | 96.08 | 30 |
| 16-F22 | SAMN16625864 | 12556783 | 100 | 96.09 | 33 |
| 16-F23 | SAMN16625865 | 10470666 | 100 | 97.59 | 28 |
| 16-F24 | SAMN16625866 | 8987040 | 100 | 97.04 | 24 |
| 16-F25 | SAMN16625867 | 11928975 | 100 | 96.69 | 31 |
| 16-F26 | SAMN16625868 | 10870741 | 100 | 96.64 | 28 |
| 16-F27 | SAMN16625869 | 11647159 | 100 | 96.38 | 30 |
| 16-F28 | SAMN16625870 | 11051769 | 100 | 95.63 | 29 |
| 16-F29 | SAMN16625871 | 12977620 | 100 | 95.68 | 34 |

| | | | | | |
|---|---|---|---|---|---|
| 16-F30 | SAMN16625872 | 11117965 | 100 | 96.37 | 29 |
| 16-F31 | SAMN16625873 | 12540975 | 100 | 95.32 | 32 |
| 16-F32 | SAMN16625874 | 14742865 | 100 | 96.24 | 38 |
| 16-F33 | SAMN16625875 | 13308868 | 100 | 95.06 | 34 |
| 16-F34 | SAMN16625876 | 14109197 | 100 | 96.54 | 37 |
| 16-F35 | SAMN16625877 | 13761148 | 100 | 95.91 | 36 |
| 16-F36 | SAMN16625878 | 11071775 | 100 | 96.74 | 29 |
| 16-F37 | SAMN16625879 | 12359187 | 100 | 97.07 | 32 |
| 16-F38 | SAMN16625880 | 11481667 | 100 | 96.57 | 30 |
| 16-F39 | SAMN16625881 | 12845661 | 100 | 96.18 | 33 |
| 16-F40 | SAMN16625882 | 10745460 | 100 | 98.09 | 28 |
| 16-F41 | SAMN16625883 | 10921792 | 100 | 97.72 | 29 |
| 16-F42 | SAMN16625884 | 12209843 | 100 | 96.34 | 32 |
| 16-F43 | SAMN16625885 | 13446151 | 100 | 96.2 | 35 |
| 16-F44 | SAMN16625886 | 12746723 | 100 | 97.21 | 33 |
| 16-F45 | SAMN16625887 | 10878158 | 100 | 96.03 | 28 |
| 16-F46 | SAMN16625888 | 12654252 | 100 | 97.06 | 33 |
| 16-F47 | SAMN16625889 | 11235283 | 100 | 96.2 | 29 |
| 16-F48 | SAMN16625890 | 9994376 | 100 | 96.07 | 26 |
| 16-F49 | SAMN16625891 | 11624916 | 100 | 96.49 | 30 |
| 16-F50 | SAMN16625892 | 12212953 | 100 | 97.35 | 32 |
| 16-F51 | SAMN16625893 | 12106448 | 100 | 96.02 | 31 |
| 16-F52 | SAMN16625894 | 11021092 | 100 | 97.42 | 29 |
| 16-F53 | SAMN16625895 | 12586280 | 100 | 95.94 | 33 |
| 16-F54 | SAMN16625896 | 10207809 | 100 | 96 | 26 |
| 16-F55 | SAMN16625897 | 9733604 | 100 | 96.6 | 25 |
| 16-F56 | SAMN16625898 | 12880745 | 100 | 97.14 | 34 |
| 16-F57 | SAMN16625899 | 13340824 | 100 | 97.4 | 35 |
| 16-F58 | SAMN16625900 | 10964146 | 100 | 95.33 | 28 |
| 16-F59 | SAMN16625901 | 11398428 | 100 | 95.98 | 30 |
| 16-16 | SAMN16625902 | 15393529 | 100 | 95.45 | 40 |

| | | | | | |
|---|---|---|---|---|---|
| 16-45 | SAMN16625903 | 15338757 | 100 | 95.17 | 39 |
| 16-46 | SAMN16625904 | 15337957 | 100 | 94.76 | 39 |
| 16-62 | SAMN16625905 | 15248588 | 100 | 94.95 | 39 |
| 16-92 | SAMN16625906 | 15075186 | 100 | 90.35 | 37 |
| 16-131 | SAMN16625907 | 15179611 | 100 | 94.25 | 39 |
| 16-175 | SAMN16625908 | 13872122 | 100 | 95.74 | 36 |
| 16-339 | SAMN16625909 | 12810204 | 100 | 95.31 | 33 |
| 16-591 | SAMN16625910 | 10930000 | 100 | 95.98 | 28 |
| 16-665 | SAMN16625911 | 12737134 | 100 | 95.95 | 33 |
| 16-735 | SAMN16625912 | 12459059 | 100 | 96.02 | 32 |
| 16-781 | SAMN16625913 | 10895348 | 100 | 95.99 | 28 |
| 16-787 | SAMN16625914 | 10523070 | 100 | 96.08 | 27 |
| 16-805 | SAMN16625915 | 12611611 | 100 | 95.92 | 33 |
| 16-810 | SAMN16625916 | 9525664 | 100 | 95.79 | 25 |
| 16-890 | SAMN16625917 | 11498111 | 100 | 96.39 | 30 |
| 16-1148 | SAMN16625918 | 10304038 | 100 | 95.94 | 27 |
| 16-26 | SAMN16625919 | 13359646 | 100 | 95.09 | 34 |
| 16-33 | SAMN16625920 | 13508108 | 100 | 95.67 | 35 |
| 16-72 | SAMN16625921 | 13793982 | 100 | 94.15 | 35 |
| 16-93 | SAMN16625922 | 13455330 | 100 | 93.24 | 34 |
| 16-317 | SAMN16625923 | 13515885 | 100 | 95.53 | 35 |
| 16-414 | SAMN16625924 | 13338997 | 100 | 94.32 | 34 |
| 16-498 | SAMN16625925 | 13530538 | 100 | 94.87 | 35 |
| 16-523 | SAMN16625926 | 13478816 | 100 | 95.15 | 35 |
| 16-666 | SAMN16625927 | 13418499 | 100 | 95.07 | 34 |
| 16-704 | SAMN16625928 | 13727960 | 100 | 94.66 | 35 |
| 16-716 | SAMN16625929 | 15463661 | 100 | 92.43 | 39 |
| 16-722 | SAMN16625930 | 15239636 | 100 | 95.86 | 39 |
| 16-724 | SAMN16625931 | 15323018 | 100 | 95.85 | 40 |
| 16-790 | SAMN16625932 | 15338450 | 100 | 96.02 | 40 |
| 16-819 | SAMN16625933 | 15289810 | 100 | 96.14 | 40 |

| | | | | | |
|---|---|---|---|---|---|
| 16-868 | SAMN16625934 | 15338708 | 100 | 95.81 | 40 |
| 16-1105 | SAMN16625935 | 15381554 | 100 | 95.88 | 40 |
| 16-1169 | SAMN16625936 | 15320524 | 100 | 95.25 | 39 |
| 16-F61 | SAMN16625937 | 8889309 | 150 | 96.35 | 35 |
| 16-F62 | SAMN16625938 | 8931302 | 150 | 96.21 | 35 |
| 16-F63 | SAMN16625939 | 8922341 | 150 | 96.72 | 35 |
| 16-F64 | SAMN16625940 | 8929826 | 150 | 96.04 | 35 |
| 16-F65 | SAMN16625941 | 8170519 | 150 | 95.83 | 32 |
| 16-100 | SAMN16625942 | 8154726 | 150 | 96.03 | 32 |
| 16-1022 | SAMN16625943 | 8387188 | 150 | 95.48 | 32 |
| 16-1083 | SAMN16625944 | 8181686 | 150 | 94.99 | 31 |
| 16-1110 | SAMN16625945 | 8322394 | 150 | 96.25 | 32 |
| 16-1124 | SAMN16625946 | 8213048 | 150 | 95.8 | 32 |
| 16-1129 | SAMN16625947 | 8228243 | 150 | 96.2 | 32 |
| 16-1138 | SAMN16625948 | 8229474 | 150 | 95.84 | 32 |
| 16-1151 | SAMN16625949 | 8021108 | 150 | 95.8 | 31 |
| 16-1155 | SAMN16625950 | 8249103 | 150 | 96.73 | 32 |
| 16-1211 | SAMN16625951 | 8204928 | 150 | 96.05 | 32 |
| 16-1214 | SAMN16625952 | 8233643 | 150 | 95.79 | 32 |
| 16-123 | SAMN16625953 | 8141259 | 150 | 96 | 32 |
| 16-141 | SAMN16625954 | 8217388 | 150 | 96.3 | 32 |
| 16-163 | SAMN16625955 | 8113850 | 150 | 96.36 | 32 |
| 16-1044 | SAMN16625956 | 7390194 | 150 | 95.91 | 29 |
| 16-181 | SAMN16625957 | 8203241 | 150 | 96.45 | 32 |
| 16-186 | SAMN16625958 | 8190961 | 150 | 96.11 | 32 |
| 16-224 | SAMN16625959 | 7214579 | 150 | 95.53 | 28 |
| 16-270 | SAMN16625960 | 8360024 | 150 | 96.19 | 33 |
| 16-29 | SAMN16625961 | 8038226 | 150 | 96.29 | 31 |
| 16-326 | SAMN16625962 | 8025845 | 150 | 96.07 | 31 |
| 16-351 | SAMN16625963 | 8064410 | 150 | 96.25 | 31 |
| 16-36 | SAMN16625964 | 8079058 | 150 | 96.04 | 31 |

| | | | | | |
|---|---|---|---|---|---|
| 16-363 | SAMN16625965 | 8075125 | 150 | 96.38 | 32 |
| 16-374 | SAMN16625966 | 7468268 | 150 | 94.99 | 29 |
| 16-412 | SAMN16625967 | 8153998 | 150 | 96.12 | 32 |
| 16-438 | SAMN16625968 | 8150620 | 150 | 96.01 | 32 |
| 16-451 | SAMN16625969 | 8233113 | 150 | 96.25 | 32 |
| 16-503 | SAMN16625970 | 8181538 | 150 | 95.36 | 32 |
| 16-54 | SAMN16625971 | 8047244 | 150 | 95.97 | 31 |
| 16-546 | SAMN16625972 | 8567009 | 150 | 62.02 | 22 |
| 16-569 | SAMN16625973 | 8223639 | 150 | 95.99 | 32 |
| 16-638 | SAMN16625974 | 8242271 | 150 | 95.25 | 32 |
| 16-723 | SAMN16625975 | 8288548 | 150 | 96.3 | 32 |
| 16-731 | SAMN16625976 | 8181859 | 150 | 96 | 32 |
| 16-795 | SAMN16625977 | 8252996 | 150 | 96.61 | 32 |
| 16-827 | SAMN16625978 | 8230298 | 150 | 96.33 | 32 |
| 16-842 | SAMN16625979 | 8269336 | 150 | 96.48 | 32 |
| 16-879 | SAMN16625980 | 8239261 | 150 | 97.09 | 32 |
| 16-90 | SAMN16625981 | 8233696 | 150 | 96.57 | 32 |
| 16-938 | SAMN16625982 | 7329161 | 150 | 96 | 28 |
| 16-946 | SAMN16625983 | 8281553 | 150 | 96.83 | 32 |
| 16-987 | SAMN16625984 | 8325224 | 150 | 96.44 | 32 |
| 17-1015 | SAMN16625985 | 8224169 | 150 | 96.73 | 32 |
| 17-1045 | SAMN16625986 | 8214141 | 150 | 96.48 | 32 |
| 17-1047 | SAMN16625987 | 8166920 | 150 | 95.85 | 32 |
| 17-1070 | SAMN16625988 | 8247959 | 150 | 96.21 | 32 |
| 17-1096 | SAMN16625989 | 8671766 | 150 | 51.31 | 18 |
| 17-1109 | SAMN16625990 | 8255032 | 150 | 96.31 | 32 |
| 17-1121 | SAMN16625991 | 8183028 | 150 | 96.5 | 32 |
| 17-1127 | SAMN16625992 | 8240663 | 150 | 96.2 | 32 |
| 17-1132 | SAMN16625993 | 7646731 | 150 | 95.88 | 30 |
| 17-1162 | SAMN16625994 | 8272871 | 150 | 96.92 | 32 |
| 17-132 | SAMN16625995 | 8195993 | 150 | 96.39 | 32 |

| | | | | | |
|---|---|---|---|---|---|
| 17-177 | SAMN16625996 | 8146135 | 150 | 96.37 | 32 |
| 17-214 | SAMN16625997 | 8255259 | 150 | 96.84 | 32 |
| 17-235 | SAMN16625998 | 8300163 | 150 | 96.46 | 32 |
| 17-251 | SAMN16625999 | 8254369 | 150 | 96.46 | 32 |
| 17-300 | SAMN16626000 | 8213705 | 150 | 97 | 32 |
| 17-307 | SAMN16626001 | 8192075 | 150 | 96.58 | 32 |
| 17-354 | SAMN16626002 | 8186482 | 150 | 96.66 | 32 |
| 17-363 | SAMN16626003 | 8200985 | 150 | 96.58 | 32 |
| 17-385 | SAMN16626004 | 8165428 | 150 | 95.39 | 32 |
| 17-390 | SAMN16626005 | 8246570 | 150 | 96.42 | 32 |
| 17-410 | SAMN16626006 | 8235966 | 150 | 96.51 | 32 |
| 17-443 | SAMN16626007 | 8231728 | 150 | 96.62 | 32 |
| 17-478 | SAMN16626008 | 8223309 | 150 | 96.57 | 32 |
| 17-48 | SAMN16626009 | 8208937 | 150 | 96.65 | 32 |
| 17-51 | SAMN16626010 | 8291290 | 150 | 96.44 | 32 |
| 17-516 | SAMN16626011 | 8198392 | 150 | 96.74 | 32 |
| 17-528 | SAMN16626012 | 8169356 | 150 | 97.23 | 32 |
| 17-585 | SAMN16626013 | 8211420 | 150 | 96.59 | 32 |
| 17-594 | SAMN16626014 | 8250461 | 150 | 97.4 | 33 |
| 17-633 | SAMN16626015 | 8214609 | 150 | 96.88 | 32 |
| 17-661 | SAMN16626016 | 8375374 | 150 | 96.36 | 33 |
| 17-696 | SAMN16626017 | 8229868 | 150 | 96.61 | 32 |
| 17-724 | SAMN16626018 | 8320330 | 150 | 96.34 | 32 |
| 17-741 | SAMN16626019 | 8165723 | 150 | 96.58 | 32 |
| 17-760 | SAMN16626020 | 8307505 | 150 | 96.76 | 33 |
| 17-799 | SAMN16626021 | 8306128 | 150 | 96.4 | 32 |
| 17-82 | SAMN16626022 | 8153198 | 150 | 96.63 | 32 |
| 17-837 | SAMN16626023 | 8412166 | 150 | 96.97 | 33 |
| 17-852 | SAMN16626024 | 8252594 | 150 | 96.38 | 32 |
| 17-869 | SAMN16626025 | 8332658 | 150 | 96.75 | 33 |
| 17-912 | SAMN16626026 | 8363854 | 150 | 96.7 | 33 |

| | | | | | |
|---|---|---|---|---|---|
| 17-933 | SAMN16626027 | 8290847 | 150 | 96.86 | 33 |
| 16-1170 | SAMN16626028 | 8267245 | 150 | 96.54 | 32 |
| 16-892 | SAMN16626029 | 8184082 | 150 | 96.5 | 32 |
| 17-1087 | SAMN16626030 | 8215639 | 150 | 96.84 | 32 |
| 17-256 | SAMN16626031 | 8297680 | 150 | 96.04 | 32 |
| 17-432 | SAMN16626032 | 8236108 | 150 | 96.53 | 32 |
| 17-501 | SAMN16626033 | 8200429 | 150 | 96.78 | 32 |
| 17-950 | SAMN16626034 | 8291920 | 150 | 96.74 | 32 |

[a]Biosample accession number in the NCBI Short Read Archive, allowing raw reads to be freely downloaded.

**Table S3. Thirteen significant associations for GWAS of tetraconazole sensitivity at a significance threshold of $-\log_{10}(p) = 4.5$.**

| Chr[a] | Position (bp) | *P*-value | -log10(*P*) | Reference allele | Alternate allele | Gene ID | Gene annotation | Mutation | MAF[b] |
|---|---|---|---|---|---|---|---|---|---|
| 1 | 1890198 | 7.04E-06 | 5.1524 | C | - | CB0940_00689 | Sorting nexin-12 | 5'UTR 76 bp indel | 0.27 |
| 1 | 2637787 | 9.41E-06 | 5.0264 | A | G | CB0940_00999 | Hypothetical protein | S474G | 0.47 |
| 9 | 1503309 | 1.56E-05 | 4.8069 | C | G | CB0940_11399 | ABC transporter CDR4 | Synonymous (L191) | 0.47 |
| 9 | 1452111 | 1.83E-05 | 4.7375 | A | G | CB0940_11379 | *CbCYP51* | 124 bp upstream of ATG | 0.47 |
| 9 | 1503297 | 1.84E-05 | 4.7352 | A | C | CB0940_11399 | ABC transporter CDR4 | Synonymous (L195) | 0.47 |
| 9 | 1503321 | 1.95E-05 | 4.7100 | A | T | CB0940_11399 | ABC transporter CDR4 | Synonymous (G187) | 0.47 |
| 9 | 1503303 | 1.95E-05 | 4.7100 | T | C | CB0940_11399 | ABC transporter CDR4 | Synonymous (K193) | 0.46 |
| 9 | 1503315 | 1.96E-05 | 4.7077 | G | A | CB0940_11399 | ABC transporter CDR4 | Synonymous (S189) | 0.47 |
| 9 | 1503312 | 1.96E-05 | 4.7077 | C | G | CB0940_11399 | ABC transporter CDR4 | Synonymous (T190) | 0.47 |
| 9 | 1502965 | 2.21E-05 | 4.6556 | T | G | CB0940_11399 | ABC transporter CDR4 | Intron | 0.48 |
| 4 | 3725552 | 2.59E-05 | 4.5867 | A | - | CB0940_05141 | Dual specificity protein kinase Pom1 | 5'UTR 63 bp indel | 0.23 |
| 9 | 1502968 | 2.88E-05 | 4.5406 | G | C | CB0940_11399 | ABC transporter CDR4 | Intron | 0.47 |
| 9 | 1502966 | 2.88E-05 | 4.5406 | C | T | CB0940_11399 | ABC transporter CDR4 | Intron | 0.47 |

[a]Chromosome/scaffold number

[b]Minor allele frequency

**Table S4. Predicted transcription factor binding sites for the 9_1452111 SNP upstream of *CbCYP51* using the JASPAR database.**

| Sequence ID[a] | Matrix ID[b] | TF Name[c] | Score[d] | Relative score[e] | Start (bp)[f] | End (bp)[f] | Strand[g] | Predicted sequence[h] |
|---|---|---|---|---|---|---|---|---|
| Sensitive_C | MA0393.1 | STE12 | 10.00 | 0.96 | 6 | 12 | + | tgaaacc |
| Sensitive_C | MA0411.1 | UPC2 | 8.70 | 0.94 | 12 | 18 | + | cgtacga |
| Sensitive_C | MA0411.1 | UPC2 | 5.80 | 0.84 | 11 | 17 | - | cgtacgg |
| Sensitive_C | MA0292.1 | ECM22 | 5.98 | 0.84 | 9 | 15 | + | aaccgta |
| Sensitive_C | MA0392.1 | STB5 | 6.32 | 0.84 | 6 | 13 | - | cggtttca |
| Resistant_T | MA0411.1 | UPC2 | 8.70 | 0.94 | 11 | 17 | - | cgtacga |
| Resistant_T | MA0411.1 | UPC2 | 8.70 | 0.94 | 12 | 18 | + | cgtacga |
| Resistant_T | MA0307.1 | GLN3 | 4.82 | 0.86 | 8 | 12 | - | gattt |
| Resistant_T | MA0393.1 | STE12 | 2.35 | 0.83 | 6 | 12 | + | tgaaatc |
| Resistant_T | MA0388.1 | SPT23 | 5.00 | 0.82 | 5 | 12 | - | gatttcaa |
| Resistant_T | MA0388.1 | SPT23 | 4.20 | 0.80 | 7 | 14 | + | gaaatcgt |
| Resistant_T | MA0388.1 | SPT23 | 4.11 | 0.80 | 8 | 15 | + | aaatcgta |

[a] Sequences of 21 bp in length were used as queries with the 11$^{th}$ bp as the sensitive C allele or the resistant T allele.

[b]Position frequency matrices in JASPAR database representing DNA-binding sites determined experimentally for a transcription factor and used to scan query sequences.

[c]Name of transcription factor in *Saccharomyces cerevisiae*.

[d]Raw score for binding site.

[e]Relative score is the fraction of the potential maximum score for the binding site.

[f]The predicted start and end base pairs of the binding site within the query sequence.

[g]Binding site is present on either the positive (forward) or negative (reverse) strand of DNA.

[h]Predicted binding site within query sequence.

**Table S5. Codon usage for the *C. beticola* 09-40 reference genome.**

| Amino acid | Codon | Total number | Proportion of amino acid codon usage (%) |
|---|---|---|---|
| Ala | GCG | 118196 | 22 |
| Ala | GCA | 145227 | 27 |
| Ala | GCT | 140842 | 26 |
| Ala | GCC | 142178 | 26 |
| Cys | TGT | 33364 | 37 |
| Cys | TGC | 56751 | 63 |
| Asp | GAT | 161658 | 47 |
| Asp | GAC | 179197 | 53 |
| Glu | GAG | 209892 | 56 |
| Glu | GAA | 162472 | 44 |
| Phe | TTT | 62313 | 30 |
| Phe | TTC | 147005 | 70 |
| Gly | GGG | 53396 | 13 |
| Gly | GGA | 111288 | 26 |
| Gly | GGT | 90404 | 21 |
| Gly | GGC | 166563 | 40 |
| His | CAT | 72845 | 46 |
| His | CAC | 87140 | 54 |
| Ile | ATA | 35549 | 13 |
| Ile | ATT | 86957 | 32 |
| Ile | ATC | 150516 | 55 |
| Lys | AAG | 187064 | 66 |
| Lys | AAA | 98367 | 34 |
| Leu | TTG | 97005 | 19 |
| Leu | TTA | 20659 | 4 |
| Leu | CTG | 126837 | 25 |
| Leu | CTA | 41126 | 8 |
| Leu | CTT | 85003 | 17 |
| Leu | CTC | 141355 | 28 |
| Met | ATG | 123264 | 100 |
| Asn | AAT | 89810 | 42 |
| Asn | AAC | 126518 | 58 |
| Pro | CCG | 81410 | 23 |
| Pro | CCA | 112773 | 32 |
| Pro | CCT | 91854 | 26 |
| Pro | CCC | 70565 | 20 |
| Gln | CAG | 140058 | 52 |
| Gln | CAA | 128544 | 48 |
| Arg | AGG | 48644 | 12 |

| | | | |
|---|---|---|---|
| Arg | AGA | 65561 | 16 |
| Arg | CGG | 47562 | 12 |
| Arg | CGA | 96019 | 23 |
| Arg | CGT | 56083 | 14 |
| Arg | CGC | 98064 | 24 |
| Ser | AGT | 62408 | 13 |
| Ser | AGC | 111131 | 23 |
| Ser | TCG | 87936 | 18 |
| Ser | TCA | 70090 | 14 |
| Ser | TCT | 76449 | 16 |
| Ser | TCC | 81223 | 17 |
| Thr | ACG | 86442 | 24 |
| Thr | ACA | 94404 | 26 |
| Thr | ACT | 83744 | 23 |
| Thr | ACC | 96140 | 27 |
| Val | GTG | 106502 | 30 |
| Val | GTA | 41238 | 12 |
| Val | GTT | 70894 | 20 |
| Val | GTC | 132150 | 38 |
| Trp | TGG | 92885 | 100 |
| Tyr | TAT | 61128 | 38 |
| Tyr | TAC | 101542 | 62 |
| End | TGA | 18264 | 62 |
| End | TAG | 5995 | 20 |
| End | TAA | 5048 | 17 |

**Table S6. Demographic models were compared based on their Akaike information criterion (AIC) score.** The observed data showed the best fit to the model with the lowest AIC score.

| | Model | | | | |
|---|---|---|---|---|---|
| | Bottleneck | Expansion | Bottleneck-bottleneck | Bottleneck-Expansion | Observed |
| Max log-likelihood | -48,637,770 | - 122,648.287 | - 48,640.631 | - 48,712.283 | -48,448.960 |
| AIC | 188.81 | 74,199.327 | 191,671 | 263,323 | -- |

**Table S11. Difference between the expected and the observed maximum likelihood with different mutation rates.**

| Mutation Rate | $5 \times 10^{-7}$ | $5 \times 10^{-8}$ | $3 \times 10^{-8}$ | $1 \times 10^{-8}$ |
|---|---|---|---|---|
| Expected - observed maximum likelihood | 640,638 | 216.939 | 203.997 | 195.318 |

**Table S12. Genomic features within the selective sweep region on chromosome 4 that overlaps with the GWAS DMI resistance DYRK gene candidate (CB0940_05141).**

| Chromosome | Gene ID | Annotation | Feature | Start | End | Strand |
|---|---|---|---|---|---|---|
| CM008502.1 | CB0940_05140 | Hypothetical protein | exon | 3724388 | 3724629 | - |
| CM008502.1 | | | CDS | 3724388 | 3724629 | - |
| CM008502.1 | | | exon | 3724298 | 3724326 | - |
| CM008502.1 | | | CDS | 3724298 | 3724326 | - |
| CM008502.1 | | | CDS | 3724046 | 3724218 | - |
| CM008502.1 | CB0940_05141 | Dual specificity protein kinase pom1 | CDS | 3725740 | 3725866 | + |
| CM008502.1 | | | exon | 3725915 | 3726213 | + |
| CM008502.1 | | | CDS | 3725915 | 3726213 | + |
| CM008502.1 | | | exon | 3726268 | 3726427 | + |
| CM008502.1 | | | CDS | 3726268 | 3726427 | + |
| CM008502.1 | | | exon | 3726489 | 3727396 | + |
| CM008502.1 | | | CDS | 3726489 | 3726694 | + |
| CM008502.1 | CB0940_05142 | Hypothetical protein | exon | 3729922 | 3730137 | - |
| CM008502.1 | | | CDS | 3729922 | 3730137 | - |
| CM008502.1 | | | exon | 3729722 | 3729850 | - |
| CM008502.1 | | | CDS | 3729722 | 3729850 | - |
| CM008502.1 | | | exon | 3729450 | 3729675 | - |
| CM008502.1 | | | CDS | 3729450 | 3729675 | - |
| CM008502.1 | | | exon | 3727783 | 3729398 | - |
| CM008502.1 | | | CDS | 3727783 | 3729398 | - |
| CM008502.1 | | | gene | 3727441 | 3730137 | - |
| CM008502.1 | | | mRNA | 3727441 | 3730137 | - |
| CM008502.1 | | | exon | 3727441 | 3727719 | - |
| CM008502.1 | | | CDS | 3727441 | 3727719 | - |
| CM008502.1 | CB0940_05143 | Hypothetical protein | CDS | 3733971 | 3734059 | + |
| CM008502.1 | | | exon | 3734117 | 3734179 | + |
| CM008502.1 | | | CDS | 3734117 | 3734179 | + |
| CM008502.1 | | | exon | 3734235 | 3734719 | + |
| CM008502.1 | | | CDS | 3734235 | 3734719 | + |
| CM008502.1 | | | exon | 3734790 | 3735754 | + |
| CM008502.1 | | | CDS | 3734790 | 3735754 | + |
| CM008502.1 | | | exon | 3735813 | 3736463 | + |
| CM008502.1 | | | CDS | 3735813 | 3736292 | + |
| CM008502.1 | CB0940_05144 | Copper amine oxidase 1 | exon | 3740066 | 3740619 | - |
| CM008502.1 | | | CDS | 3740066 | 3740554 | - |
| CM008502.1 | | | gene | 3738441 | 3740619 | - |
| CM008502.1 | | | mRNA | 3738441 | 3740619 | - |

| | | | | |
|---|---|---|---|---|
| CM008502.1 | exon | 3738441 | 3740012 | - |
| CM008502.1 | CDS | 3738441 | 3740012 | - |

**Table S13. Genomic features within the selective sweep region on chromosome 9 that overlaps with the GWAS DMI resistance ABC PDR transporter gene candidate (CB0940_11399).**

| Chromosome | Gene ID | Annotation | Feature | Start | End | Strand |
|---|---|---|---|---|---|---|
| CM008507.1 | CB0940_11398 | Hypothetical protein | gene | 1496431 | 1497642 | - |
| CM008507.1 | | | mRNA | 1496431 | 1497642 | - |
| CM008507.1 | | | exon | 1496431 | 1497642 | - |
| CM008507.1 | | | CDS | 1496431 | 1497642 | - |
| CM008507.1 | CB0940_11399 | ABC transporter CDR4 | exon | 1503156 | 1504102 | - |
| CM008507.1 | | | CDS | 1503156 | 1503881 | - |
| CM008507.1 | | | exon | 1502986 | 1503095 | - |
| CM008507.1 | | | CDS | 1502986 | 1503095 | - |
| CM008507.1 | | | exon | 1502807 | 1502933 | - |
| CM008507.1 | | | CDS | 1502807 | 1502933 | - |
| CM008507.1 | | | exon | 1502149 | 1502751 | - |
| CM008507.1 | | | CDS | 1502149 | 1502751 | - |
| CM008507.1 | | | exon | 1500978 | 1502099 | - |
| CM008507.1 | | | CDS | 1500978 | 1502099 | - |
| CM008507.1 | | | exon | 1500599 | 1500918 | - |
| CM008507.1 | | | CDS | 1500599 | 1500918 | - |
| CM008507.1 | | | exon | 1499196 | 1500542 | - |
| CM008507.1 | | | CDS | 1499196 | 1500542 | - |
| CM008507.1 | | | CDS | 1499010 | 1499136 | - |

**Table S14. Allelic effect estimates for the most significantly associated markers with tetraconazole sensitivity.**

| Chromosome | Position (bp) | Associated gene annotation | Marker type[a] | Allelic effect on tetraconazole sensitivity ($EC_{50}$)[b] | Allelic effect on growth rate (mm/day)[b] | Allelic effect on growth rate under salt stress (mm/day)[b] |
|---|---|---|---|---|---|---|
| 1 | 1890198 | Sorting nexin-12 | Indel | -0.372 | 0.003 | -0.020 |
| 1 | 2637787 | Hypothetical protein | SNP | -0.615 | -0.014 | -0.023 |
| 9 | 1503309 | ABC transporter CDR4 | SNP | -0.541 | -0.030 | -0.006 |
| 9 | 1452111 | Upstream of *CbCYP51* | SNP | -0.591 | -0.039 | -0.034 |
| 9 | 1503297 | ABC transporter CDR4 | SNP | -0.540 | -0.030 | -0.007 |
| 9 | 1503321 | ABC transporter CDR4 | SNP | -0.540 | -0.030 | -0.008 |
| 9 | 1503303 | ABC transporter CDR4 | SNP | 0.540 | -0.030 | 0.009 |
| 9 | 1503315 | ABC transporter CDR4 | SNP | 0.540 | -0.030 | 0.007 |
| 9 | 1503312 | ABC transporter CDR4 | SNP | -0.540 | -0.030 | -0.007 |
| 9 | 1502965 | ABC transporter CDR4 | SNP | 0.525 | 0.029 | -0.003 |
| 4 | 3725552 | Dual specificity protein kinase Pom1 | Indel | -0.376 | -0.014 | 0.005 |
| 9 | 1502968 | ABC transporter CDR4 | SNP | 0.522 | 0.027 | -0.003 |
| 9 | 1502966 | ABC transporter CDR4 | SNP | -0.522 | 0.027 | 0.003 |

[a]Type of marker used in association analyses – an indel or single nucleotide polymorphism.

[b]Allelic effect estimates on phenotype derived from association analyses in GAPIT software.

**Table S15. Sequences of sgRNAs and donor template oligonucleotides used in Cas9-RNP editing.**

| Mutation | Attempt | SgRNA oligonucleotides[a] | Donor template[b] |
|---|---|---|---|
| E198A | 1 | TTCTAATACGACTCACTATAGACCTTCTGTATCGACAACGGTTTTAGAGCTAGA | GCCACTCTGTCCGTTCACCAGCTCGTCGAGAACTCCGACGC GACCTTCTGTATCGACAACGAGGCGCTGTACGACATTTGC |
| | 2 | TTCTAATACGACTCACTATAGTCGATACAGAAGGTCTCGTGTTTTAGAGCTAGA | |
| E170 | 1 | TTCTAATACGACTCACTATAGAAGAATTGGCGCGTTTCTTGTTTTAGAGCTAGA | CTGCTCTGCAGTCCTATGTCACATTGATCACCGAAGAAACG CGCCAATTCTTCTCTAAGAACAACCCACATAAGCGATTC |
| | 2 | TTCTAATACGACTCACTATAGGTTGTTCTTAGAGAAGAATGTTTTAGAGCTAGA | |
| | 3 | TTCTAATACGACTCACTATAGTCGAAGCGAATCGCTTATGGTTTTAGAGCTAGA | |
| L144F (C) | 1 | TTCTAATACGACTCACTATAGCTCCATGAATTTGGAGTTGGTTTTAGAGCTAGA | TTGGCAAGGACGTCGTCTACGACTGCCCCAACTCCAAATTC ATGGAGCAGAAGAAGTTTGTCAAGTTCGGCTTGACCAGCG |
| | 2 | TTCTAATACGACTCACTATAGTGGGGCAGTCGTAGACGACGGTTTTAGAGCTAGA | |
| | 3 | TTCTAATACGACTCACTATAGCTGCCCCAACTCCAAATTGAGTTTTAGAGCTAGA | |
| L144F (T) | 1 | TTCTAATACGACTCACTATAGCTCCATGAATTTGGAGTTGGTTTTAGAGCTAGA | TTGGCAAGGACGTCGTCTACGACTGCCCCAACTCCAAATTT ATGGAGCAGAAGAAGTTTGTCAAGTTCGGCTTGACCAGCG |
| | 2 | TTCTAATACGACTCACTATAGTGGGGCAGTCGTAGACGACGGTTTTAGAGCTAGA | |
| | 3 | TTCTAATACGACTCACTATAGCTGCCCCAACTCCAAATTGAGTTTTAGAGCTAGA | |
| Hygromycin resistance cassette insertion (intergenic) | 1 | TTCTAATACGACTCACTATAGTAAGCTTGTTAGTGTAATCAGTTTTAGAGCTAGA | PCR of pDAN with primers F: GTCGCGGGTTAAGCTTGTTAGTGTAATCAGGGTGCAACCC GACGTTGTAAAACGACGGCCAGTG and R: GGCACGCCAACGTTAGCATTGTTAAGCTCGAGTATAGCGC CACAGGAAACAGCTATGACCATGA |
| | 2 | TTCTAATACGACTCACTATAGACGGACTGCTCCTATCGCGGTTTTAGAGCTAGA | |

[a]Oligonucleotides used to synthesize single guide RNAs *in vitro*.

[b]Donor template is double-stranded and is produced by either complexing forward and reverse complement oligonucleotides (at room temperature) or amplifying a PCR product from template DNA (hygromycin resistance insertion cassette).
